# Stress adaptation pathways and HA–CD44 signaling maintain the survival of pancreatic cancer cells with centrosome amplification

**DOI:** 10.64898/2026.01.24.701523

**Authors:** Selahattin Can Ozcan, Evrim Goksel, Batuhan Mert Kalkan, Enes Cicek, Beste Kanevetci, Ceyda Acilan

## Abstract

Centrosome amplification (CA) is a hallmark of aggressive cancers, including pancreatic ductal adenocarcinoma (PDAC), and is linked to genomic instability and poor prognosis. While CA promotes tumor evolution, it also imposes substantial intracellular stress that cells must overcome to survive. However, the specific metabolic adaptations that enable cancer cells to tolerate stress induced by supernumerary centrosomes remain poorly understood. Here, we show that PDAC cells with CA acquire distinct metabolic dependencies that sustain survival. A metabolism-focused CRISPR-Cas9 screen, coupled with functional validations, identified critical vulnerabilities in three inter-connected axes: redox homeostasis, nucleotide sugar metabolism, and the unfolded protein response (UPR). Specifically, CA elevates intracellular reactive oxygen species (ROS), creating a reliance on glutamine metabolism and NRF2-driven antioxidant signaling. CRISPR screen hits in the hexosamine and uronic acid pathways revealed dependencies that converge on hyaluronic acid (HA) metabolism, and functional assays demonstrated that the HA–CD44 axis is required for centrosome clustering and mitotic fidelity, with its disruption increasing lethal multipolar divisions. In parallel, CA activated all branches of the UPR, and both hyper-activation and suppression of ER stress proved detrimental, indicating a finely tuned proteostatic equilibrium is essential for adaptation. Together, these findings show that, in a PLK4-driven model, centrosome-amplified cells rely on coordinated redox control, proteostatic buffering, and extracellular matrix signaling to tolerate CA-induced stress, revealing selective vulnerabilities that could be therapeutically exploited to target aggressive, therapy-resistant tumor subpopulations.

## 1 Introduction

Chromosomal instability (CIN) is a hallmark of cancer that drives tumor evolution while simultaneously imposing cellular stress that threatens cell viability^1^. One major source of CIN is centrosome amplification (CA), the presence of supernumerary centrosomes within a single cell. CA disrupts mitotic spindle organization, leading to merotelic attachments, chromosome mis-segregation, and aneuploidy^2,3^. To prevent lethal multipolar divisions, cancer cells rely on centrosome clustering mechanisms to restore the pseudo-bipolar spindle geometry, creating a structural vulnerability that has been proposed as a therapeutic target^4,5^.

CA is highly prevalent across cancers and particularly enriched in pancreatic ductal adenocarcinoma (PDAC), where it associates with advanced disease, metastasis, and poor patient survival^6–10^. Mechanistically, CA can arise from centriole overduplication (e.g., PLK4 or SAS-6 overexpression), loss of tumor suppressors such as p53 or BRCA1/2, or deregulated cell cycle progression^11,12^. Indeed, high expression of CA-associated genes including PLK4, STIL, and NEK2 predicts poor prognosis in PDAC patients^8^. These observations highlight CA as both a driver of aggressive tumor biology and a marker of lethal disease.

Beyond its direct impact on mitosis, CA triggers broader cellular stress adaptation programs. Centrosome-amplified cells exhibit a secretory phenotype characterized by increased release of cytokines and growth factors, which can reshape the tumor microenvironment^13,14^, and experience increased oxidative stress^14^. Supporting such outputs likely imposes proteotoxic and biosynthetic demands, while extra centrosomes themselves may create unique dependencies on cellular structures and signaling pathways beyond centrosome clustering. Consequently, CA likely necessitates a profound rewiring of cellular metabolism to fuel these adaptive responses and maintain survival. However, while the structural adaptations to CA, such as clustering, are well studied, the specific metabolic dependencies that enable cancer cells to tolerate the constant stress of supernumerary centrosomes remain a critical unanswered question.

PDAC represents a particularly relevant context in which to interrogate these adaptations. PDAC is defined by profound metabolic plasticity, including reliance on aerobic glycolysis, rewired glutamine metabolism, and scavenging of extracellular nutrients^15,16^. Moreover, PDAC cells must tolerate a hypoxic, nutrient-poor, and fibrotic microenvironment^17^, conditions that may exacerbate the stress imposed by CA. Yet, despite the prevalence of CA in PDAC and its association with poor prognosis, the metabolic requirements that allow cancer cells to tolerate supernumerary centrosomes remain poorly understood.

Here, we combined a doxycycline-inducible PLK4 model of CA with a metabolism-focused CRISPR-Cas9 screen in PDAC cells to systematically identify the survival pathways essential in the context of CA. This approach revealed that centrosome-amplified cells become critically dependent on specific pathways for redox homeostasis, nucleotide sugar metabolism, and UPR signaling. Furthermore, we discovered that the hyaluronic acid (HA)–CD44 axis is up-regulated and required for maintaining both centrosome clustering and cytokinesis fidelity. Our findings define a suite of targetable metabolic vulnerabilities that are essential for centrosome-amplified cell survival under CA-induced stress, revealing a new dimension of cancer cell addiction rooted in genomic instability and division disorders.

## 2 Results

### 2.1 Cells with centrosome amplification requires L-glutamine availability and GLS activity

We first utilized a doxycycline-inducible PLK4 over-expression system^18^ to explore the metabolic dependencies associated with CA in PDAC cells. After three days of doxycycline (dox) treatment, approximately 60% of Panc1-PLK4 and Mia Paca-2-PLK4 cells exhibited increased centrosome numbers (Fig. 1A, S1A). Upon extended duration of dox treatment, both cell lines maintained this increase in centrosome numbers for up to 20 days (Fig. 1B). While dox-induced CA reduced the proliferation rates in Panc1 cells over the course of three weeks, the proliferative decrease in Mia Paca-2 cells was partially rescued by week 3 (Fig. 1C). Additionally, we examined the cellular response to prolonged CA in U2OS cells, which harbor wild-type *p53*. Although the initial CA rate after 3 days was comparable to PDAC cell lines, U2OS cells did not sustain elevated centrosome numbers over time (Fig. S1B), a difference that could be linked to PIDDosome-mediated activation of the p53 pathway, which restricts the persistence of supernumerary centrosomes^19^. While cell proliferation was significantly affected by dox induction during the first week, this effect was diminished in the following two weeks (Fig. S1C). The ability of both PDAC cell lines, which carry mutant p53, to survive prolonged CA makes them good models for studying the long-term effects of CA in cancer.

**Figure 1.**
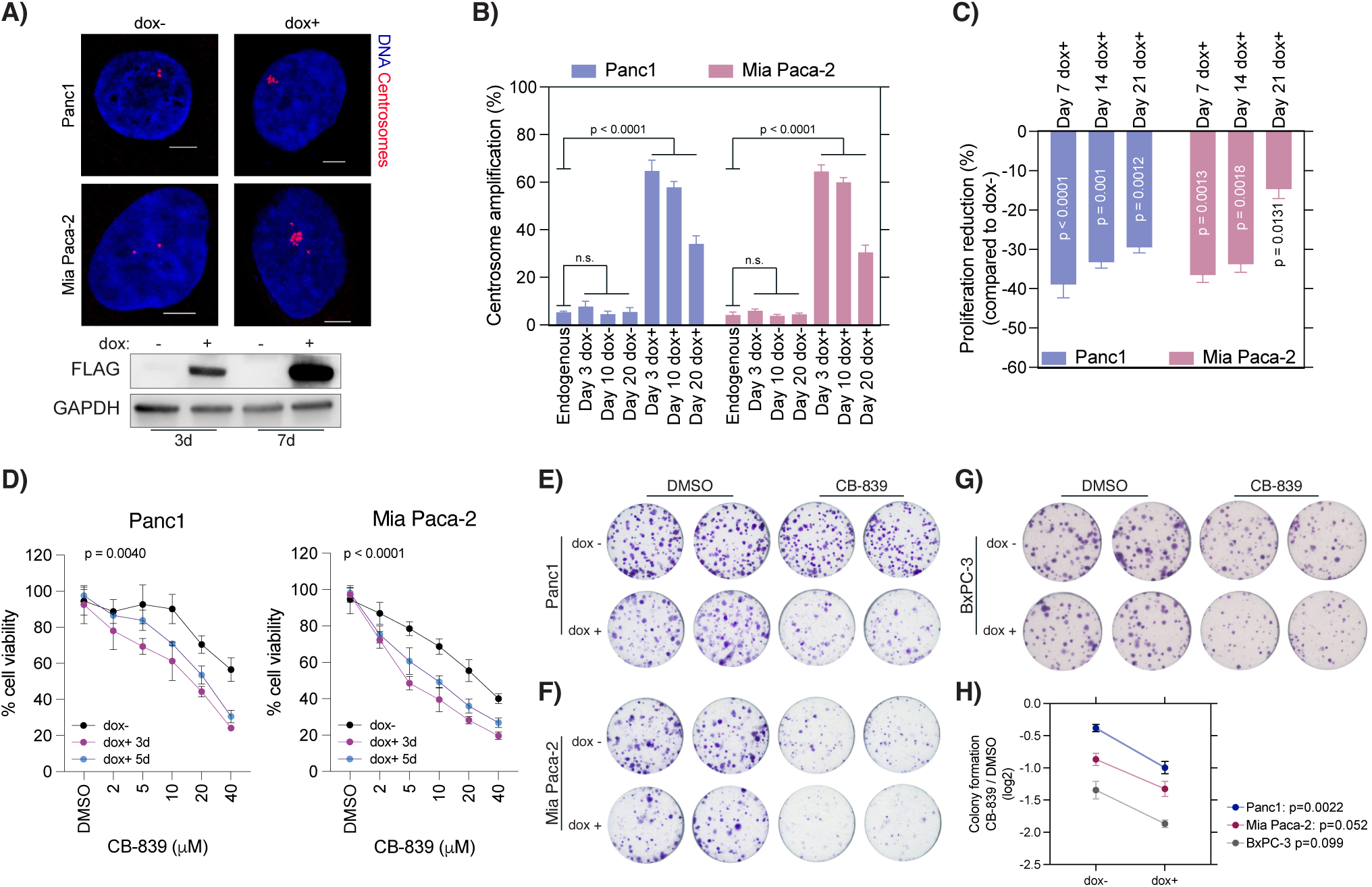
Centrosome amplification increases dependency on L-Glutamine metabolism in PDAC cells. A) Centrosome amplification in PDAC cell lines Panc1 and Mia Paca-2. Top panel: Confocal microscopy images. Blue: DAPI, nuclei; Red: γ-tubulin, centrosomes. Bottom panel: Induction of PLK4 expression by doxycycline. GAPDH was used as loading control. B) PDAC cells sustain high levels of CA over time. C) Persistent CA reduces cell proliferation rates in PDAC cells. D) PDAC cells with CA exhibit increased sensitivity to GLS1 inhibition by CB-839. Left panel: Panc1 cells, Right panel: Mia Paca-2 cells. E-H) CB-839 treatment significantly decreases the colony-formation ability of PDAC cells with CA. (E) Panc1 cells (F) Mia Paca-2 cells. (G) BxPC-3 cells. (H) Quantification results of colony formation experiments. Statistical significances were measured by two-way ANOVA in B and H, by two-tailed t-test on C, by non-linear curve fitting in D. p values were reported on graphs.

Since CA is linked to elevated intracellular reactive oxygen species (ROS) levels^14^, and glutamine metabolism plays a critical role in multiple ROS-eliminating pathways^20,21^, we hypothesized that cells with CA would exhibit increased vulnerability to disruptions in glutamine metabolism. Supporting this hypothesis, cells with CA exhibited increased lethality when treated with CB-839 (Telaglenastat), a glutaminase-1 (GLS1) inhibitor^22^ (Fig. 1D). Notably, both short-term (3d) and long-term (7d) centrosome-amplified cells demonstrated similar sensitivity to CB-839 treatment. Comparable results were observed in colony formation (CF) assays in both cell lines (Fig. 1E, 1F, 1H). Given that mutant KRAS rewires glutamine metabolism in a non-canonical cytoplasmic NADPH-synthesizing pathway in PDAC cells^23^, we extended this analysis to KRAS wild-type BxPC-3 cells. Similar to Panc1 and Mia Paca-2 cells, BxPC-3 cells with CA displayed increased sensitivity to CB-839 treatment (Fig. 1G, 1H), suggesting that the vulnerability is not attributable to the non-canonical glutamine function described. To rule out possible drug-specific off-target effects, we also tested another GLS1 inhibitor, BPTES, in Panc1-doxPLK4 cells and observed the reduction in colony formation assays as CB-839 (Fig. S1D). Finally, we evaluated the potential combined effects of doxycycline and CB-839 in cells lacking a doxycycline-inducible PLK4 construct. No significant differences were observed in colony formation across all three cell lines in the presence or absence of doxycycline (Fig. S1E). Together, these results indicate that, independent of KRAS mutation status, PDAC cells with PLK4-induced CA display increased sensitivity to GLS1 inhibition.

We next assessed the dependency for L-glutamine (L-gln) availability on the survival of cells with CA. Reducing L-gln concentrations in the culture medium significantly impaired cell viability in Panc1 and Mia Paca-2 cells. Notably, Panc1 cells with long-term CA (7d) displayed greater sensitivity to L-gln reduction compared to cells with short-term amplification (3d) (Fig. S1F). In colony formation assays, when CA was induced at the time of seeding, no colonies formed in the absence of Na-pyruvate (Na-pyr), regardless of L-gln concentration (Fig. S1G). While cells with long-term CA were able to tolerate Na-pyr withdrawal and still form colonies when supplemented with 4 mM L-gln, lowering L-gln levels markedly impaired colony formation (Fig. S1H-J). Importantly, reduced L-gln concentrations in the culture media had no significant effect on dox–control cells. Together, these results suggest that CA imposes an absolute requirement for Na-pyr at early stages, and that with prolonged amplification, cells accumulate further metabolic stress, resulting in a conditional dependency on extracellular L-gln. Taken together, these results suggest that L-gln metabolism is essential for the survival of the PDAC cells with CA.

### 2.2 Disruption of redox homeostasis pathways is lethal in cells with centrosome amplification

L-glutamine (L-gln) is a central metabolic node in PDAC cells. Following its conversion to L-glutamate (L-glu) by GLS1, it fuels multiple essential processes, including TCA cycle anaplerosis and glutathione (GSH) synthesis (Fig. 2A). To disentangle these functions, we examined the dual contributions of L-gln to core metabolism and stress adaptation. In particular, we assessed its entry into the TCA cycle and its role in GSH production, focusing on how these pathways intersect with NRF2-driven antioxidant responses, since CA elevates ROS and activates NRF2 signaling^14^ (Fig. 2B). Using the fluorogenic probe DCFDA, we detected a modest increase in intracellular ROS (∼1.5-fold) upon PLK4 induction in Panc1, Mia Paca-2, and BxPC-3 cells (Fig. 2C, 2D). Treatment with CB-839 or ML385 (NRF2 inhibitor) had little impact on ROS levels after 48 hours, whereas inhibition of glutathione synthesis with buthionine sulfoximine (BSO) caused marked ROS accumulation (Fig. S2A, S2B). Among the tested PDAC models, BxPC-3 cells with CA displayed the highest ROS levels across treatments, indicating greater sensitivity to oxidative stress (Fig. S2B). Consistent with this, PLK4 induction led to a decrease in the GSH:GSSG ratio in all three cell lines, reflecting increased utilization of glutathione to maintain redox balance under centrosome amplification (Fig. 2E).

**Figure 2.**
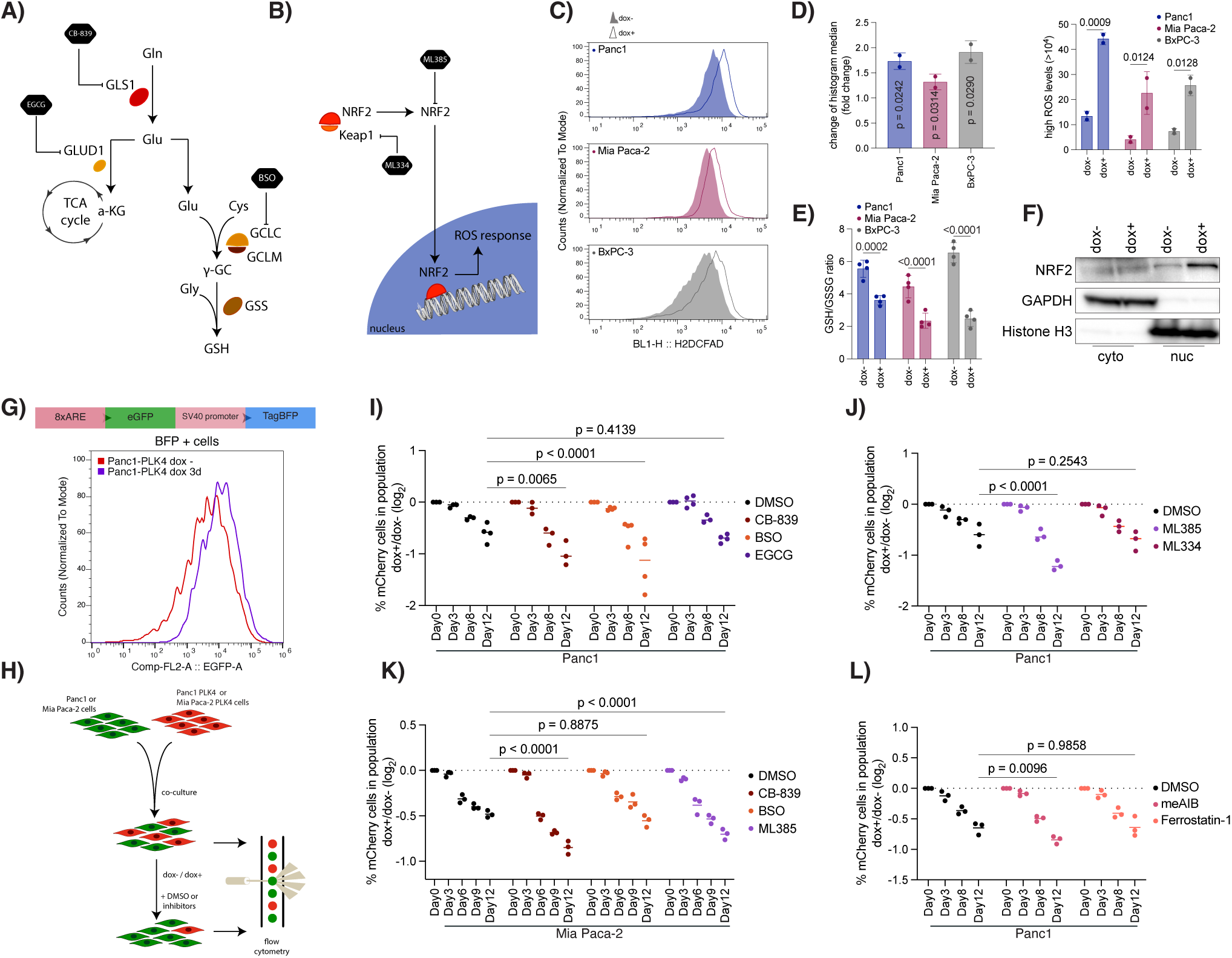
Centrosome amplification increases ROS, creating a vulnerability to ROS elimination pathway inhibition. A) Schematic representation of L-glutamine metabolism pathways and enzymes targeted by specific inhibitors in the following experiments. B) Diagram of the NRF2 signaling pathway and inhibitors targeting NRF2 and Keap1. C) CA increases intracellular ROS levels in Panc1, Mia Paca-2 and BxPC-3 cells. D) Quantification of ROS measurement results in 2C. Left panel: Changes in histogram median values. Right panel: Percentage of cells with high ROS levels. E) Induction of CA decreases GSH:GSSG ratios in PDAC cell lines. F) Induction of CA increases nuclear localization of NRF2 in Panc1 cells. GAPDH and Histone H3 blots represent cytoplasmic and nuclear fractionation. G) CA increases the Antioxidant Response Element (ARE)-mediated gene expression in Panc1 cells. H) Overview of the competition experiments performed in panels H-K. I-J) Treatment with CB-839, BSO and ML385 significantly reduces the viability of Panc1 cells with CA in in-vitro competition assays. K) CB-839 and ML385 treatments diminish the survival of Mia Paca-2 cells with CA in in vitro competition assays. L) Inhibition of SNAT1-mediated glutamine uptake reduces the viability of Panc1 cells with CA in in vitro competition assays. Statistical significances were measured by two-tailed t-test in D (left panel), and by two-way ANOVA in D (right panel), E, and I-L. Dots represent individual repeats. p values were reported on graphs.

As intracellular ROS accumulation triggers NRF2 activation via dissociation from Keap1 and nuclear translocation (Fig. 2B), we investigated the nuclear localization of NRF2. Western blot analysis showed increased nuclear NRF2 levels in dox-treated Panc1-PLK4 cells (Fig. 2F). Furthermore, CA increased antioxidant responsive element (ARE) activation in Panc1 cells, as demonstrated by plasmid-based reporter assays^24^ (Fig. 2G and Fig. S2C). Notably, doxycycline treatment of Panc1 cells lacking the dox-inducible PLK4 construct did not result in elevated ARE activity, confirming that the observed effect was specific to CA (Fig. S2D).

To assess the long-term requirement of L-gln metabolism and NRF2 pathway for the survival of PDAC cells with CA, we leveraged dual-color competition assays. We combined H2B-GFP expressing Panc1 and Mia Paca-2 cells with H2B-mCherry expressing dox-PLK4 counterparts, and tracked the changes of cell populations on different time points (Fig. 2H). This method offered two key advantages over traditional cell viability experiments: first, it enabled the assessment of long-term outcomes; and second, it eliminated potential confounding effects of drug-dox interactions, as both cell populations were exposed to identical concentrations of the drug and doxycycline simultaneously. In all long-term competition experiments, H2B-mCherry-expressing dox-PLK4 cells were progressively depleted over time in the DMSO treated control groups, highlighting the anti-proliferative impact of CA (Fig. 2I-L). Treatment of the mixed Panc1 cell populations with CB-839 or BSO led to a greater depletion of mCherry+ cells, whereas EGCG had no significant effect, suggesting that GLUD1-mediated incorporation of L-glutamine into the TCA cycle does not hold differential importance for cells with CA (Fig. 2I, Fig. S3A-C). Similarly, ML385 treatment caused a significant depletion of mCherry+ cells; however, inhibition of Keap1 by ML334 failed to rescue this depletion (Fig. 2J, Fig. S3D), possibly due to already elevated NRF2 nuclear localization in centrosome-amplified cells as a result of oxidative stress. In Mia Paca-2 cells, CB-839 and ML385 treatment also led to a marked depletion of mCherry+ cells, while BSO treatment did not cause a significant reduction compared to DMSO (Fig. 2K, Fig. S4A-C). Furthermore, BSO treatment impaired colony formation in Panc1 cells with CA, whereas ML385 treatment showed no notable effect on colony formation (Fig. S2E). In contrast, Mia Paca-2 cells were insensitive to BSO but showed marked sensitivity to ML385 in colony formation assays (Fig. S2F). These findings further suggest that distinct mechanisms of ROS elimination may be critical in different PDAC models. Specifically, one cell type may rely more heavily on GSH synthesis, while another may depend predominantly on NRF2 signaling. Additionally, the consistency between long-term competition assays and independent colony formation experiments supports that our dual-color competition strategy faithfully captures the true biological vulnerabilities of centrosome-amplified cells.

L-gln import in cancer cells predominantly relies on the ASCT2 (SLC1A5), SNAT1 (SLC38A1), and SNAT2 (SLC38A2) transport systems^25,26^. To evaluate the dependency of cells with CA on L-gln uptake, we used 2-methylamino isobutyrate (meAIB), a specific inhibitor of SNAT1/SLC38A1^27^, in dual-color competition experiments. Our findings revealed that SNAT1-mediated L-gln import is critically required for the survival of Panc1 cells with CA (Fig. 2L, Fig. S5). As glutathione depletion is a hallmark trigger of ferroptosis, we examined whether ferroptotic cell death could account for the CA-associated viability reduction. To this end, we treated cells with Ferrostatin-1, a well-characterized inhibitor of ferroptotic cell death^28^. Nevertheless, Ferrostatin-1 treatment did not prevent the depletion of mCherry+ cells, indicating that ferroptosis is unlikely to play a role (Fig. 2L, Fig. S5). Collectively, these findings underscore the critical dependence of PDAC cells with CA on L-glutamine import, glutathione biosynthesis, and NRF2-mediated antioxidant signaling for their survival.

### 2.3 Metabolism focused CRISPR screen identifies metabolic dependencies of cells with centrosome amplification

To further delineate the metabolic dependencies of cells with CA, we performed a metabolism-focused CRISPR-Cas9 screen in Panc1-PLK4 cells (Fig. 3A). Cells were transduced with a pooled CRISPR library targeting metabolic enzymes at a low multiplicity of infection (MOI = 0.6) and cultured for 21 days under doxycycline-treated and untreated conditions. sgRNA abundances in the final populations were compared to the initial library transduced cells to determine gene-level essentiality. Comparative analysis of beta-scores revealed that while most genes exhibited similar depletion profiles under both conditions, a distinct subset showed differential depletion, suggesting CA-specific metabolic vulnerabilities (Fig. 3B, 3C). Gene Set Enrichment Analysis (GSEA) of the ranked gene list highlighted consistent enrichment of two recurring pathways across multiple terms: (i) nucleotide sugar and N-glycan biosynthesis, and (ii) reactive oxygen species (ROS) detoxification pathways (Fig. 3D). Furthermore, by applying a threshold of two median absolute deviations (2-MAD), we identified 109 genes as significantly depleted in the centrosome-amplified condition, which were used for downstream pathway-level analyses (Fig. S6A). Among these genes, 80 have well-characterized intracellular localizations according to the Human Protein Atlas (HPA) database^29^. The majority were localized to the cytoplasm (33) and nucleoplasm (27), with 15 associated with mitochondria, 15 with the plasma membrane, and 3—CPT1C, CPE, and ENGASE—reported to localize to the centrosome (Fig. S6B). Pathway enrichment analysis of this gene set revealed significant over-representation of the reactive oxygen species pathway, as well as glycosaminoglycan and nucleotide sugar biosynthesis pathways, based on GO-BP, MSigDB, and KEGG analyses (Fig. S6C). Furthermore, transcription factor enrichment analysis using the TR-RUST database identified MTF1 and NFE2L2 (NRF2) as key upstream regulators, suggesting these factors may orchestrate survival-associated transcriptional programs in centrosome-amplified cells (Fig. S6C). Integration of our results with previously identified dependencies from Panc1 CRISPR screens (DepMap) revealed that while several top-depleted genes such as SOD1, SOD2, and DPAGT1 overlapped with known dependencies, many of our differentially depleted hits including GFPT2, TXNRD2, PRDX1, CHST7, SLC5A3, and UGDH were not classified as common dependencies (Fig. 3E). Markov Cluster Algorithm (MCL) clustering of top differentially depleted genes revealed several protein associations, further suggesting pathway-level dependencies (Fig. 3F, 3G).

**Figure 3.**
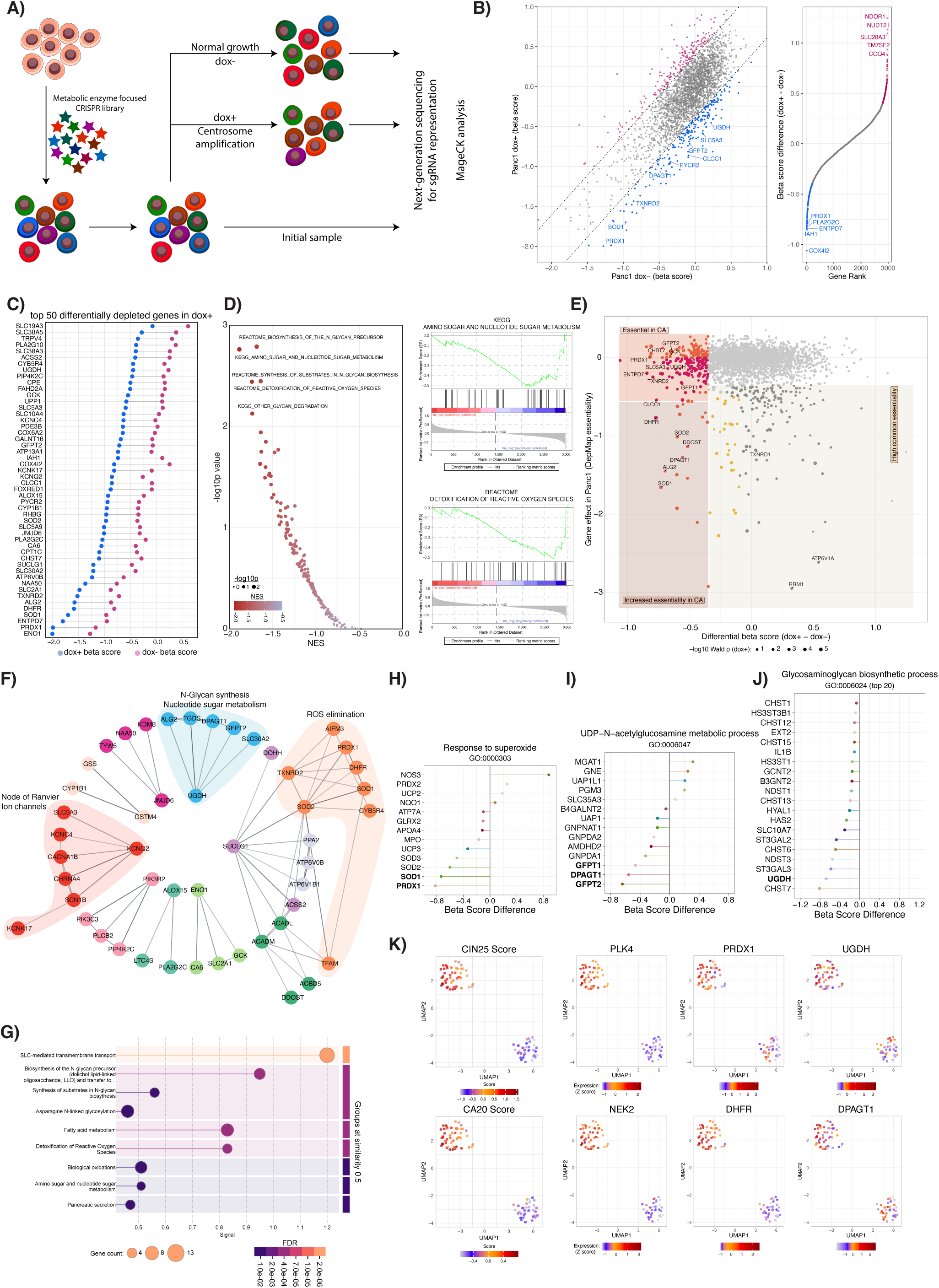
Metabolism targeted CRISPR screen identifies increased dependency for ROS detoxification and nucleotide sugar metabolism in cells with PLK4-induced supernumerary centrosomes. A) Schematic representation of metabolic enzyme focused CRISPR screen experiment design. B) Top depleted hits in dox+ and dox- cells compared to initial sample. Left panel: Scatterplot of beta scores for dox+ and dox- sample. Pink dots in the scatterplot represent genes with a beta score that increased after CA. Blue dots represent genes with a beta score that decreased after CA. Right panel: Rank plot showing the genes based on differential beta score in which dox- beta score is subtracted from the dox+ beta score. C) Top 50 differentially depleted genes in dox+ samples. Pink dots represent beta score in dox- comparison, blue dots represent beta score in dox+ comparison. D) GSEA analysis of CRISPR screen results. E) Comparison of differential beta score values of CRISPR screen with Panc1 DepMap essentialities. F) MCL clustering results of top differentially depleted metabolic genes in cells with CA. Genes that were not included in a cluster (singletons) and clusters contain less than three proteins were removed. G) Enrichment analysis of protein-protein interaction network. H-J) Pathway-specific differentially depleted genes in cells with CA. (H) Response to superoxide (GO:0000303). (I) UDP-*N*-acetylglucosamine metabolic process (GO:0006047). (J) Glycosaminoglycan biosynthetic process (GO:0006024). K) UMAP projection of TCGA PDAC data for selected genes. CIN25 and CA20 gene expression scores was shown on the left side plots. PLK4 and NEK2: CA20 genes; PRDX1 and DHFR: ROS elimination; UGDH and DPAGT1: N-glycan synthesis/nucleotide sugar metabolism. Gene expression Z-scores were used in plots.

To further investigate the role of metabolic enzymes in specific pathways and cellular functions, we analyzed the CRISPR screen results by filtering for targeted Gene Ontology (GO) terms (Fig. 3H-J & Fig. S6D-G). Among the genes involved in the cellular response to superoxide, PRDX1, SOD1, SOD2, and SOD3 were significantly depleted in cells with CA (Fig. 3H). Similarly, within the N-acetylglucosamine metabolic process, GFPT2, DPAGT1, and GFPT1 emerged as top-depleted genes, while CHST7 and UGDH were identified as key hits in glycosaminoglycan biosynthesis (Fig. 3I, 3J). Additionally, GFPT2 and GFPT1 were prominent among enzymes that have role in glutamine metabolic processes (Fig. S6D), while GALNT16, UGT1A7, and UGT1A8 were top hits among enzymes functioning as UDP-glycosyltransferases (Fig. S6E). PYCR2 was the sole depleted gene identified in the proline biosynthesis pathway (Fig. S6F), whereas GSS, GCLC, and MGST2 were highlighted in the glutathione biosynthesis process (Fig. S6G). To extend our findings, we integrated the top CA-specific differentially depleted genes from our CRISPR screen with results from two previously conducted unbiased gene expression studies in cells that have CA^14,30^. Additionally, we incorporated a curated list of NRF2-regulated genes^31^ into the analysis, motivated by our observation of increased NRF2 nuclear localization in centrosome-amplified cells (Fig. 2E). This integrative approach identified UGDH as a shared hit across all datasets from multiple cell lines (Fig. S6H). Furthermore, filtering the screen results for NRF2-regulated genes highlighted DHFR and UGDH as top differentially depleted hits in cells with CA (Fig. S6I).

Additionally, we analyzed TCGA patient data to evaluate whether the expression of top CA–specific hits (differential LFC < −0.5) from our CRISPR screen correlated with chromosomal instability (CIN25)^32^ and centrosome amplification (CA20)^33^ transcriptional signatures. We also included NRF2 (NFE2L2), ATF4, and ATF6 expression, given their central roles in cellular stress responses. Unsupervised clustering of pancreatic adenocarcinoma samples (n = 82) revealed that CIN25 and CA20 expression profiles were the dominant factors driving patient stratification within this gene set (Fig. S7A). UMAP projection yielded a comparable distribution of samples, again primarily structured by CIN25 and CA20 expression (Fig. 3K). Notably, patient subsets with high CIN25 and CA20 scores exhibited elevated NRF2 and ATF4 expression, suggesting a potential link between genomic instability, CA, and activation of stress response pathways (Fig. S7A). Furthermore, several top hits from our CRISPR screen—including DHFR, GSTM4, UGDH, SOD1, PRDX1, and DPAGT1—were highly expressed in patients with elevated CIN25 and CA20 signatures, highlighting their potential clinical relevance as metabolic requirements in genomically unstable pancreatic tumors (Fig. 3K, S7A). Expression of DHFR, DPAGT1, SLC5A3, PRDX1, and UGDH also correlated strongly with PLK4 levels (Fig. S7B, S7C, 3K), further strengthening the connection between these metabolic genes and CA in patient tumors. To gain additional insight into the transcriptional programs associated with PLK4, we performed gene set enrichment analysis (GSEA) using the GENI platform^34^. As expected, PLK4 expression was significantly enriched for cell cycle–related pathways, including E2F targets, G2/M checkpoint, and mitotic spindle (Fig. S7D). Intriguingly, enrichment was also observed for pathways involved in the unfolded protein response (UPR), protein secretion, and interferon signaling, indicating that PLK4 over-expression may engage broader stress-response and immune signaling programs in pancreatic tumors (Fig. S7D).

To better understand the CA–associated dependencies, we also analyzed the expression changes of a selected gene panel following CA in Panc1 and Mia Paca-2 cells. After three days of doxycycline induction, both cell lines showed modest increases in PRDX1, DPAGT1, and GFPT2 expression, accompanied by a slight reduction in HAS2 expression (Fig. S6J). Cell line–specific responses were also observed: GLUL was up-regulated only in Mia Paca-2 cells, whereas CHST7, MGST2, and GALNT16 showed increased expression exclusively in Panc1 cells (Fig. S6J).

Altogether, our analyses highlight ROS detoxification and nucleotide sugar/glycan biosynthesis as key metabolic dependencies in cells with PLK4-induced CA, suggesting their potential as therapeutic vulnerabilities in cancers characterized by CA and genomic instability.

### 2.4 Inhibition of ROS elimination pathways is selectively lethal in cells with centrosome amplification

Given that ROS elimination pathways were among the top enriched terms in our list of genes selectively depleted in centrosome-amplified cells—and that key antioxidant genes such as SOD1, PRDX1, and TXNRD2 were prominently featured—we initially focused our analysis on this axis. The thioredoxin system, which includes thioredoxin and thioredoxin reductases such as TXNRD2, plays a critical role in regulating protein redox status by facilitating disulfide bond reduction and maintaining proteins in a reduced state^35^. In parallel, superoxide dismutases (SODs) catalyze the conversion of superoxide anions into hydrogen peroxide, thereby mitigating oxidative stress and preventing ROS-mediated cellular damage^36^ (Fig. S8A). Therefore, we employed LCS-1, a potent and selective SOD1 inhibitor, and auranofin, a clinically approved thioredoxin reductase inhibitor, in competition experiments. Both treatments led to an increased reduction in cell populations with CA in Panc1 cells (Fig. S8B, S9A), highlighting the critical role of these antioxidant systems in maintaining the survival of these cells. In contrast, treatment with N-acetyl cysteine (NAC) or apocynin, which broadly scavenges ROS or inhibits NADPH oxidase, had no impact on the cell viability of centrosome-amplified cells in long-term competition experiments. These results suggest that centrosome-amplified cells depend specifically on enzymatic ROS detoxification mechanisms, rather than general oxidative stress buffering, for their survival (Fig. S8C, S9B).

Additionally, DHFR (dihydrofolate reductase) emerged as one of the top depleted hits in centrosome-amplified cells (Fig. 3C, 3E, 3F), it ranked highest among NRF2 target genes (Fig. S6I), and showed a high correlation with PLK4 expression in TCGA mRNA expression data (Fig. S7A and S7B). DHFR catalyzes the conversion of dihydrofolate to tetrahydrofolate, a key step in folate metabolism, and has been previously linked to the regulation of cellular redox balance^37^(Fig. S8D). To evaluate its functional relevance, we inhibited DHFR using pralatrexate in competition assays, which led to a greater depletion of centrosome-amplified Panc1 cells compared to controls (Fig. S8E, S9C). Furthermore, DHFR inhibition in these cells resulted in elevated ROS levels (Fig. S8F and S8G), supporting its requirement in maintaining redox homeostasis under CA-induced stress. Competition experiments employing LCS-1, auranofin, and pralatrexate in MiaPaCa-2 cells recapitulated the results observed in Panc1 cells (Fig. 8H, S10). Consistent with this, DHFR inhibition in MiaPaCa-2 cells with CA similarly increased intracellular ROS levels (Fig. S8I and S8J), confirming that DHFR contributes to redox homeostasis across multiple CA models. Together, these findings establish ROS detoxification as critical survival dependencies in centrosome-amplified cells.

### 2.5 PDAC cells with centrosome amplification have increased dependency for uronic acid and hexosamine biosynthetic pathways

Analysis of differentially depleted sgRNAs in centrosome-amplified cells revealed a strong enrichment for genes involved in nucleotide sugar metabolism and N-glycan biosynthesis. Among these, UGDH, GFPT1, GFPT2, and DPAGT1 emerged as top hits, highlighting dependencies in the uronic acid and hexosamine biosynthesis pathways (Fig. 3I, J). These pathways generate essential nucleotide sugars required for protein glycosylation, glycosaminoglycan biosynthesis, and hyaluronic acid production, processes that may buffer CA-induced proteotoxic and mechanical stress^38,39^. To functionally characterize these dependencies, we pharmacologically inhibited key enzymes in these pathways using Azaserine, 4-MU, FR054, Tunicamycin, and OSMI-1 in the competition experiments (Fig. 4A). Treatment with FR054, 4-MU, and Tunicamycin led to greater selective depletion of centrosome-amplified cells compared to DMSO treatment, providing evidence for increased dependency (Fig. 4B, S11). Although Azaserine and OSMI-1 induced substantial cell death, this effect was not specific to the centrosome-amplified population. These results demonstrate that CA imposes a specific requirement for uronic acid and hexosamine pathway activity and for N-linked glycosylation, while no specific dependence for O-linked glycosylation was observed under tested conditions.

**Figure 4.**
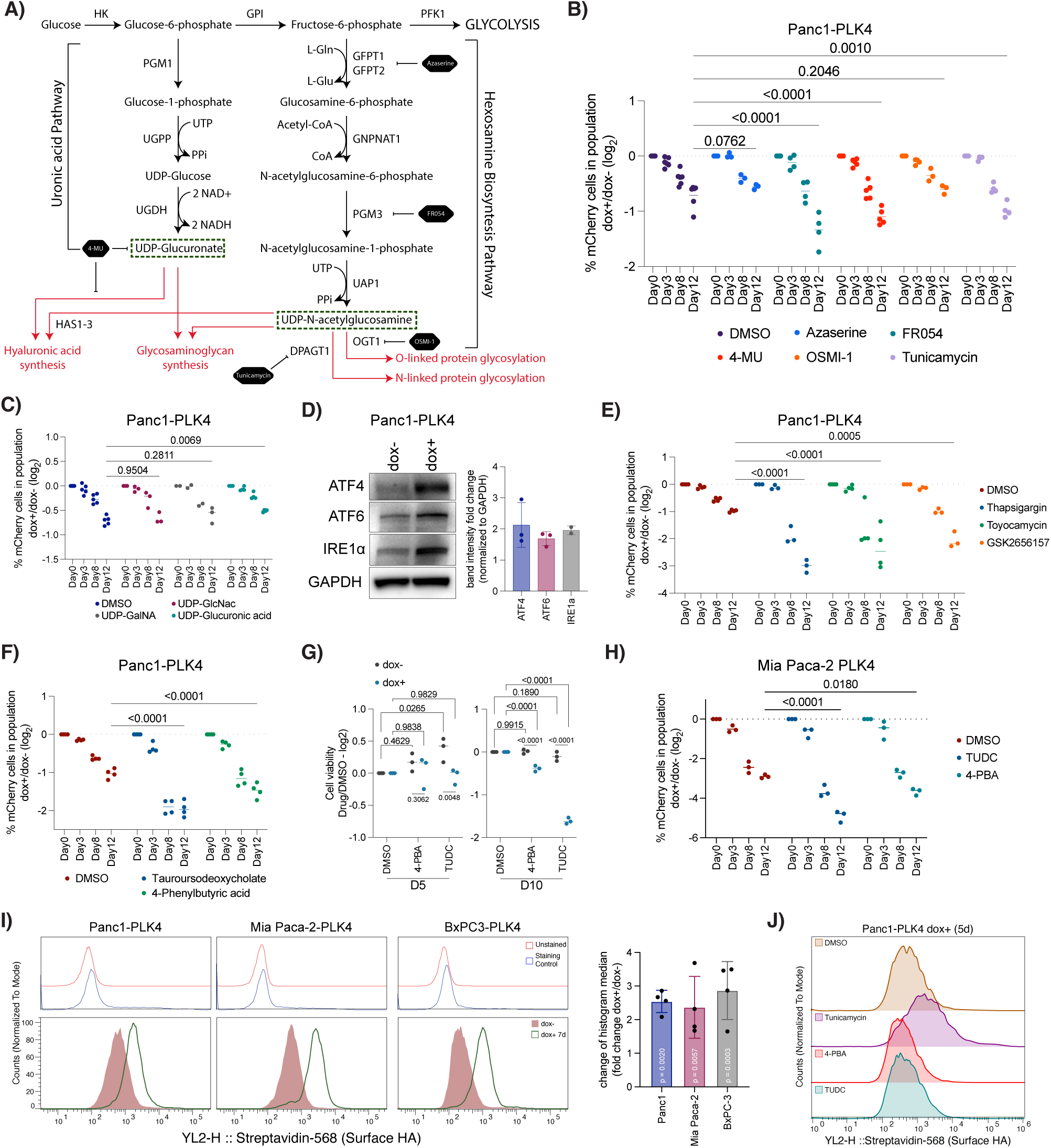
Cells with centrosome amplification has increased dependency for uronic acid and hexosamine pathways. A) Schematic representation of uronic acid and hexosamine biosynthesis pathways and enzymes targeted by specific inhibitors in the following experiments in Fig. 4B. Green boxes show metabolites that were used in competition experiments in Fig. 4C. B) 4-MU, FR054 and Tunicamycin treatment reduces viability of the cells with CA more compared to normal counterparts. C) Supplementation of UDP-glucuronic acid reduces depletion of cells with CA in competition experiments. D) CA induces ER stress / UPR associated protein levels. Left panel: A representative western blot result. GAPDH was used as loading control. Right panel: Quantification of the western blot results. Dots represent independent experiment repeats. E) Induction of ER-stress and disruption of UPR signaling mechanisms increase the depletion of cells with CA in competition experiments. F) Reduction of ER-stress by TUDC and 4-PBA increase the depletion of cells with CA in competition experiments. G) Reduction of ER-stress by TUDC and 4-PBA decrease viability of cells with CA. H) Reduction of ER-stress by TUDC and 4-PBA increase the depletion of cells with CA in competition experiments in Mia Paca-2 cells. I) CA increases cell surface hyaluronic acid levels in PDAC cells. Left panel: Cell surface hyaluronic acid levels in control and dox+ PDAC cell lines. Right panel: Quantification of histogram median shift fold changes. Statistical significances were measured by two-way ANOVA in B, C, E, F, G, and H, by two-tailed t-test in I. Dots represent independent experiment repeats. p values were reported on graphs.

The functional interpretation of negative selection in metabolic CRISPR screens can be complex; depletion of a metabolic enzyme could indicate either a critical dependence on its product for survival or a toxic buildup of its substrate. To distinguish between these models for the nucleotide sugar pathway hits, we supplemented cells with the products of the pathways: UDP-GlcNAc (Uridine diphosphate-N-acetyl-glucosamine), UDP-GalNAc (UDP-N-acetyl-galactosamine), and UDP-glucuronic acid. Among these, UDP-glucuronic acid was the only sugar that partially rescued the depletion of centrosome-amplified cells (Fig. 4C, S13A). The hexosamine biosynthesis pathway includes well-characterized salvage routes that allow cells to re-utilize sugar metabolites to maintain UDP-sugar pools. Free N-acetylglucosamine (GlcNAc), derived from glycoconjugate degradation or extracellular uptake, can re-enter the pathway via phosphorylation by NAGK^40^. Similarly, glucosamine and N-acetylgalactosamine (GalNAc) can be salvaged and funneled into the synthesis of UDP-GlcNAc and UDP-GalNAc (Fig. S12A). To test whether salvage pathway metabolites promote the survival of centrosome-amplified cells, we treated cells with glucosamine and GlcNAc. Interestingly, only glucosamine supplementation improved the survival of centrosome-amplified cells in competition experiments (Fig. S12B, S13B).

Since tunicamycin-induced DPAGT1 inhibition is a well-established method for studying ER stress-induced activation of the unfolded protein response (UPR), and given that tunicamycin exerted a stronger effect on centrosome-amplified cells (Fig. 4B), we examined the cellular response to ER stress and UPR activation in cells with CA. Western blot analysis showed increased ATF4, ATF6, and IRE1α abundance in PLK4-induced CA cells, consistent with engagement of the UPR (Fig. 4D). Additionally, increased nuclear localization of ATF4 was observed (Fig. S12C), further supporting activation of the PERK branch of the UPR. To evaluate the functional importance of UPR signaling, we treated cells with (i) thapsigargin, a SERCA inhibitor, (ii) toyocamycin, an inhibitor of IRE1-mediated XBP1 splicing, and (iii) GSK2656156, an ATP-competitive PERK inhibitor. In competition experiments, all three treatments led to increased depletion of centrosome-amplified cells compared to DMSO controls (Fig. 4E, S14A). These results suggest that all three canonical branches of the UPR; IRE1, PERK, and ATF6, contribute to the adaptive stress response that supports the survival of PLK4-induced centrosome-amplified cells.

We next reduced ER stress by treating cells with Tauroursodeoxycholate (TUDC, a chemical chaperone that alleviates ER stress) and 4-Phenylbutyric acid (4PBA, an ER stress–reducing agent) in competition experiments, which also resulted in depletion of centrosome-amplified cells (Fig. 4F, S14B). Although this outcome may initially seem counterintuitive, it is consistent with the dual role of the UPR, which can either promote survival or trigger apoptosis depending on the intensity and context of ER stress^41^. By facilitating the protein folding and reducing misfolded protein burden, TUDC and 4-PBA likely attenuate the adaptive, pro-survival arm of the UPR. Because competition experiments could reflect either a true reduction in the proliferation of centrosome-amplified cells or a relative effect caused by increased proliferation of non-PLK4 over-expressing cells, we tested individually seeded cell populations. This confirmed a genuine reduction, as TUDC and 4-PBA treatments decreased the proliferation specifically in centrosome-amplified cells (Fig. 4G). Importantly, similar results were observed in Mia Paca-2 competition experiments (Fig. 4H, S15A), highlighting that centrosome-amplified cells may require a finely tuned level of UPR activity for viability.

Since the glucuronic acid and hexosamine biosynthesis pathways contribute to hyaluronic acid (HA) synthesis (Fig. 4A), and we observed increased depletion with 4-MU (Fig. 4B), we next examined HA production in these cells. Because HA is secreted following synthesis and associates with the cell surface, we measured surface HA levels and observed significantly higher levels in cells with CA (Fig. 4I). Importantly, doxycycline treatment in cells lacking the doxycycline-inducible PLK4 construct did not increase HA levels, confirming that this effect stems from CA rather than doxycycline exposure (Fig. S12E). In competition assays, adding exogenous HA to the cell culture media did not improve the survival of centrosome-amplified cells (Fig. S12E, S15B), but it rescued the depletion caused by 4-MU treatment, though not by tunicamycin (Fig. S12F, S16). Given the established link between the unfolded protein response and hexosamine pathway activity^42,43^, we next asked whether ER stress modulates HA production in centrosome-amplified cells. Pharmacological induction of ER stress with tunicamycin increased surface HA, whereas reducing ER stress with 4-PBA or TUDC lowered HA levels in centrosome-amplified Panc1 cells (Fig. 4J, S12G, S12H). Together, these findings indicate that CA enhances HA synthesis and couples it to ER-stress status, creating a metabolic requirement that supports the survival of centrosome-amplified PDAC cells.

In addition, our CRISPR screen identified the myo-inositol transporter SLC5A3 as a selective dependency in CA cells (Fig. 3B, C, E, F), consistent with its proposed oncogenic role in other cancers^44,45^. Imported myo-inositol contributes to phosphatidylinositol synthesis or can be oxidized to D-glucuronate, a potential entry point into the pentose phosphate pathway (Fig. S12I). Although D-glucuronate can be converted to UDP-glucuronate in some species, this pathway is absent in humans due to the lack of UDP-glucuronate dehydrogenase (UGD)^46^. Supplementation with either myo-inositol or D-glucuronate significantly increased the survival of centrosome-amplified cells in competition assays (Fig. S12J, S17), indicating an increased requirement for these metabolites. Expression analysis further revealed strong down-regulation of IMPA1, which encodes a key enzyme for endogenous inositol biosynthesis, in Mia PaCa-2 cells with CA (Fig. S12K). While Panc1 cells did not show altered IMPA1 expression, they nevertheless remained dependent on exogenous myo-inositol.

In summary, our data show that CA increases dependence on nucleotide-sugar biosynthesis, hyaluronic acid production, and extracellular myo-inositol uptake. These metabolic alterations coincide with elevated ER stress, activation of all three UPR branches, and a requirement for adaptive stress signaling to maintain cell survival.

### 2.6 Disruption of hyaluronic acid synthesis triggers cytokinesis failure in centrosome-amplified cells

To gain mechanistic insight into how sugar metabolism and glycosylation pathways support centrosome-amplified cells, we examined 4-MU (glucuronic acid metabolism), tunicamycin (N-linked protein glycosylation), and FR054 (hexosamine biosynthesis). DNA content analysis revealed shifts in cell-cycle profiles in both centrosome-amplified (dox+) and non-amplified (dox–) cells (Fig. 5A). Changes in the G1 peak indicated cell cycle (DNA content per cell) abnormalities (Fig. 5B), while the appearance of sub-G1 peaks, particularly in centrosome-amplified Mia Paca-2 cells (Fig. 5A, S18A), reflected increased apoptotic cell death. To better understand the specific effects in centrosome-amplified cells, we compared cell viability at different time points across three cell lines. Tunicamycin significantly reduced proliferation in Panc1 and Mia Paca-2 cells after five days of treatment, whereas BxPC-3 cells showed no differential effect (Fig. 5C). In contrast, 4-MU selectively impaired the viability in centrosome-amplified cells across all three lines (Fig. 5D), prompting further investigation. After 10 days of treatment with 4-MU, confocal imaging revealed pronounced increases in cell size and the accumulation of multinucleated cells, a definitive indicator of cytokinesis failure and impaired proliferative capacity (Fig. 5E, S18B). Quantification confirmed that 4-MU treatment did not alter the frequency of CA itself but significantly increased the proportion of multinucleated cells (Fig. 5F). These findings suggest that suppression of glucuronic acid–dependent HA synthesis perturbs not only extracellular matrix interactions but also intracellular processes critical for mitotic fidelity.

**Figure 5.**
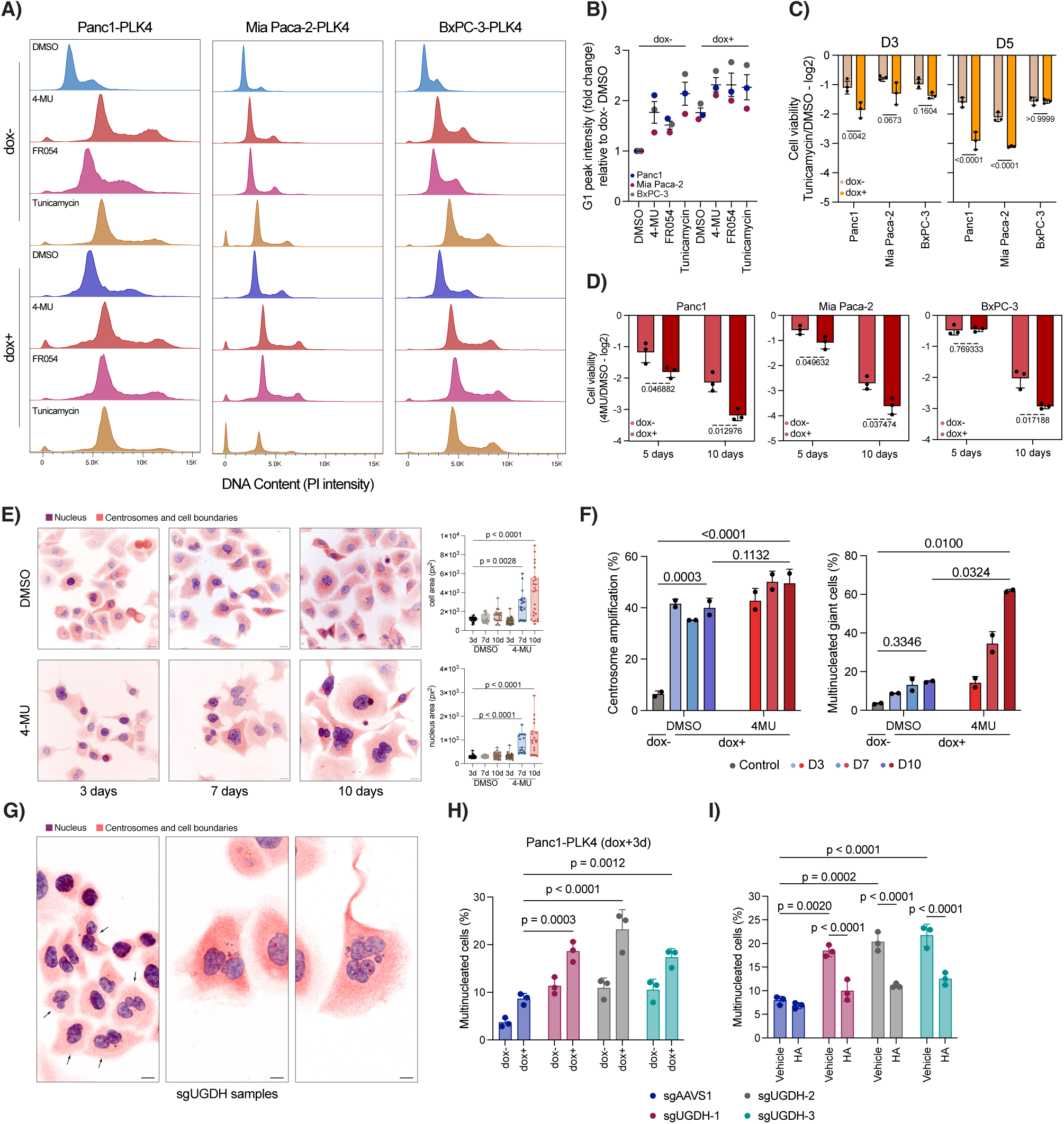
Chemical and genetic perturbation of hyaluronic acid synthesis induces multinucleation in pancreatic cancer cells with centrosome amplification. A) 4-MU, tunicamycin and FR054 treatments increase DNA content of individual cells. B) Quantification of the G1-peak intensities in Fig. 5A. C) Tunicamycin treatment significantly reduces proliferation of centrosome-amplified Panc1 and Mia Paca-2 cells compared to control. D) 4-MU treatment significantly reduces proliferation of centrosome-amplified Panc1, Mia Paca-2, and BxPC-3 cells compared to control. E) Long-term 4-MU treatment results in generation of multinucleated cells. Left panel: Inverted confocal images. Purple color shows DNA content of the cells, and orange color shows centrosomes and cell boundaries. Scale bar: 20 µm. Right panel: Quantification of cell area and nucleus area in pixel squares. F) Quantification of multinucleated giant cells in DMSO and 4-MU treated cells with CA. Left panel: Quantification of CA. Right panel: Quantification of multinucleated cells. G) CRISPR/Cas9 targeted disruption of UGDH gene results in generation of multinucleated cells. Purple color shows DNA content of the cells, and orange color shows centrosomes and cell boundaries. Scale bar: 20 µm. H) Quantification of multinucleated cells in sgAAVS1 and sgUGDH expressing centrosome-amplified and control cells. I) Quantification of multinucleated cells in sgAAVS1 and sgUGDH expressing HA or Vehicle treated cells with CA. Significance was determined by two-tailed t-test in C, by two-way ANOVA test in D, F, H, and I, by one-way ANOVA in E. Dots represent individual repeats. p values were reported on graph.

To directly test whether disruption of HA biosynthesis contributes to the generation of multinucleated cells, we performed genetic perturbation of UGDH using individual sgRNAs. In sgUGDH-transduced cells, we observed increased multinucleation even in non-centrosome–amplified cells, with a higher percentage after CA (Fig. 5H, S18C). Consistent with its role in HA synthesis, sgUGDH was also associated with reduced surface HA levels (Fig. S18D). Importantly, supplementation with exogenous HA partially rescued the multinucleation phenotype in sgUGDH cells (Fig. 5I), supporting the conclusion that impaired HA production contributes to cytokinesis defects.

Collectively, our data show that disruption of nucleotide sugar–dependent pathways, including glucuronic acid–mediated HA synthesis, exerts a greater inhibitory effect on PLK4-induced centrosome-amplified pancreatic cancer cells. This metabolic interference not only diminishes proliferative capacity, but also provokes cytokinesis defects, establishing a mechanistic link between metabolic dependencies, mitotic fidelity, and cell survival that may be leveraged therapeutically.

### 2.7 CD44 activation contributes to centrosome clustering in PDAC cells with centrosome amplification

Since surface HA levels were elevated in centrosome-amplified PDAC cells, we next examined the expression of major HA-binding receptors. Among CD44, RHAMM, LYVE1, and HARE, CD44 is the most abundantly expressed in PDAC and is the main mediator of HA-dependent signaling^47–49^. RHAMM can cooperate with CD44 by promoting its surface localization and stabilizing HA binding, particularly when HA is immobilized^50^. However, RHAMM lacks a trans-membrane domain and also carries out intracellular functions^49,51^. Based on these features, we focused our analysis on CD44 as the primary receptor for HA signaling in PDAC cells.

To test whether CA alters CD44 expression, we first analyzed an unbiased gene expression dataset^14^, which revealed significant up-regulation of CD44 in centrosome-amplified cells (Fig. S19A). Consistently, CD44 and RHAMM (HMMR) expression levels were positively correlated in TCGA PDAC samples (Fig. S19B). Flow cytometry further confirmed increased surface CD44 levels in centrosome-amplified Panc1, Mia Paca-2, and BxPC-3 cells (Fig. 6A, S19C). Notably, BxPC-3 cells exhibited the greatest increase, whereas Mia Paca-2 cells showed a more modest elevation among the tested cell lines (Fig. S19C). Finally, doxycycline treatment alone in PDAC cells lacking the doxPLK4 construct did not affect CD44 levels (Fig. S19D, S19E), confirming that CD44 up-regulation is specifically driven by CA. To assess the role of CD44 in the survival of centrosome-amplified cells, we sorted subpopulations of Panc1-PLK4 and BxPC-3-PLK4 cells with high (top 10%) or low (bottom 10%) CD44 surface expression, induced CA, and monitored cell proliferation for 10 days. CD44-low Panc1-PLK4 cells exhibited markedly reduced survival compared to CD44-high cells (Fig. 6B). In contrast, this effect was not observed in BxPC-3-PLK4 cells, suggesting that CD44 dependency may be context-specific, potentially influenced by *KRAS* mutation status. To directly evaluate the requirement of CD44, we generated CD44-KO cells by targeting CD44 with three different sgRNAs and isolating populations that lost CD44 surface expression by FACS (Fig. S19F). Consistently, loss of CD44 further compromised the survival of centrosome-amplified Panc1 cells (Fig. 6C), reinforcing its role in supporting tolerance to CA.

**Figure 6.**
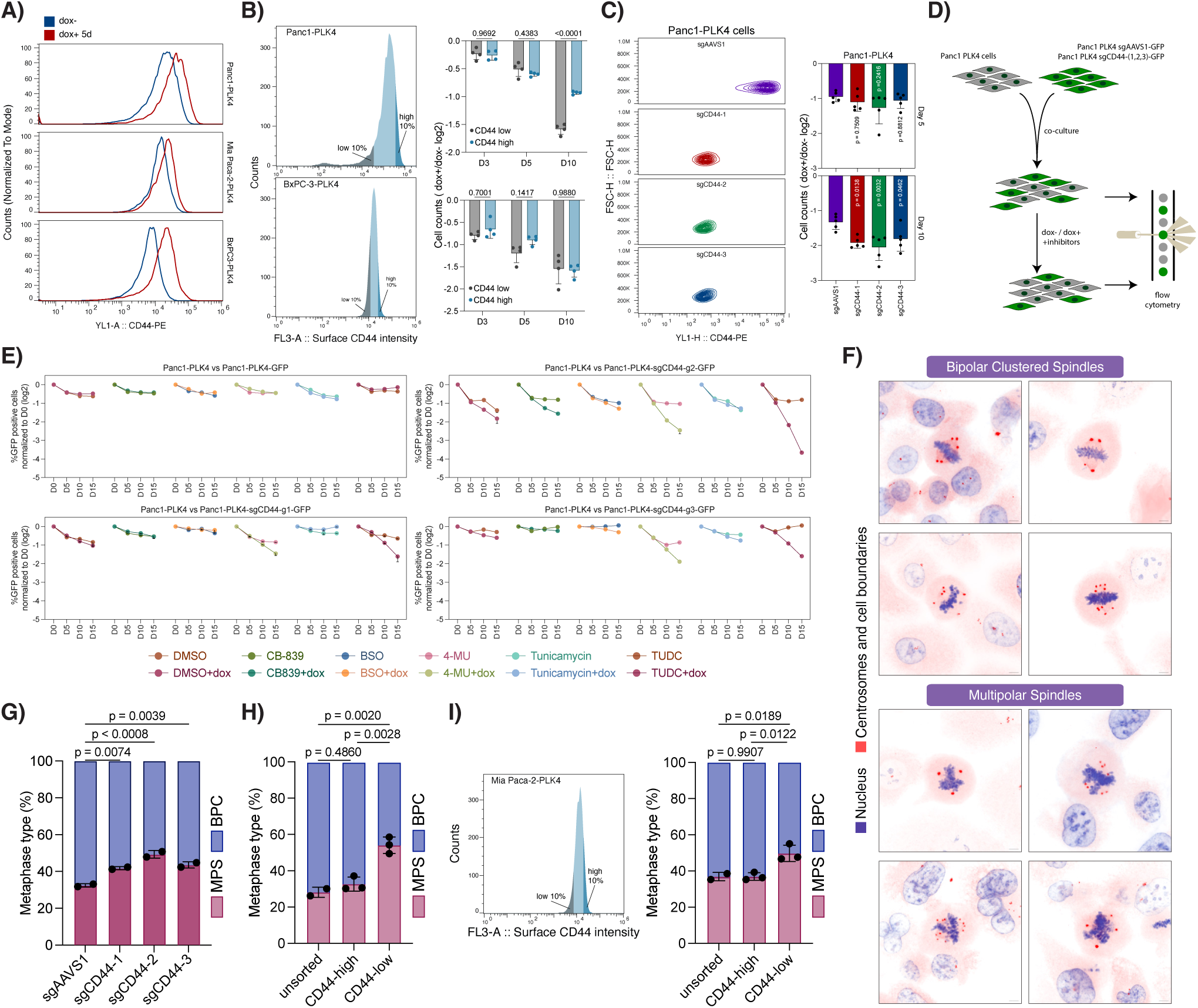
Cell surface CD44 is increased upon centrosome amplification and contributes to centrosome clustering in PDAC cells. A) Flow cytometry analysis showing elevated CD44 surface levels in centrosome-amplified PDAC cells. B) CD44-low Panc1 cells are more sensitive to CA. Left panel: FACS-sorted CD44-low and CD44-high populations in Panc1 and BxPC-3 cells. Right panel: Cell proliferation following different durations (3, 5, and 10 days) of CA. C) CD44-KO Panc1 cells are more sensitive to CA. Left panel: Flow cytometry confirming loss of CD44 expression in CD44-KO cells. Right panel: Cell proliferation following CA (5 and 10 days). D) Schematic of the competition assay design used in panel E. E) CD44-KO generates an increased vulnerability for UPR reduction in centrosome-amplified Panc1 cells. F) Representative confocal images of metaphase spindle organizations in Panc1 cells with CA. Top panel: bipolar clustered spindles; Bottom panel: multipolar spindles. G) Quantification showing reduced centrosome clustering in CD44-KO Panc1-PLK4 cells. H) CD44-low Panc1-PLK4 cells have increased multipolar spindle formation in metaphase. I) CD44-low Mia Paca-2-PLK4 cells have increased multipolar spindle formation in metaphase. Significance was determined by two-way ANOVA test in B, by one-way ANOVA in C, G, H, and I. Dots represent individual repeats. p values were reported on graph.

Because CD44 function is regulated through alternative splicing, with the standard isoform (CD44s) and variant isoforms (CD44v) linked to distinct cancer phenotypes in PDAC^52^, we next examined whether CA alters CD44 splicing. RT-PCR analysis revealed that splicing patterns of CD44 were unchanged in centrosome-amplified Panc1 and MiaPaCa-2 cells, whereas induction of variant isoforms was observed in BxPC-3 cells upon CA (Fig. S20A). This BxPC-3–specific shift toward CD44v may underlie their reduced reliance on overall CD44 levels for survival, providing a potential explanation for the context-dependent requirement of CD44 in tolerating CA. Since CD44v has been linked to redox regulation through stabilization of the cystine/glutamate antiporter xCT and support of glutathione synthesis^53^, we tested whether this splicing shift affected ROS levels. In Panc1 cells, which predominantly expressed CD44s even after CA, CD44 knockout did not alter ROS levels (Fig. S20B). By contrast, in BxPC-3 cells, where CA induced CD44v expression, CD44 loss led to a modest but significant increase in ROS (Fig. S20C), consistent with a partial role of CD44v in redox control. This difference may reflect how KRAS genetic context modulates the cellular response to centrosome amplification, including ROS regulation.

To directly assess the functions of CD44 in the context of CA, we performed competition assays by mixing CD44-KO (sg1–3) Panc1-PLK4-GFP cells with parental Panc1-PLK4 cells and treating them with inhibitors to evaluate whether they became more sensitive to glutamine utilization and ROS modulation (CB-839, BSO), inhibition of hyaluronic acid synthesis (4-MU), increased ER stress (tunicamycin), or reduced UPR signaling (TUDC) (Fig. 6D). Results revealed that CD44-KO cells were associated with increased sensitivity to 4-MU and TUDC only, suggesting that CD44 contributes to stress adaptation when UPR signaling is reduced by TUDC treatment. Notably, the increased 4-MU sensitivity suggests that HA remains functionally important even in the absence of CD44, consistent with HA signaling through additional receptors (Fig. 6E). As a control, sgAAVS1-GFP transduced Panc1-PLK4 cells were not differentially depleted under any treatment, with or without doxycycline induction (Fig. 6E). We also confirmed reduced survival of CD44-KO cells upon TUDC treatment in individually seeded cell groups. Although 4-MU also impaired the survival of CD44-KO cells, TUDC exhibited a stronger differential effect in CA cells (Fig. S20D). These results position CD44 as a key node in CA-induced stress tolerance, particularly under conditions of UPR suppression.

Since HA–CD44 interactions can activate diverse downstream pathways depending on receptor interactions^47^, and CD44 has been implicated in MAPK signaling^48,51^, we examined phosphorylation of p38 MAPK following HA treatment. CA elevated basal phospho-p38 levels in Panc1 cells, consistent with enhanced stress signaling, whereas exogenous HA supplementation selectively reduced this phosphorylation (Fig. S20E). Functionally, HA treatment negatively affected the proliferation in control (dox–) cells but had no effect in cells with CA (Fig. S20F). Given that p38 is a stress-activated kinase capable of inducing apoptosis in response to cellular damage^54^, these findings indicate that HA–CD44 engagement attenuates p38-driven stress responses. In this context, HA may partially buffer centrosome-amplified cells against the detrimental consequences of stress signaling.

Since CA is associated with aberrant cell divisions and increased chromosomal instability, we next investigated the role of CD44 in maintaining mitotic fidelity under these conditions. Extra centrosomes are linked to multipolar spindle formation in metaphase, but cancer cells typically rely on centrosome clustering mechanisms to prevent lethal multipolar divisions^55,56^ (Fig. 6F). To assess how CD44 influences centrosome clustering, we used two approaches. First, we induced CA in CD44-KO Panc1-PLK4 cells and quantified multipolar spindle formation, observing a significant increase in multipolar metaphases in all three sgRNA transduced groups compared to controls (Fig. 6G). Second, we compared CD44^low^ (bottom 10%) and CD44^high^ (top 10%) sorted cells. After 7 days of CA, multipolar spindles were more frequent in CD44^low^ cells (∼50%) than in CD44^high^ (∼30%) or unsorted populations (∼30%) (Fig. 6H). We further tested this in Mia Paca-2-PLK4 cells, which showed a weaker CD44 increase upon CA compared to Panc1 (Fig. 6A, S19B). Nevertheless, CD44^low^ Mia Paca-2 cells still exhibited increased multipolar spindle formation, mirroring the pattern observed in Panc1 (Fig. 6I).

Since centrosome amplification is associated with aggressive, drug-resistant cancers and poor patient outcomes^8,10^, we next evaluated the prognostic value of the HA-CD44 axis in TCGA patient samples. We first compared overall survival in patients stratified by PLK4, CD44, or HMMR expression levels. While PLK4 or HMMR levels alone did not significantly affect survival, CD44 expression levels were associated with a marked reduction in overall survival (Fig. S21A-C). We then examined the combined effects and found that patients with both PLK4^high^/CD44^high^ expression had significantly worse survival compared to PLK4^high^/CD44^low^ (p = 0.0287) and PLK4^low^/CD44^low^ patients (p = 0.0116). No significant effect was observed for the combined effects of PLK4 and HMMR (Fig. S21D-E). These findings highlight CD44 as a key modifier of the poor prognosis associated with centrosome amplification.

Together, these results highlight CD44 as a critical regulator of centrosome clustering and a potential vulnerability in centrosome-amplified PDAC cells. Importantly, our data indicate that this role extends beyond spindle mechanics: HA–CD44 signaling simultaneously reduces p38-mediated stress responses and cooperates with UPR pathways to buffer proteotoxic stress, as reflected by the increased sensitivity of CD44-KO cells to TUDCA-induced UPR inhibition. Thus, CD44 safeguards centrosome-amplified cells by integrating centrosome clustering with stress adaptation mechanisms.

## 3 Discussion

Centrosome amplification (CA) is a hallmark of many cancers^5^, including pancreatic ductal adenocarcinoma (PDAC), where it is associated with genomic instability and poor outcomes^7,8^. While CA promotes tumor evolution, it also imposes significant stress that must be managed for cell survival^55,56^. Our study reveals that PDAC cells with CA adopt distinct metabolic adaptations, creating specific, targetable dependencies in redox homeostasis, unfolded protein response (UPR) signaling, and hyaluronic acid (HA) synthesis (Fig. 7). These adaptations are not correlative but essential, as their disruption selectively impairs the survival of cells with CA.

**Figure 7.**
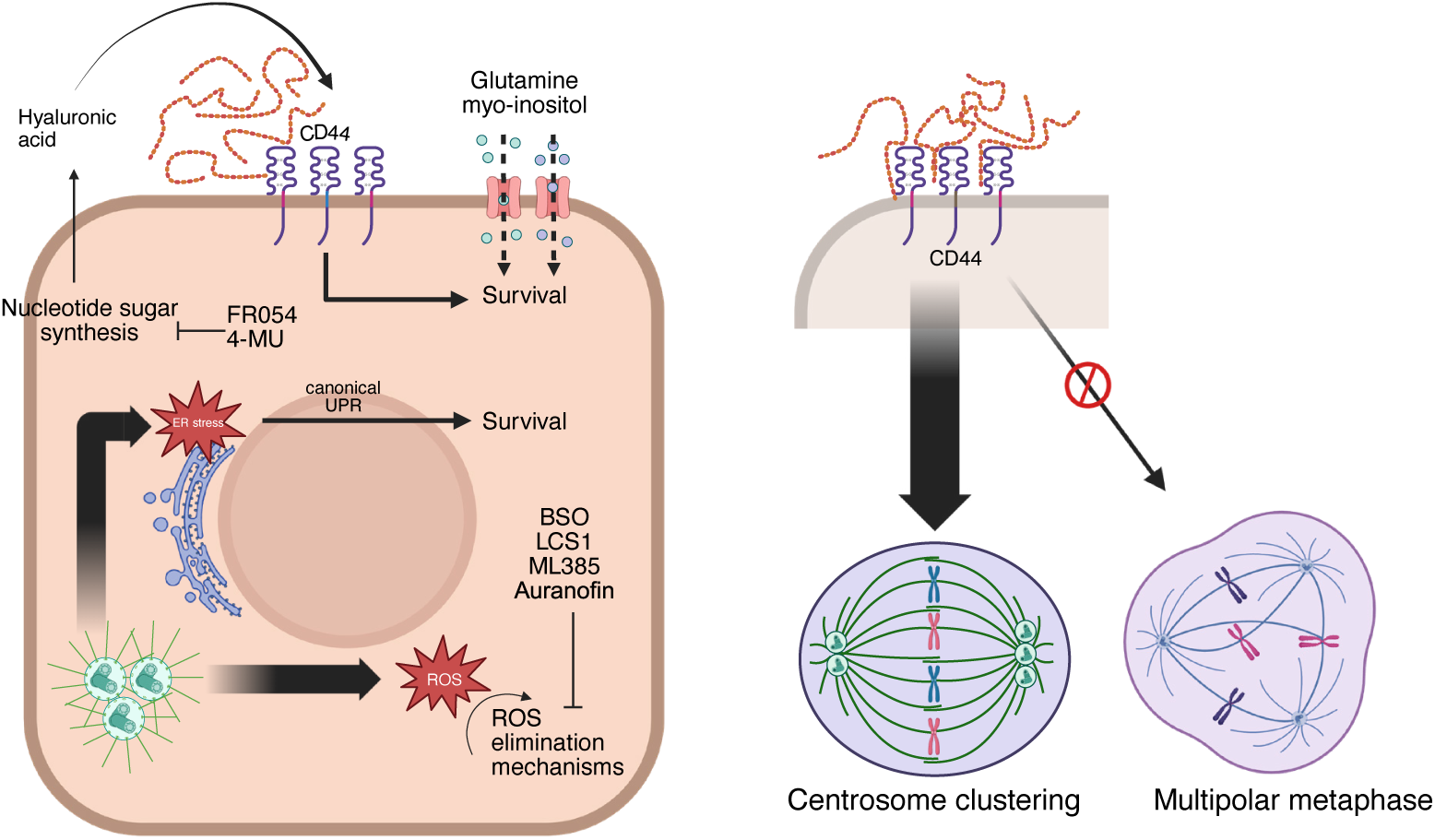
Proposed model of metabolic dependencies in PDAC cells with PLK4-induced centrosome amplification. CA increases cellular vulnerability to inhibition of ROS detoxification pathways and to suppression of both the glucuronic acid and hexosamine biosynthetic pathways (HBP). These cells also exhibit increased dependence on hyaluronic acid synthesis, extracellular glutamine and inositol availability. Moreover, disruption of HA–CD44 signaling impairs centrosome clustering, thereby compromising mitotic fidelity. Created in BioRender. Ozcan, S. (2026) https://BioRender.com/hrtnym8

A primary consequence of CA is increased intracellular reactive oxygen species (ROS)^14^, which we confirmed across PDAC models. This oxidative stress creates a strong reliance on antioxidant defenses. CA cells were highly sensitive to inhibition of glutaminase (GLS1), the rate-limiting enzyme in glutamine catabolism, consistent with the role of glutamine as a precursor for glutathione (GSH) synthesis^20^. Inhibiting GSH synthesis with BSO was selectively lethal to CA cells. Because GSH depletion is a canonical trigger of ferroptosis, we tested whether ferroptotic death contributed to this phenotype. Ferrostatin-1, a potent ferroptosis inhibitor, did not rescue CA cells, suggesting that classical ferroptosis is not the dominant death mechanism. Nonetheless, given the central role of GSH in ferroptotic regulation and the variability of ferroptosis across tumors, we cannot exclude its contribution under conditions not captured here.

Our metabolism-focused CRISPR screen and subsequent validation experiments further pinpointed antioxidant dependencies, identifying essential roles for the thioredoxin system (e.g., TXNRD2), superoxide dismutases (SOD1/2/3), and peroxiredoxins (PRDX1) in CA cell survival. This is consistent with glutamine-derived glutamate fueling GSH synthesis, supporting redox balance beyond anaplerosis^21^. Dihydrofolate reductase (DHFR), a folate cycle enzyme, also emerged as a novel redox vulnerability: its inhibition increased ROS and impaired survival, suggesting a potential role in redox regulation via tetrahydrobiopterin and nitric oxide synthase activity^37^. Together, these results position CA cells as reliant on a multilayered antioxidant network, largely fueled by glutamine metabolism, to tolerate oxidative stress. In parallel, CA was associated with activation of NRF2 signaling, as shown by NRF2 nuclear translocation and increased ARE reporter activity, consistent with prior reports^14^. The differential sensitivities observed, greater GSH dependence in Panc1 and a stronger reliance on NRF2 signaling in MiaPaCa-2, likely reflect likely reflect the redundancy of antioxidant systems and the genetic context of PDAC models^57^.

Alongside redox stress, CA imposed a strong reliance on protein quality-control associated pathways. The depleted hits in CRISPR screen revealed enrichment for genes in the hexosamine biosynthesis and uronic acid pathways, which generate UDP-sugars for glycosylation and glycosaminoglycan synthesis. Functional validations confirmed that inhibition of HA synthesis (4-MU), HBP flux (FR054, targeting PGM3), or N-linked glycosylation (tunicamycin) was selectively toxic to cells with CA, whereas O-linked glycosylation, though important for centrosome-regulated polarity^58^, was dispensable. Mechanistically, CA triggered robust activation of all three UPR branches, PERK, IRE1α, and ATF6, indicating elevated proteostatic stress. This activation was adaptive: both hyper-activation and suppression (via chemical chaperones TUDC and 4-PBA) were detrimental, suggesting that CA cells maintain a precarious “hyper-equilibrium” of UPR signaling^41^. Together, these observations raise the important question of how distinct stress-responsive transcription factors, including NRF2, ATF4, and ATF6, differentially contribute to the survival of centrosome-amplified cells. Determining whether these pathways function redundantly or in a context-dependent manner to buffer oxidative and proteostatic stress will be an important direction for future investigation.

The reliance on hexosamine and uronic acid metabolism is consistent with previous findings that the HBP constitutes a metabolic vulnerability in cancers such as lung^59^ and breast^60^. However, in our experiments, this vulnerability was not a general feature of PDAC cells, but rather emerged specifically in the context of CA-induced stress, indicating that HBP and uronic acid pathway dependence are tightly linked to this altered cellular state. This is reminiscent of a recently described conditional essentiality in sugar nucleotide metabolism, where cells with high UGDH expression become dependent on UXS1 to detoxify accumulated UDP-glucuronic acid and maintain Golgi homeostasis^61^. In our system, supplementation with UDP-glucuronic acid—the metabolite produced by UGDH—partially rescued viability, suggesting that CA cells may experience increased utilization of this key metabolite, creating a dependency on its production. Conversely, the failure of N-acetyl glucosamine (GlcNAc) to provide a survival advantage points to a bottleneck in the salvage pathway, potentially at the level of N-acetylglucosamine kinase (NAGK) activity or transport. Collectively, our findings suggest that centrosome amplification creates a unique metabolic state characterized by an increased demand for specific sugar nucleotides, unveiling a targetable liability that could be exploited therapeutically.

The identification of SLC5A3 as a selective dependency of centrosome-amplified cells, together with the ability of myo-inositol or D-glucuronate supplementation to rescue their survival, indicates that CA creates an increased requirement for extracellular inositol. This may arise from impaired *de novo* biosynthesis, as suggested by IMPA1 downregulation in Mia PaCa-2 cells, or from functional bottlenecks in inositol metabolism or flux in other models. Because inositol-derived phospholipids play central roles in ER membrane composition, vesicular trafficking, and protein quality control^62^, increased reliance on inositol uptake may reflect an adaptive response to the proteotoxic and membrane stress imposed by centrosome amplification. In this context, SLC5A3-mediated inositol import may act as a metabolic buffer that supports ER function and proteostasis under chronic CA-induced stress.

Additionally, our work highlights the importance of CA, chromosomal instability–associated division abnormalities, and related genetic backgrounds in shaping metabolic programs. For instance, previous studies have shown that LKB1/KRAS mutant lung adenocarcinoma cells display elevated flux through the hexosamine biosynthesis pathway (HBP) and increased dependence on GFPT2^59^. Given the established role of LKB1 (STK11) in centrosome biology^63^, as well as in regulating the chromosomal passenger complex (CPC) and maintaining genome stability^64^, it would be intriguing to investigate whether CA contributes to the enhanced HBP flux observed in LKB1/KRAS mutant cancers. Moreover, a recent aneuploidy-focused metabolic CRISPR screen identified DHODH as a top dependency in aneuploid cells^65^, highlighting how stable aneuploidy creates distinct metabolic vulnerabilities. By contrast, our findings suggest that CA—potentially together with chromosomal instability—drives reliance on ROS detoxification and nucleotide sugar/glycan biosynthesis. Together, these insights point to the value of integrating genomic instability with metabolic profiling to reveal novel, context-specific therapeutic vulnerabilities across cancers.

A central discovery of our study is that centrosome amplification (CA) co-opts hyaluronic acid (HA) metabolism to sustain the mitotic fidelity. We find that CA up-regulates both HA synthesis and the expression of its receptor, CD44, creating a dependency on this ligand-receptor axis. Functionally, we demonstrate that the HA-CD44 system is required for two critical and distinct processes: (i) centrosome clustering to suppress multipolar divisions, and (ii) successful cytokinesis to prevent multinucleation. This reveals a dual mechanism through which extracellular matrix remodeling safeguards cell division under the profound stress of CA. We propose that the HA-CD44 axis facilitates this by orchestrating cytoskeletal organization and force distribution, a role consistent with CD44-mediated mechanotransduction^48^. This is supported by our observation that CA elevates pro-apoptotic p38-MAPK phosphorylation and that exogenous HA attenuates this signal. Furthermore, HA supplementation partially rescues the cytokinesis defects induced by UGDH knockout, confirming its active role as a regulatory factor, not just a structural scaffold. Notably, the HA-CD44 pathway becomes essential when proteostatic buffering is compromised, as reducing UPR activity with TUDC selectively targets CA cells in a CD44-dependent manner. This suggests that, under extreme stress, cancer cells rely on this mechanical signaling network as a critical compensatory survival mechanism. Integrating our findings with the established role of RHAMM as a mitotic spindle regulator^66,67^, we postulate the existence of an HA-CD44-RHAMM network that synchronizes extracellular cues with intracellular machinery to monitor and execute accurate cell division. The strong correlation between RHAMM and PLK4 expression in PDAC patient tumors further underscores the clinical relevance of this mechanism. Thus, we define a critical metabolic-physical circuit that maintains viability under the proteotoxic and mitotic stress generated by centrosome amplification, revealing a new vulnerability in aggressive cancers.

Notably, our findings align with and extend the concept of an extra-centrosome associated secretory phenotype (ECASP)^13,14^. While a prior study demonstrated that CA drives the secretion of pro-inflammatory cytokines and growth factors, such as IL-8, with the potential to remodel the tumor microenvironment, our study reveals an additional dimension to this phenotype: the profound metabolic reprogramming required to sustain both cell-autonomous survival and secretory capacity. The increased dependency on hexosamine and uronic acid pathways that we identified likely supports not only intracellular stress management but also the extensive glycosylation requirements for secreted factors that characterize ECASP. This connection suggests that CA cells must tightly coordinate their metabolic and secretory activities to thrive in challenging tumor environments.

While our study reveals profound metabolic dependencies in PDAC cells with CA, a limitation is the absence of comprehensive metabolomic profiling to determine whether these vulnerabilities reflect broader metabolic rewiring. Although we identify discrete dependencies in HA synthesis, UPR signaling, and redox homeostasis, these processes are highly interconnected within cellular metabolism. CA may therefore induce metabolic adaptations that extend beyond the pathways directly tested here. For example, increased demand for nucleotide sugars in HA synthesis could alter flux through the hexosamine biosynthesis pathway, with consequences for both N-glycosylation and glycosaminoglycan synthesis. Indeed, CA cells were selectively sensitive to inhibition of N-glycosylation, consistent with an increased need for protein processing and extracellular matrix remodeling to tolerate stress. Likewise, redox stress and UPR activation may reshape amino acid and nucleotide metabolism, with alterations in glutathione and thioredoxin systems affecting redox-sensitive signaling pathways beyond oxidative damage control. Our findings further suggest that these adaptations are dynamic: under reduced UPR signaling, CA cells become increasingly reliant on the HA–CD44 axis, implying that extracellular matrix–driven mechanotransduction can substitute for diminished proteostasis in maintaining mitotic fidelity and survival. Definitive assessment of such global changes will require metabolomics and flux-based analyses, such as stable isotope tracing with ^13^C-glucose or ^15^N-glutamine, to map carbon and nitrogen flow, uncover compensatory pathways, and clarify whether the dependencies we observe represent absolute requirements or instead bottlenecks within a reorganized metabolic network.

A limitation of our study is that centrosome amplification was induced exclusively through PLK4 overexpression. Although PLK4 overexpression is a widely used and effective approach to generate supernumerary centrosomes, PLK4 has also been implicated in regulating cytoskeletal organization and signaling pathways that are intimately connected to centrosome amplification, chromosomal instability, and mitotic errors^68^. This interdependence makes it difficult to assign strict causality to centrosome number alone in PLK4 overexpression models. Accordingly, the metabolic and stress-response dependencies identified here should be interpreted as vulnerabilities associated with the PLK4-induced CA state. Validation of these dependencies using orthogonal approaches to induce centrosome amplification, such as cytokinesis failure or centriole disengagement defects, will be necessary to more fully disentangle centrosome-driven effects from potential kinase-dependent contributions of PLK4.

Taken together, our results reveal that CA drives non-redundant metabolic dependencies in redox control, proteostasis, and glycosaminoglycan synthesis. These pathways converge on the HA–CD44 signaling axis, which safeguards centrosome clustering and mitotic fidelity, integrating metabolic and structural adaptations that ensure the survival of genomically unstable CA cells. Importantly, these vulnerabilities are targetable: GLS1 inhibitors, UPR modulators, and HA synthesis blockade each selectively impaired CA cell fitness, and combined perturbations showed enhanced lethality. Since CA marks aggressive and treatment-resistant PDAC, therapeutic strategies that disrupt these adaptations could selectively eliminate the most dangerous tumor cell populations.

## 4 Methods

### Cell culture

Human pancreatic ductal adenocarcinoma cell lines Panc1 (CRL-1469), MiaPaCa-2 (CRL-1420), BxPC-3 (CRL-1687), and U2OS osteosarcoma cell line (HTB-96) were obtained from ATCC. All cell lines were tested monthly for mycoplasma contamination. Cells were maintained in DMEM (Sigma, D6429) supplemented with 10% tetracycline-free FBS (biowest, S181T) and 1% penicillin-streptomycin at 37°C in 5% CO2. In glutamine depletion experiments, a DMEM without L-gln and Na-pyr was used (Sigma, D5671). Centrosome amplification was induced with 2 µg/mL doxycycline for the indicated durations. Detailed information about the chemicals and inhibitors used in the study is provided in Supplementary tables 1 and 2.

### Plasmids and lentivirus generation

Doxycycline-inducible cell lines were generated by lentiviral transduction with pCW57-PLK4^18^ at an MOI (multiplicity of infection) of 5, followed by hygromycin selection (200 µg/mL). For competition assays, lentiviral H2B-GFP (Addgene, 21210) and H2B-mCherry (Addgene, 21217) expression plasmids were used. For CRISPR knockout, sgRNAs targeting UGDH and CD44 (gRNA sequences are provided in Supplementary table 3) were cloned into lentiCRISPR-v2 (Addgene, 52961). Lentiviral particles were produced in HEK293T cells using psPAX2 (Addgene, 12260) and VSV.G (Addgene, 14888) packaging plasmids. Target cells were infected at an MOI of 2 in the presence of 8 µg/mL polybrene and selected with the appropriate antibiotics. For ARE-reporter assays, pREP-8xARE-GFP-SV40-BFP (Addgene, 134910) plasmid was transfected to cells with Lipofectamine 3000 (Thermo Fisher, L3000015).

### Metabolic enzyme targeted CRISPR screen

The metabolism-focused CRISPR knockout library^69^, targeting 2,981 metabolic genes with 10 sgRNAs per gene (29,790 sgRNAs total), was obtained from Addgene (110066) and amplified following published protocols^70^. Library quality was confirmed by next-generation sequencing to ensure uniform sgRNA representation prior to lentiviral production. Viral titers were determined in Panc1 cells, and infections were carried out at a multiplicity of infection (MOI) of 0.6 to favor single-copy integration while maintaining 300x coverage. After puromycin selection, baseline (day 0) samples were collected, and cells were split into doxycycline-treated and untreated groups for 21 days of culture. At each passage, cell numbers were monitored to preserve library representation and minimize dropout. CRISPR screening experiments were conducted in two independent repeats. Genomic DNA was isolated (Macherey-Nagel, NucleoSpin Tissue, 740952), sgRNA cassettes were PCR-amplified, and sequencing was performed on an Illumina NovaSeq 6000 platform.

### CRISPR screen data analysis

Sequencing reads were processed and analyzed using MAGeCK^71^, with normalization performed at the sgRNA level. Gene essentiality was quantified by *β* scores, and differential essentiality was defined as *β*_dox+_ − *β*_dox-_. MAGeCK-Flute was used for cell cycle normalization of *beta* scores and for the visualization of MAGeCK-MLE analysis^72^. To generate a top depleted gene list, a depletion threshold was established by calculating the median of all differential *β* scores and subtracting twice the median absolute deviation (MAD) from this median. Genes with differential *β* scores falling below this threshold were considered significantly depleted. Additionally, *Waldp* values from the MAGeCK MLE analysis were used in downstream analyzes (Fig. 3E). Gene Set Enrichment Analysis (GSEA) was carried out using MSigDB gene sets, and protein-protein interaction networks were constructed with STRINGdb (https://string-db.org) and clustered using the Markov Clustering Algorithm (MCL), with visualization in Cytoscape (v.3.10.3). Gene Ontology annotations were obtained from QuickGO (https://www.ebi.ac.uk/QuickGO/), with corresponding datasets filtered for human genes and used to analyze our screening results. GO-BP, MSigDB, KEGG, and TR-RUST analyzes were performed using ShinyGO (v.0.82, https://bioinformatics.sdstate.edu/go/).

### TCGA data analysis

TCGA pancreatic cancer dataset was accessed from the NIH–National Cancer Institute (NCI) web portal (https://portal.gdc.cancer.gov/projects/TCGA-PAAD; last accessed 04/30/2025). Patient samples were filtered to include only adenocarcinoma cases (*n* = 82). Gene Expression Clustering analysis was performed, applying unsupervised Euclidean clustering across both genes and patients. Gene lists were restricted to top CRISPR screen hits as well as the CA20 and CIN25 gene signatures. All genes included in the analysis are presented in Supplementary Fig. 7A. Uniform Manifold Approximation and Projection (UMAP) clustering was applied to the downloaded dataset using the umap package (v.0.2.10) in R (v.4.5.1)^73^. Signature scores for chromosomal instability (CIN25) and centrosome amplification (CA20) were calculated as the mean Z-score normalized expression of their respective gene sets. Z-score normalization was performed across samples prior to score calculation. Gene expression correlations in the TCGA dataset were assessed using cBioPortal (https://www.cbioportal.org). Additionally, Gene Expression-based Network Inference (GENI) analysis was conducted via the online platform (https://www.shaullab.com/geni).

### Flow cytometry assays

For intracellular ROS measurement, cells were stained with 10 µM H_2_DCFDA (Thermo Fisher, D399) for 20 minutes at 37°C. For cell surface hyaluronic acid detection, cells were stained with biotinylated HA-binding peptide (Anaspec, AS-65199), followed by Streptavidin-AlexaFluor-568 (Thermo Fisher, S11226), in accordance with a previously published method^74^. CD44 surface levels were measured using PE-conjugated anti-CD44 antibody (BioLegend, 103007). For cell cycle distribution analysis, cells were fixed in 70% ethanol, treated with RNase A (Thermo Fisher, EN0531), and stained with propidium iodide. In all assays, FSC/SSC populations were gated to identify live cell populations, and single cells were selected using SSC-H/SSC-A gating strategies. All analyses were performed on single-cell gated populations. For ARE-activation experiments, BFP-positive cells were first gated, and GFP intensity in this subpopulation was measured and compared. Single fluorescent expressing cells were used for compensation. Flow cytometry analyses were performed using either an Attune NxT flow cytometer (Thermo Fisher) or a BD CytoFLEX instrument (BD Biosciences). All flow cytometry data were analyzed and visualized using FlowJo software (v10.8.1).

### Dual color competition assays

H2B-GFP and H2B-mCherry labeled cells were mixed at 1:1 ratio and plated in 6-well plates. Cells were treated with indicated compounds and doxycycline, with media and compounds refreshed every two days. At indicated time points, cells were trypsinized and analyzed by flow cytometry to determine GFP/mCherry ratios. Relative depletion was calculated as log_2_(%mCherry_dox+_ / %mCherry_dox-_). Representative flow cytometry plots showing non-normalized %mCherry and %GFP levels were presented in supplementary figures.

### Glutathione assays

Intracellular reduced and oxidized glutathione levels were quantified using the GSH/GSSG-Glo assay (Promega, V6611) according to the manufacturer’s instructions. Control and centrosome-amplified (5d) cells were seeded in white, clear-bottom 96-well plates and allowed to adhere overnight. Cells were lysed and bioluminescent signals corresponding to total and oxidized (GSSG) glutathione were measured using a plate reader. GSH/GSSG ratios were calculated for each condition from the respective luminescence values. To correct for differences in cell number, signals were normalized to total protein content determined from parallel wells using a BCA assay.

### Fluorescence activated cell sorting

For CD44 high/low and CD44-KO experiments, cells were stained with PE-conjugated anti-CD44 antibody (BioLegend, 103007) and sorted on a cell sorter (Sony, SH800S). The top and bottom 10% of CD44-expressing populations were collected for downstream assays. All fluorescence-activated cell sorting (FACS) procedures were carried out using the ultra-purity sorting mode. The sorted populations are displayed in the corresponding plots within the manuscript figures.

### Immunofluorescence and confocal microscopy

Cells grown on glass coverslips were fixed in methanol at −20^◦^C for 10 minutes, washed three times with PBS, and blocked with 5% BSA in PBS. Cells were then stained with anti–*γ*-tubulin antibody (Sigma, T6557; 1:500 dilution), followed by Alexa Fluor 488–conjugated secondary antibody (Invitrogen, A-11001; 1:500 dilution). DNA was counterstained with DAPI, and coverslips were mounted on glass slides. Images were acquired using either a Leica DMI8 widefield microscope or a Nikon AXR confocal microscope. Maximum intensity projections of Z-stacks were generated. Images were analyzed using ImageJ and QuPath. For quantification of cell and nuclear size, cell boundaries were identified using thresholding of the cytoplasmic signal, and nuclei were segmented by automated thresholding of the DAPI channel (Otsu method).

### Western blotting and antibodies

Cells were lysed in RIPA buffer with protease and phosphatase inhibitors. Nuclear and cytoplasmic fractions were separated using a commercial kit (Thermo Fisher, 78833). Proteins were separated by SDS-PAGE, transferred to PVDF membranes, and probed with antibodies: FLAG (Sigma, F1804, 1:1000), GAPDH (Cell Signaling, 2118, 1:2000), Histone H3 (Cell Signaling, 4499, 1:2000), NRF2 (Cell Signaling, 12721, 1:1000), ATF4 (Cell Signaling, 11815, 1:1000), ATF6 (Cell Signaling, 65880, 1:1000), IRE1α (Cell Signaling, 3294, 1:1000), phospho-p38 (Cell Signaling, 4511, 1:1000). Blots were developed with ECL reagent (Luminata Forte, Millipore, WBLUF0020) and imaged on the ChemiDoc system (Bio-Rad).

### RT-qPCR and RT-PCR

Total RNA was extracted using the NucleoSpin RNA kit (Macherey-Nagel; 740955), and cDNA was synthesized from 1 µg RNA with M-MLV reverse transcriptase (Invitrogen; 28025013). For RT-qPCR, 10 ng of cDNA was amplified with SYBR Green Master Mix (Roche; 04707516001), using GAPDH as the endogenous control. Primer sequences are listed in Supplementary Table 4. For CD44 splicing analysis by RT-PCR, 2 µl of synthesized cDNA was used as template. The primers were: forward 5^′^-AGTCACAGACCTGCCCAATGCCTTT-3^′^ and reverse 5^′^-TTTGCTCCACCTTCTTGACTCCCATG-3^′^.

### Cell viability and colony formation assays

Cell viability in 96-well plates was assessed using sulforhodamine B (SRB) staining. For viability assays in 6-well plates, cell numbers were quantified using an automated cell counter (BioRad) with the trypan blue exclusion method. For colony formation assays, 300 cells (Panc1) or 500 cells (Mia Paca-2 and BxPC-3) were seeded per well in standard 6-well plates and cultured under the indicated conditions. Colonies were fixed with methanol, stained with crystal violet, and imaged. Stain was then solubilized in acetic acid (3%) and quantified using a microplate reader at 590 nm absorbance.

### Statistical analyses

All experiments were performed with multiple independent biological replicates, and independent repeats were shown in the related plots. Data are presented as mean ± SD, unless otherwise stated. Statistical analyzes were performed using GraphPad Prism 9 and R. For comparisons between two groups, unpaired two-tailed Student’s t-test was used. For multiple group comparisons, a one-way ANOVA with appropriate post-hoc tests was applied. For competition assays, two-way ANOVA with multiple comparisons was used. A p-value of < 0.05 was considered statistically significant.

## 5 Acknowledgements

The authors gratefully acknowledge the use of the services and facilities of the Koç University Research Center for Translational Medicine (KUTTAM), funded by the Presidency of Turkey, Presidency of Strategy and Budget.

## 6 Author contributions statement

Conceptualization: SCO, Investigation: SCO, EG, BMK, EC, and BK, Methodology: SCO, EG, Visualization: SCO, Project Administration: SCO, CAA, Funding Acquisition: SCO, Writing - original draft: SCO, Writing - review & editing: CA.

## 7 Additional information

This research was funded by TUSEB (23066, SCO) and in part by TUBITAK (120Z830, SCO). The funders had no role in study design, data collection and analysis, the decision to publish, or preparation of the manuscript.

## Data availability

The raw and processed sequencing data from the CRISPR screen are available as Data Table 1. All other data are available from the corresponding authors (S.C.O. and C.A.A.) upon request.

## 8 Supplementary Tables

**Table 1.** Supplementary table - 1: Inhibitors used in this study.

| Compound | Target | Vendor | Catalog Number | Concentration Range |
| --- | --- | --- | --- | --- |
| CB-839 | GLS1 inhibitor | MedChemExpress | HY-12248 | 10-20 μM |
| BPTES | GLS1 inhibitor | MedChemExpress | HY-12683 | 10-20 μM |
| BSO | GCL inhibitor | MedChemExpress | HY-106376A | 100-200 μM |
| EGCG | GLUD1/2 inhibitor | MedChemExpress | HY-13653 | 20-40 μM |
| ML385 | NRF2 inhibitor | MedChemExpress | HY-100523 | 5-10 μM |
| ML334 | Keap1-NRF2 disruptor | MedChemExpress | HY-110258 | 5-10 μM |
| meAIB | SNAT1 inhibitor | MedChemExpress | HY-134452 | 5 mM |
| Ferrostatin-1 | Ferroptosis inhibitor | MedChemExpress | HY-100579 | 2 μM |
| LCS-1 | SOD1 inhibitor | MedChemExpress | HY-115445 | 0.5 μM |
| Auranofin | TXNRD inhibitor | MedChemExpress | HY-B1123 | 0.5 μM |
| Pralatrexate | DHFR inhibitor | TargetMol | T6120 | 1-2 nM |
| Azaserine | GFPT inhibitor | MedChemExpress | HY-B0919 | 5-10 μM |
| FR054 | PGM3 inhibitor | TargetMol | T9468 | 100-200 μM |
| 4-MU | HA synthesis inhibitor | TargetMol | T1391 | 500-750 μM |
| OSMI-1 | OGT inhibitor | MedChemExpress | HY-119738 | 20 μM |
| Tunicamycin | N-glycosylation inhibitor | TargetMol | T13229 | 0.25-1 μg/mL |
| Thapsigargin | SERCA inhibitor | TargetMol | TQ0302 | 10 nM |
| Toyocamycin | XBP1 splicing inhibitor | TargetMol | T17143 | 10 nM |
| GSK2656157 | PERK inhibitor | TargetMol | T2654 | 10 μM |
| TUDCA | Chemical chaperone | TargetMol | T2532 | 200-400 μM |
| 4-PBA | Chemical chaperone | TargetMol | T5886 | 200-400 μM |

**Table 2.** Supplementary table - 2: Chemical compounds used in this study.

| Compound | Vendor | Catalog Number | Concentration Range |
| --- | --- | --- | --- |
| Doxycycline | MedChemExpress | HY-N0565B | 1 μg/mL |
| L-Glutamine | MedChemExpress | HY-N0390 | 2-4 mM |
| Sodium pyruvate | MedChemExpress | HY-Y0810 | 1-2 mM |
| L-glutamate | MedChemExpress | HY-W337739 | 2-4 mM |
| Oxaloacetate | MedChemExpress | HY-W010382 | 2-4 mM |
| UDP-GlcNAc Disodium Salt | TargetMol | T19596 | 50-100 μM |
| UDP-GalNAc | MedChemExpress | HY-114365 | 50-100 μM |
| UDP-glucuronic acid trisodium | TargetMol | T19595 | 50-100 μM |
| D-Glucosamine sulphate | MedChemExpress | HY-N0487 | 125-250 μM |
| N-Acetyl-D-Glucosamine | TargetMol | T4514 | 50 μM |
| i-inositol | TargetMol | T0421 | 50-100 μM |
| D-Glucuronic acid sodium salt monohydrate | TargetMol | T5068 | 50-100 μM |
| Hyaluronic acid sodium (MW 20 kDa) | TargetMol | T88852 | 10-15 μM |
| Hyaluronic acid sodium (MW 40 kDa) | TargetMol | T88854 | 10-15 μM |

**Table 3.**
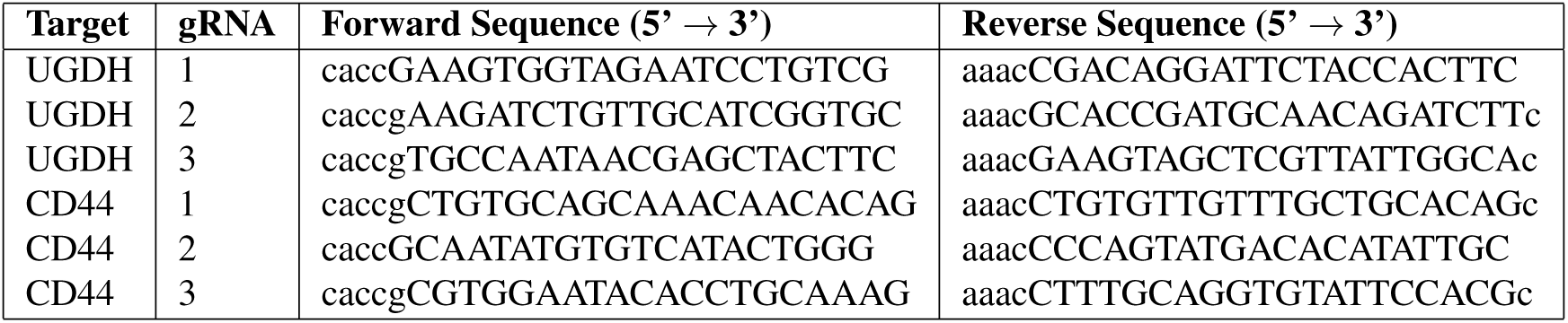
Supplementary table - 3: guide-RNA sequences used in this study.

**Table 4.**
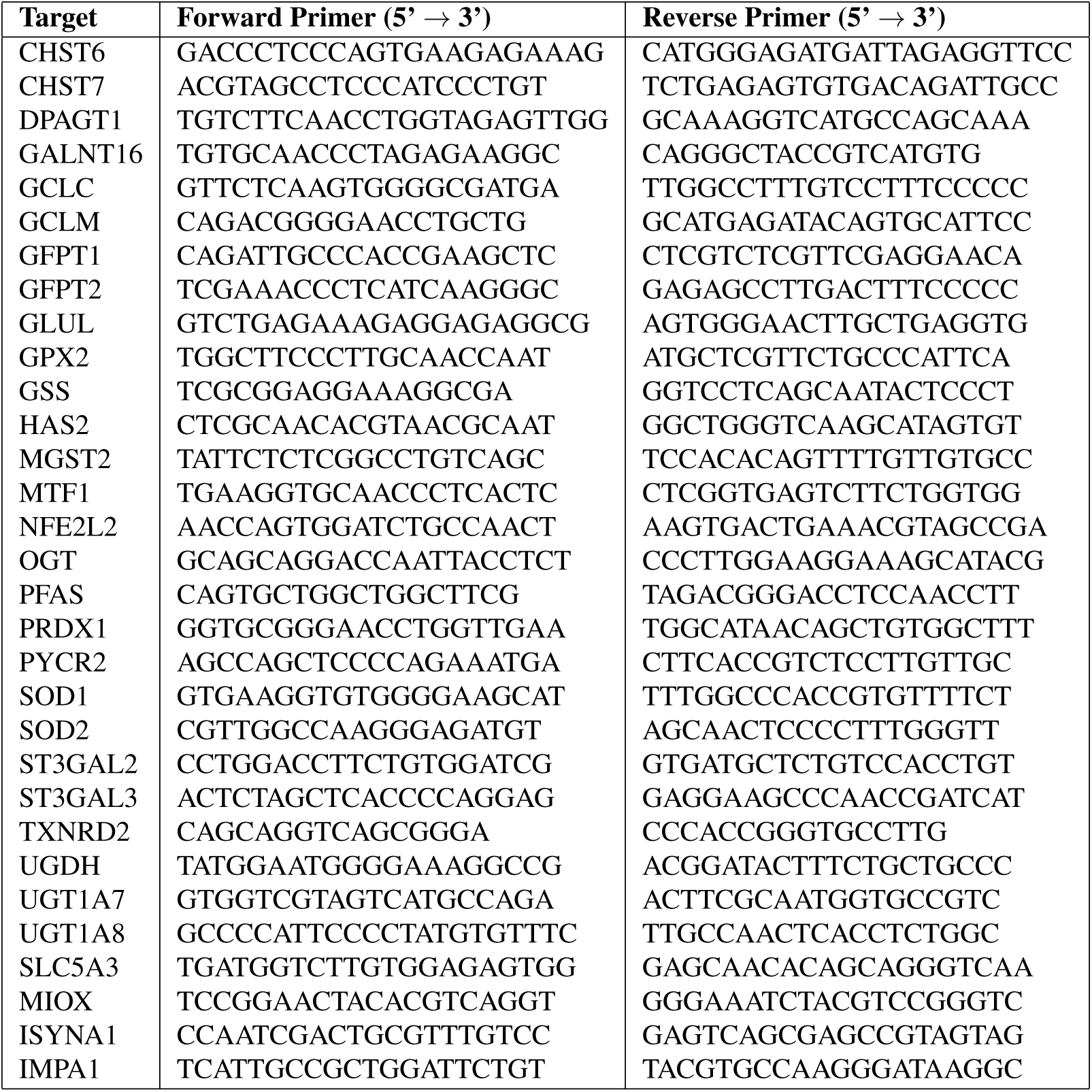
Supplementary table - 4: RT-qPCR primer sequences used in this study.

