## Supplemental Figures and Legends - 1 for "Stress adaptation pathways and HA–CD44 signaling maintain the survival of pancreatic cancer cells with centrosome amplification"

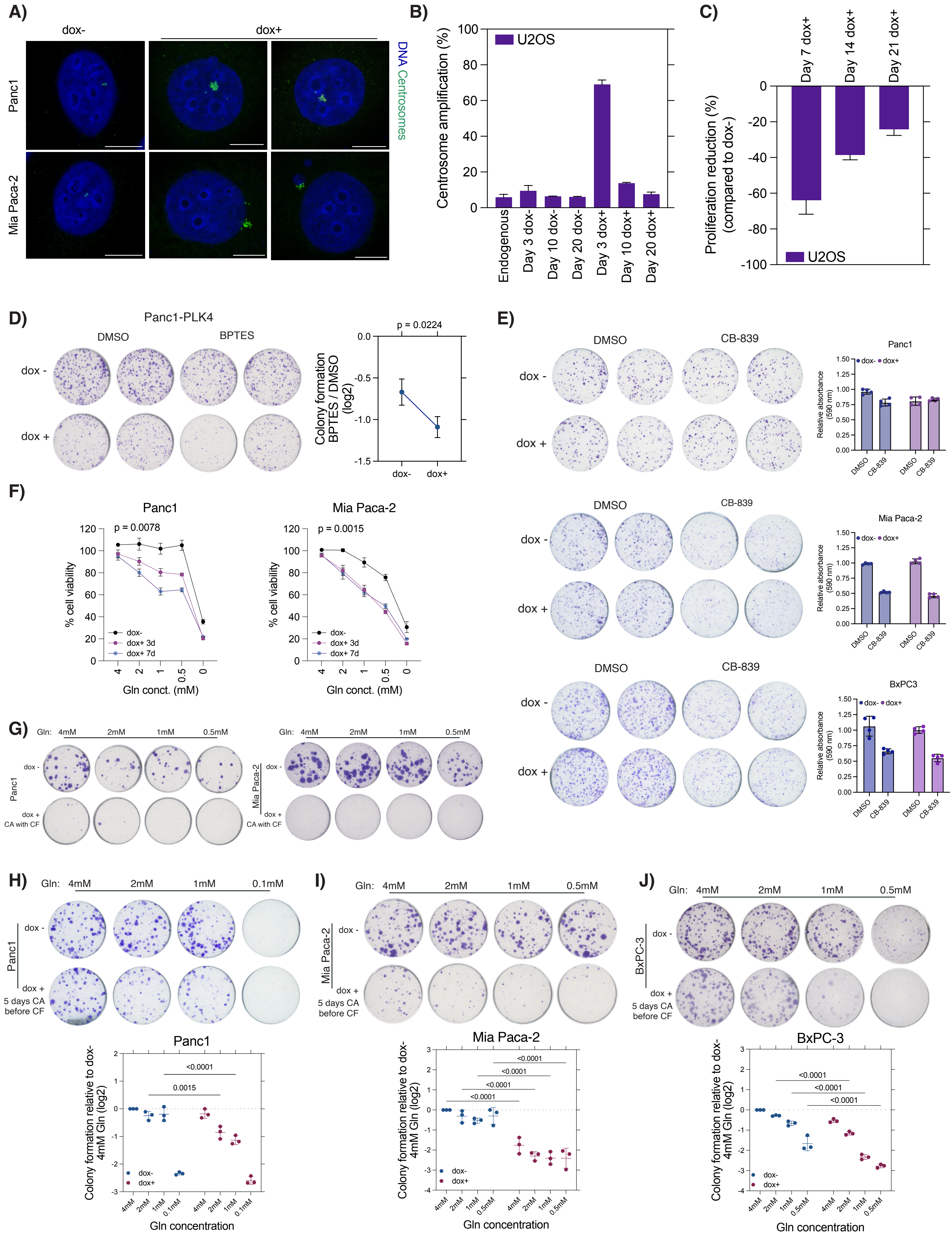

A)
dox-
dox+
Panc1
Mia Paca-2
DNA Centrosomes
B)
U2OS
Centrosome amplification (%)
Endogenous
Day 3 dox-
Day 10 dox-
Day 20 dox-
Day 3 dox+
Day 10 dox+
Day 20 dox+
C)
Day 7 dox+
Day 14 dox+
Day 21 dox+
Proliferation reduction (%) (compared to dox-)
U2OS
D)
Panc1-PLK4
DMSO
BPTES
dox -
dox +
p = 0.0224
Colony formation BPTES / DMSO (log2)
dox-
dox+
E)
DMSO
CB-839
dox -
dox +
Panc1
Mia Paca-2
BxPC3
Relative absorbance (590 nm)
F)
Panc1
Mia Paca-2
p = 0.0078
p = 0.0015
% cell viability
dox-
dox+ 3d
dox+ 7d
Gln conct. (mM)
G)
Gln: 4mM 2mM 1mM 0.5mM
dox +
CA with CF
H)
Gln: 4mM 2mM 1mM 0.1mM
5 days CA before CF
<0.0001
0.0015
Colony formation relative to dox- 4mM Gln (log2)
Gln concentration
I)
Gln: 4mM 2mM 1mM 0.5mM
J)
BxPC-3


Fig S1

**Supplementary Figure 1: Supplemental to Figure 1.**
**A)** Representative images of PLK4 induced centrosome amplification in Panc1 and Mia Paca-2 cells. Centrosomes were stained by γ-tubulin (green) and nucleus was visalized with DAPI (blue).
**B)** PLK4 overexpression in U2OS cells induces robust centrosome amplification in the short term, but the percentage of cells with amplified centrosomes decreases over time.
**C)** Increased centrosome numbers reduce U2OS cell proliferation during the first week following amplification, with gradual recovery in subsequent weeks.
**D)** BPTES treatment reduces colony formation in centrosome-amplified Panc1 cells.
**E)** CB-839 combined with doxycycline does not synergistically impair colony formation in PDAC cells (Panc1, Mia Paca-2, BxPC-3) lacking a doxycycline-inducible PLK4 construct.
**F)** Reduced L-glutamine levels in the culture media impair the viability of PDAC cells with CA. Left panel: Panc1 cells, Right panel: Mia Paca-2 cells.
**G)** Colony formation of PDAC cells with CA induced at the time of seeding is entirely abrogated in the absence of Na-pyruvate, irrespective of L-glutamine concentrations.
**H-J)** PDAC cells with long-term CA exhibit reduced colony formation potential under low L-glutamine conditions. (H) Panc1 cells. (I) Mia Paca-2 cells. (J) BxPC-3 cells. Top panel: Representative colony images from two independent experiments. Bottom panel: Quantification results
Statistical significances were determined by two-tailed t-test in D, by non-linear curve fitting in F, two-way ANOVA in H, I, and J. p values were reported on graphs.

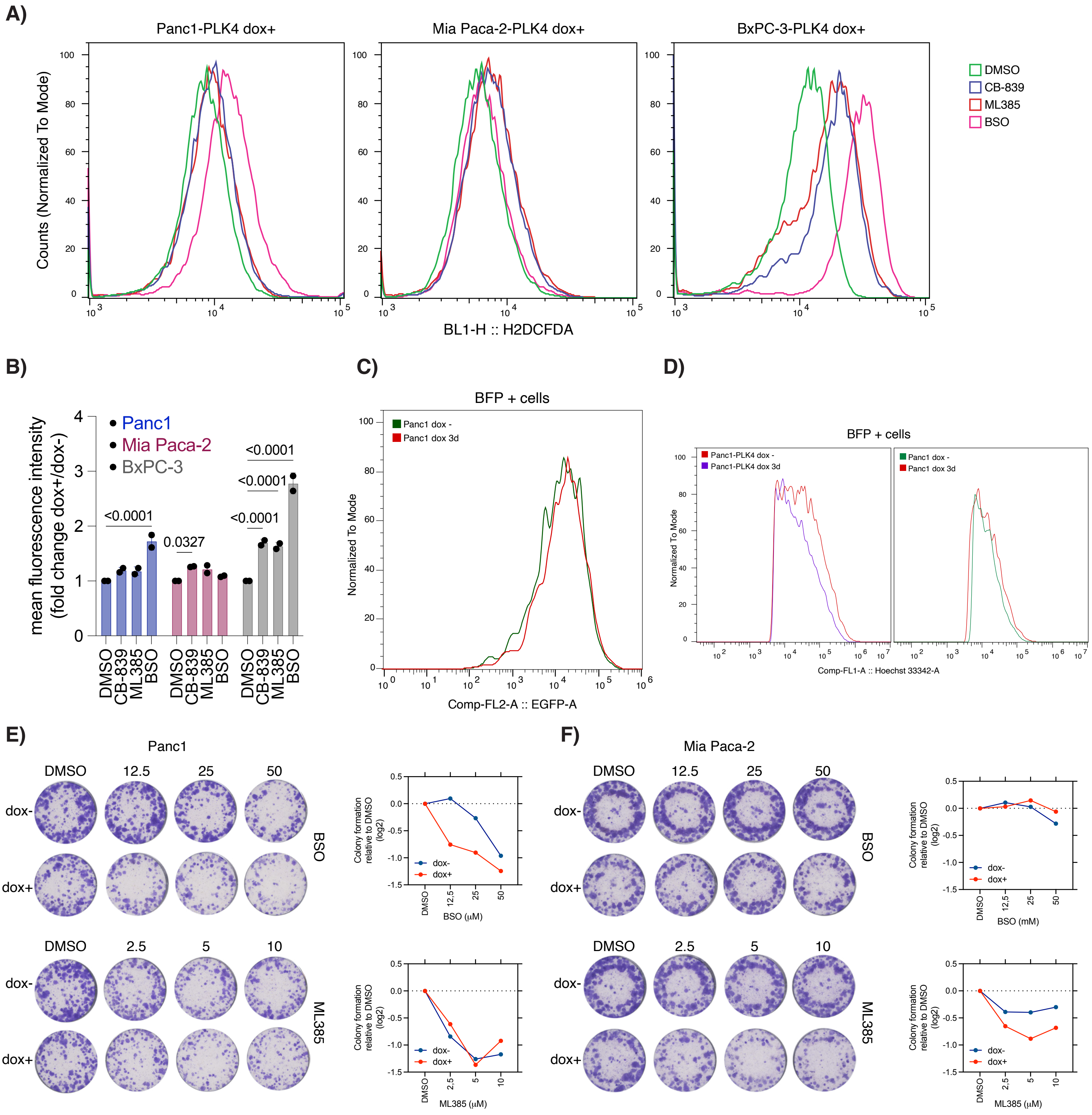
A)
Panc1-PLK4 dox+
Mia Paca-2-PLK4 dox+
BxPC-3-PLK4 dox+
Counts (Normalized To Mode)
BL1-H :: H2DCFDA
DMSO
CB-839
ML385
BSO
B)
mean fluorescence intensity (fold change dox+/dox-)
Panc1
Mia Paca-2
BxPC-3
<0.0001
0.0327
C)
BFP + cells
Panc1 dox -
Panc1 dox 3d
Normalized To Mode
Comp-FL2-A :: EGFP-A
D)
BFP + cells
Panc1-PLK4 dox -
Panc1-PLK4 dox 3d
Comp-FL1-A :: Hoechst 33342-A
E)
Panc1
F)
Mia Paca-2
dox-
dox+
BSO
ML385
Colony formation relative to DMSO (log2)
BSO (μM)
BSO (mM)
ML385 (μM)

Fig S2

**Supplementary Figure 2: Supplemental to Figure 2.**

**A)** Intracellular ROS levels in centrosome-amplified cells treated with CB-893, ML385, or BSO. Centrosome amplification was induced for 5 days, followed by inhibitor treatment for 48 hours prior to quantification using H2DCFDA and flow cytometry.

**B)** Quantification of ROS levels from panel A. Mean fluorescence intensity fold changes are shown with statistical significance determined by two-way ANOVA test.

**C)** Doxycycline treatment alone does not increase ARE-driven reporter activity in Panc1 cells. $BFP^+$ cells were gated, and GFP intensity was measured.

**D)** Representative BFP expression patterns in Panc1-PLK4 and control Panc1 cells used in the ARE reporter experiments.

**E)** BSO treatment reduces colony formation of centrosome-amplified Panc1 cells compared to non-centrosome amplified controls.

**F)** ML385 decreases colony formation in centrosome-amplified Mia Paca-2 cells, whereas BSO treatment does not exhibit a selective effect.

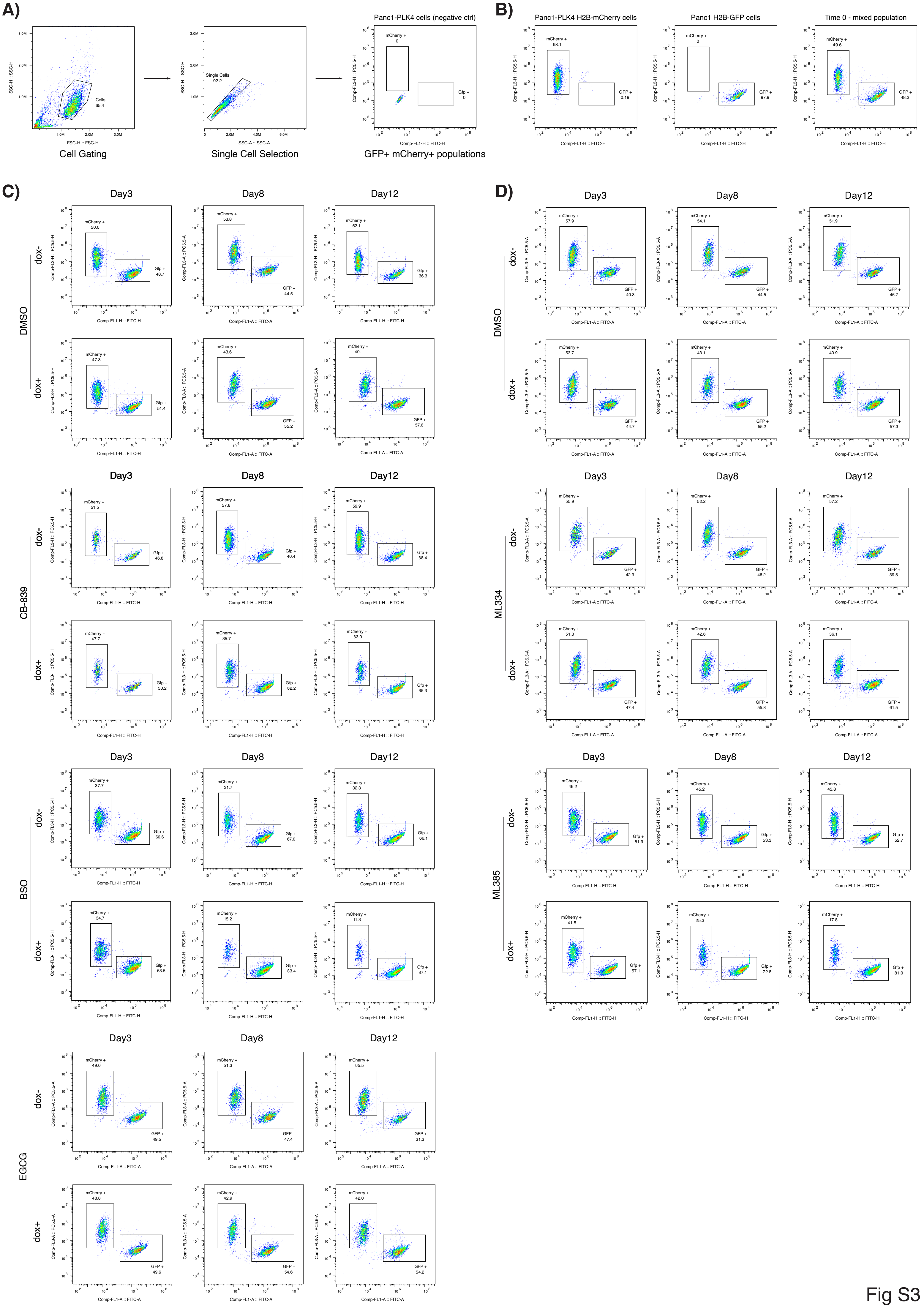
A)
Cell Gating
Single Cell Selection
Panc1-PLK4 cells (negative ctrl)
GFP+ mCherry+ populations
B)
Panc1-PLK4 H2B-mCherry cells
Panc1 H2B-GFP cells
Time 0 - mixed population
C)
D)
DMSO
CB-839
BSO
EGCG
ML334
ML385
dox-
dox+
Day3
Day8
Day12

Fig S3

**Supplementary Figure 3: Supplemental to Figure 2.** Representative flow cytometry results of competition experiments in Figure 2I and 2J.

**A)** Cell gating strategy used in competition experiments.

**B)** Representative GFP and mCherry fluorescence profiles of Panc1-PLK4-H2B-mCherry, Panc1-H2B-GFP, and mixed populations at the initial time point.

**C)** Representative flow cytometry plots from competition experiments shown in Figure 2I.

**D)** Representative flow cytometry plots from competition experiments shown in Figure 2J.

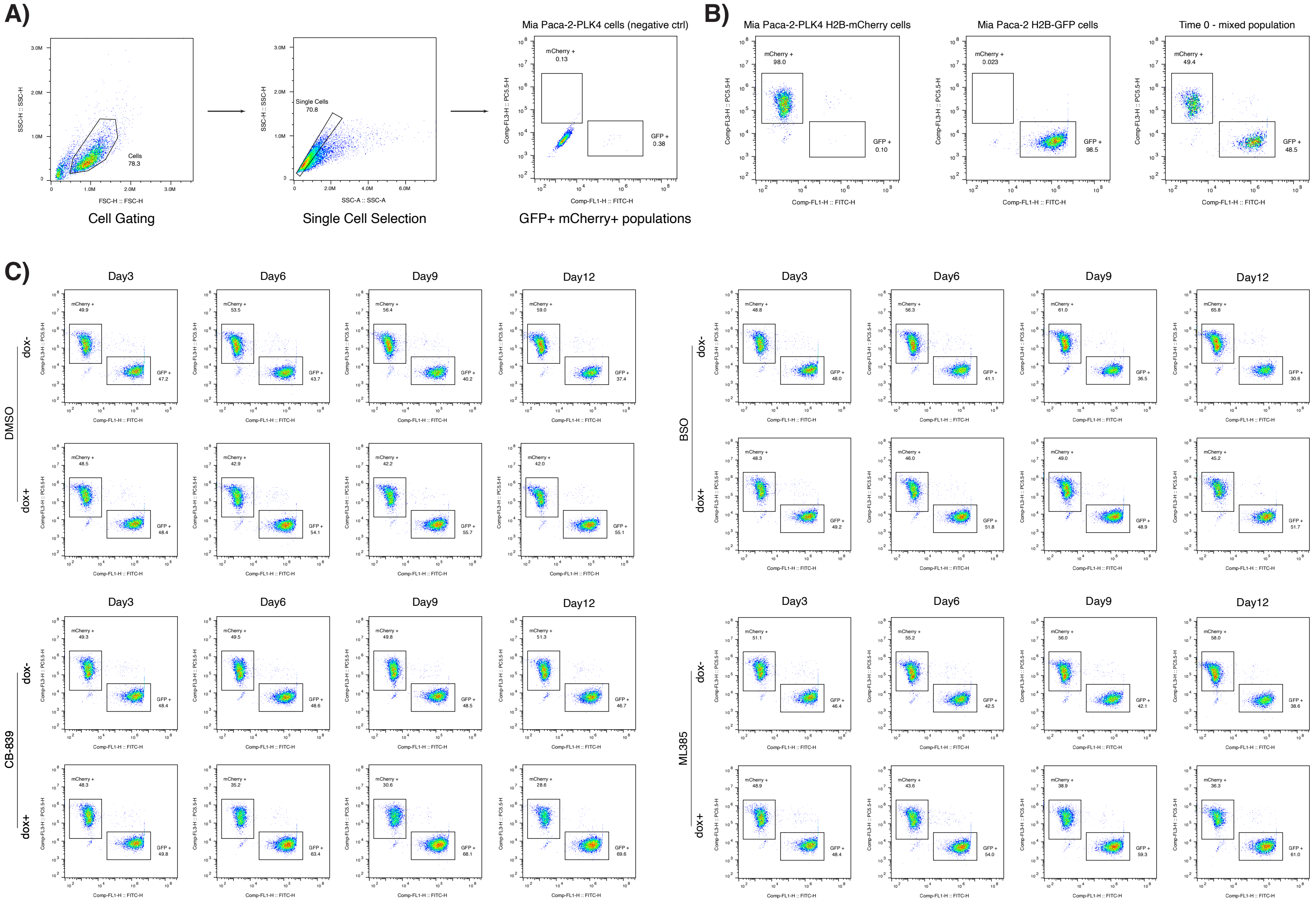


**Supplementary Figure 4: Supplemental to Figure 2.** Representative flow cytometry results of competition experiments in Figure 2K.

**A)** Cell gating strategy used in competition experiments.

**B)** Representative GFP and mCherry fluorescence profiles of Mia Paca-2-PLK4-H2B-mCherry, Mia Paca-2-H2B-GFP, and mixed populations at the initial time point.

**C)** Representative flow cytometry plots from competition experiments shown in Figure 2J.

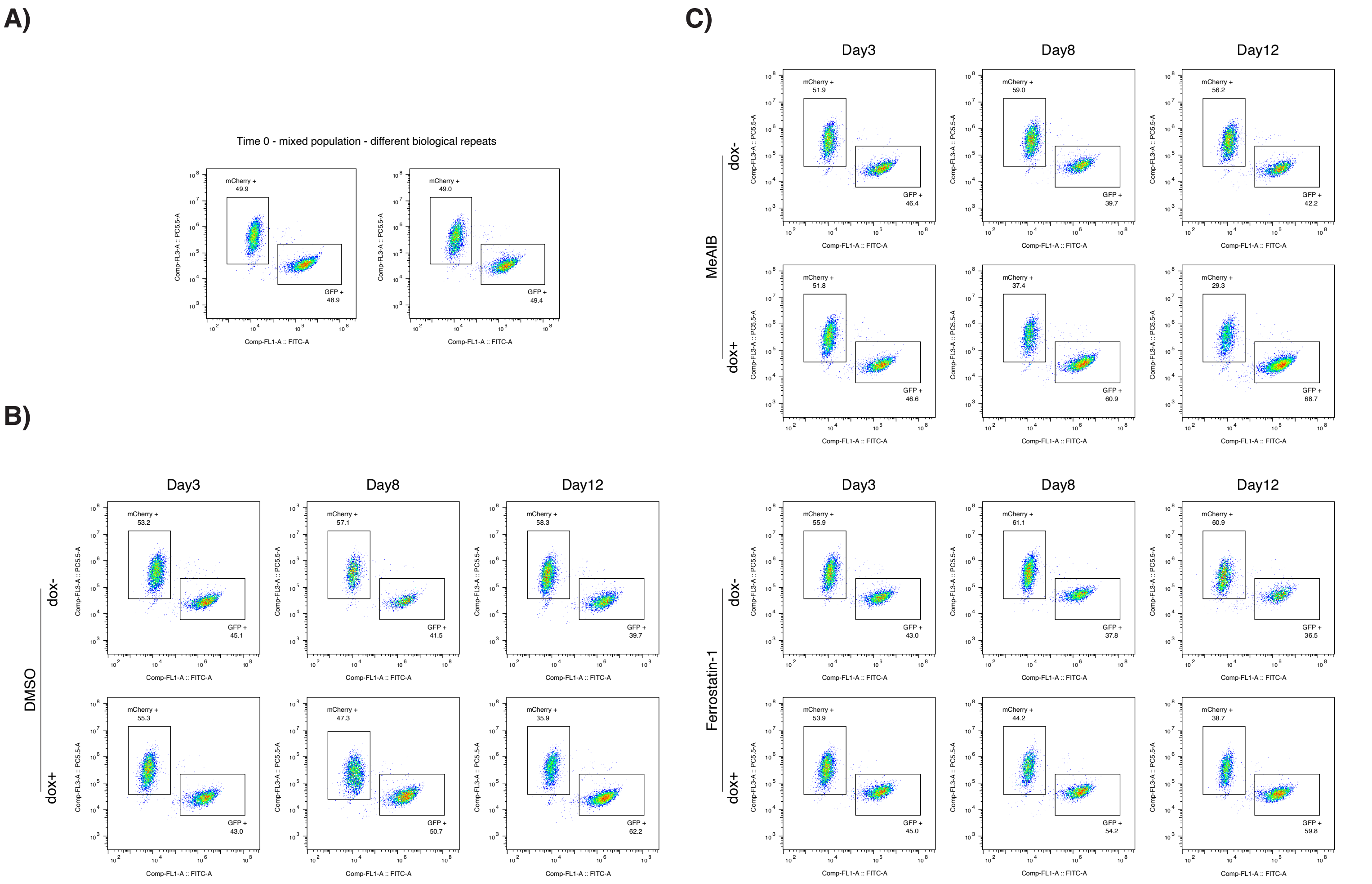


**Supplementary Figure 5: Supplemental to Figure 2.** Representative flow cytometry plots from competition experiments shown in Figure 2L.
**A)** Representative mixed populations at the initial time point.
**B)** Representative flow cytometry plots of DMSO-treated cells.
**C)** Representative flow cytometry plots of  MeAIB and Ferrostatin-1 treated cells.

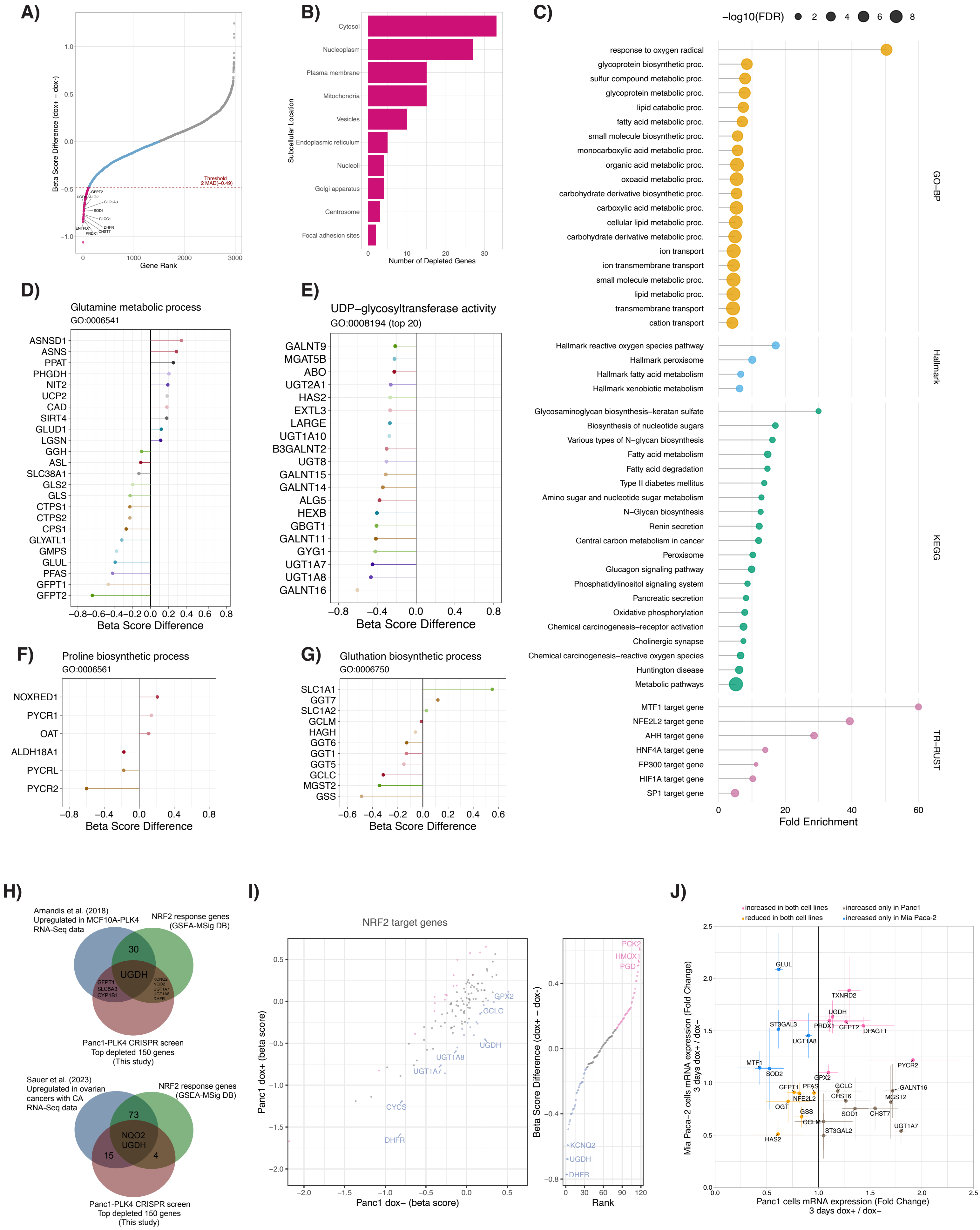

A)
Beta Score Difference (dox+ − dox-)
Gene Rank
Threshold
2 MAD(−0.49)
GFPT2
UGDH
ALG2
SLC5A3
SOD1
CLCC1
DHFR
ENTPD7
PRDX1
CHST7
B)
Subcellular Location
Cytosol
Nucleoplasm
Plasma membrane
Mitochondria
Vesicles
Endoplasmic reticulum
Nucleoli
Golgi apparatus
Centrosome
Focal adhesion sites
Number of Depleted Genes
C)
−log10(FDR)
response to oxygen radical
glycoprotein biosynthetic proc.
sulfur compound metabolic proc.
glycoprotein metabolic proc.
lipid catabolic proc.
fatty acid metabolic proc.
small molecule biosynthetic proc.
monocarboxylic acid metabolic proc.
organic acid metabolic proc.
oxoacid metabolic proc.
carbohydrate derivative biosynthetic proc.
carboxylic acid metabolic proc.
cellular lipid metabolic proc.
carbohydrate derivative metabolic proc.
ion transport
ion transmembrane transport
small molecule metabolic proc.
lipid metabolic proc.
transmembrane transport
cation transport
GO−BP
Hallmark reactive oxygen species pathway
Hallmark peroxisome
Hallmark fatty acid metabolism
Hallmark xenobiotic metabolism
Hallmark
Glycosaminoglycan biosynthesis−keratan sulfate
Biosynthesis of nucleotide sugars
Various types of N−glycan biosynthesis
Fatty acid metabolism
Fatty acid degradation
Type II diabetes mellitus
Amino sugar and nucleotide sugar metabolism
N−Glycan biosynthesis
Renin secretion
Central carbon metabolism in cancer
Peroxisome
Glucagon signaling pathway
Phosphatidylinositol signaling system
Pancreatic secretion
Oxidative phosphorylation
Chemical carcinogenesis−receptor activation
Cholinergic synapse
Chemical carcinogenesis−reactive oxygen species
Huntington disease
Metabolic pathways
KEGG
MTF1 target gene
NFE2L2 target gene
AHR target gene
HNF4A target gene
EP300 target gene
HIF1A target gene
SP1 target gene
TR−RUST
Fold Enrichment
D)
Glutamine metabolic process
GO:0006541
ASNSD1
ASNS
PPAT
PHGDH
NIT2
UCP2
CAD
SIRT4
GLUD1
LGSN
GGH
ASL
SLC38A1
GLS2
GLS
CTPS1
CTPS2
CPS1
GLYATL1
GMPS
GLUL
PFAS
GFPT1
GFPT2
Beta Score Difference
E)
UDP−glycosyltransferase activity
GO:0008194 (top 20)
GALNT9
MGAT5B
ABO
UGT2A1
HAS2
EXTL3
LARGE
UGT1A10
B3GALNT2
UGT8
GALNT15
GALNT14
ALG5
HEXB
GBGT1
GALNT11
GYG1
UGT1A7
UGT1A8
GALNT16
Beta Score Difference
F)
Proline biosynthetic process
GO:0006561
NOXRED1
PYCR1
OAT
ALDH18A1
PYCRL
PYCR2
Beta Score Difference
G)
Gluthation biosynthetic process
GO:0006750
SLC1A1
GGT7
SLC1A2
GCLM
HAGH
GGT6
GGT1
GGT5
GCLC
MGST2
GSS
Beta Score Difference
H)
Arnandis et al. (2018)
Upregulated in MCF10A-PLK4
RNA-Seq data
NRF2 response genes
(GSEA-MSig DB)
30
UGDH
GFPT1
SLC5A3
CYP1B1
KCNQ2
NQO2
UGT1A7
UGT1A8
DHFR
Panc1-PLK4 CRISPR screen
Top depleted 150 genes
(This study)
Sauer et al. (2023)
Upregulated in ovarian
cancers with CA
RNA-Seq data
NRF2 response genes
(GSEA-MSig DB)
73
NQO2
UGDH
15
4
Panc1-PLK4 CRISPR screen
Top depleted 150 genes
(This study)
I)
NRF2 target genes
GPX2
GCLC
UGDH
UGT1A8
UGT1A7
CYCS
DHFR
Panc1 dox+ (beta score)
Panc1 dox− (beta score)
PCK2
HMOX1
PGD
KCNQ2
UGDH
DHFR
Beta Score Difference (dox+ − dox-)
Rank
J)
increased in both cell lines
reduced in both cell lines
increased only in Panc1
increased only in Mia Paca-2
GLUL
TXNRD2
UGDH
ST3GAL3
PRDX1
GFPT2
DPAGT1
UGT1A8
MTF1
SOD2
PYCR2
GPX2
GFPT1
PFAS
GCLC
GALNT16
NFE2L2
CHST6
MGST2
OGT
GSS
SOD1
CHST7
GCLM
ST3GAL2
UGT1A7
HAS2
Mia Paca−2 cells mRNA expression (Fold Change)
3 days dox+ / dox−
Panc1 cells mRNA expression (Fold Change)
3 days dox+ / dox−


Fig S6

**Supplementary Figure 6: Supplemental to Figure 3.**
**A)** Ranked plot of CRISPR screen hits based on differential β-scores. A 2 MAD threshold (−0.49) was applied, and genes beyond this cutoff (109 genes) were considered as top hits and used for enrichment analysis in panel C.
**B)** Subcellular localization of the top 109 differentially depleted genes.
**C)** Functional enrichment analysis of the top depleted genes.
GO-BP: Gene Ontology – Biological Process
Hallmark: Molecular Signatures Database Hallmark gene sets
KEGG: Kyoto Encyclopedia of Genes and Genomes
TR-RUST: Transcriptional Regulatory Relationships Unraveled by Sentence-based Text mining.
**D-G)** Pathway-specific gene depletion patterns in cells with centrosome amplification:
(D) Glutamine metabolic process (GO:0006541),
(E) UDP-glycosyltransferase activity (GO:0008194),
(F) Proline biosynthetic process (GO:0006561),
(G) Glutathione biosynthetic process (GO:0006750).
**H)** Venn diagrams showing the overlap of top CRISPR screen hits with prior gene expression studies and known NRF2 target genes.
**I)** CRISPR screen results filtered for NRF2 target genes. Left panel: Scatterplot of β-scores for dox+ and dox− comparisons with the initial sample. Pink dots indicate genes with increased β-scores after centrosome amplification; blue dots indicate genes with decreased β-scores. Right panel: Ranked plot of differential β-scores (dox+ - dox−).
**J)** mRNA expression levels of selected genes in centrosome amplified Panc1 and Mia Paca-2 cells.

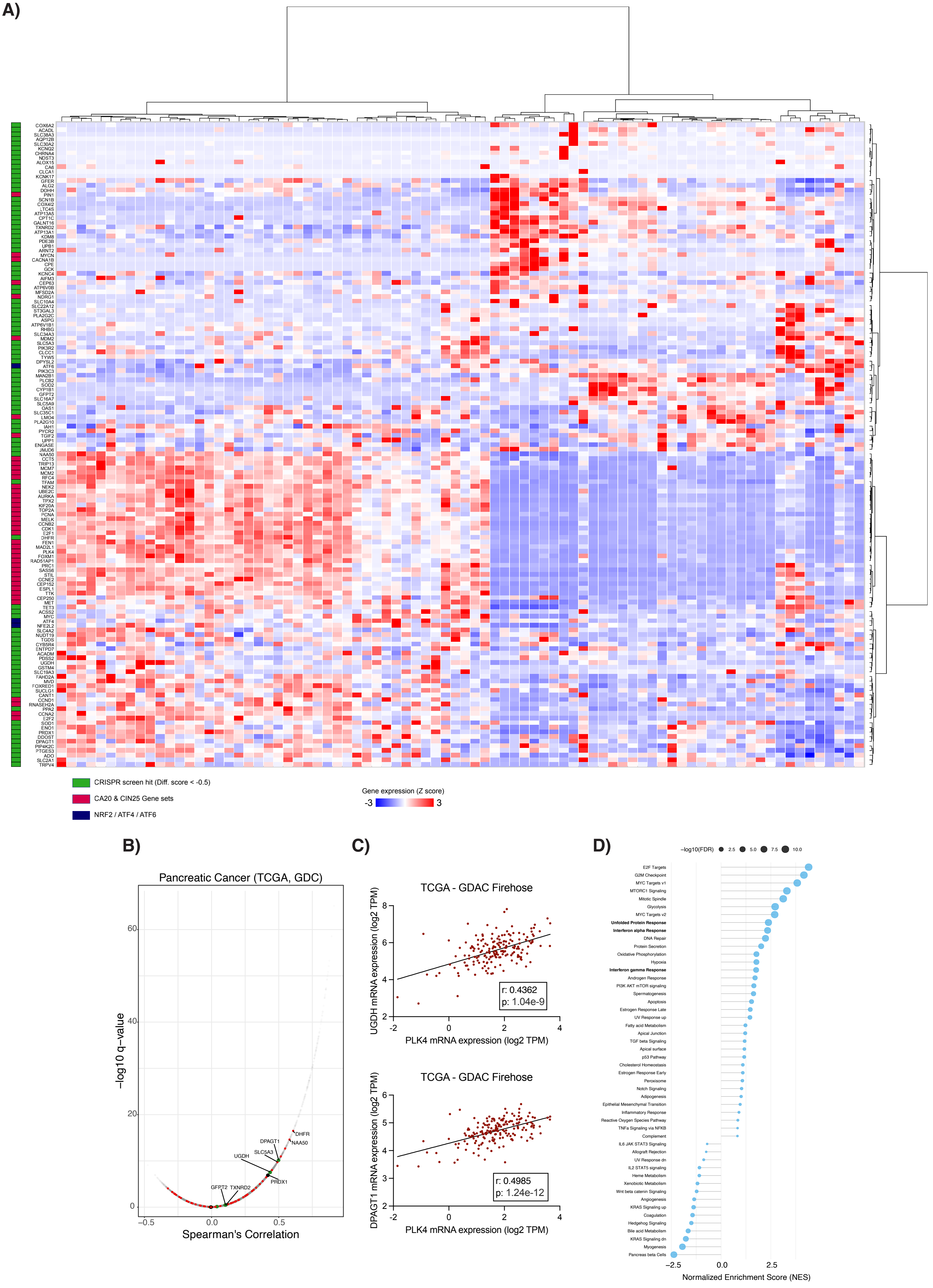
A)
COX6A2
ACADL
SLC38A3
AQP12B
SLC30A2
KCNQ2
CHRNA4
NDST3
ALOX15
CA6
CLCA1
KCNK17
GFER
ALG2
DOHH
PIN1
SCN1B
COX4I2
LTC4S
ATP13A5
CPT1C
GALNT16
TXNRD2
ATP13A1
KDM8
PDE3B
UPB1
ARNT2
MYCN
CACNA1B
CPE
GCK
KCNC4
AIFM3
CEP63
ATP6V0B
MFSD2A
NDRG1
SLC10A4
SLC22A12
ST3GAL3
PLA2G2C
ASPG
ATP6V1B1
RHBG
SLC34A3
MDM2
SLC5A3
PIK3R2
CLCC1
TYW5
DPYSL2
ATF6
PIK3C3
MAN2B1
PLCB2
SOD2
CYP1B1
GFPT2
SLC16A7
SLC5A9
OAS1
SLC35C1
LMO4
PLA2G10
IAH1
PYCR2
TGIF2
UPP1
ENGASE
JMJD6
NAA50
CCT5
TRIP13
MCM7
MCM2
RFC4
TFAM
NEK2
UBE2C
AURKA
TPX2
KIF20A
TOP2A
PCNA
MELK
CCNB2
CDK1
E2F1
DHFR
FEN1
MAD2L1
PLK4
FOXM1
RAD51AP1
PRC1
SASS6
STIL
CCNE2
CEP152
ESPL1
TTK
CEP250
MET
TET3
ACSS2
MYC
ATF4
NFE2L2
SLC4A2
NUDT19
TGDS
CYB5R4
ENTPD7
ACADM
PDSS2
UGDH
GSTM4
SLC19A3
FAHD2A
MVD
FOXRED1
SUCLG1
CANT1
CCND1
RNASEH2A
PPA2
CCNA2
E2F2
SOD1
ENO1
PRDX1
DDOST
DPAGT1
PIP4K2C
PTGES3
ADO
SLC2A1
TRPV4
CRISPR screen hit (Diff. score < -0.5)
CA20 & CIN25 Gene sets
NRF2 / ATF4 / ATF6
Gene expression (Z score)
-3
3
B)
Pancreatic Cancer (TCGA, GDC)
−log10 q−value
Spearman's Correlation
DHFR
NAA50
DPAGT1
SLC5A3
UGDH
PRDX1
GFPT2
TXNRD2
C)
TCGA - GDAC Firehose
UGDH mRNA expression (log2 TPM)
PLK4 mRNA expression (log2 TPM)
r: 0.4362
p: 1.04e-9
TCGA - GDAC Firehose
DPAGT1 mRNA expression (log2 TPM)
PLK4 mRNA expression (log2 TPM)
r: 0.4985
p: 1.24e-12
D)
−log10(FDR)
2.5
5.0
7.5
10.0
E2F Targets
G2M Checkpoint
MYC Targets v1
MTORC1 Signaling
Mitotic Spindle
Glycolysis
MYC Targets v2
Unfolded Protein Response
Interferon alpha Response
DNA Repair
Protein Secretion
Oxidative Phosphorylation
Hypoxia
Interferon gamma Response
Androgen Response
PI3K AKT mTOR signaling
Spermatogenesis
Apoptosis
Estrogen Response Late
UV Response up
Fatty acid Metabolism
Apical Junction
TGF beta Signaling
Apical surface
p53 Pathway
Cholesterol Homeostasis
Estrogen Response Early
Peroxisome
Notch Signaling
Adipogenesis
Epithelial Mesenchymal Transition
Inflammatory Response
Reactive Oxygen Species Pathway
TNFa Signaling via NFKB
Complement
IL6 JAK STAT3 Signaling
Allograft Rejection
UV Response dn
IL2 STAT5 signaling
Heme Metabolism
Xenobiotic Metabolism
Wnt beta catenin Signaling
Angiogenesis
KRAS Signaling up
Coagulation
Hedgehog Signaling
Bile acid Metabolism
KRAS Signaling dn
Myogenesis
Pancreas beta Cells
Normalized Enrichment Score (NES)

Fig S7

**Supplementary Figure 7: Supplemental to Figure 3.**
**A)** Heatmap of TCGA pancreatic adenocarcinoma patient data (n = 82) showing expression levels of the top depleted CRISPR screen hits, CIN25 and CA20 gene sets, and NRF2, ATF4, and ATF6. Z-scores of gene expression were used, and unsupervised clustering was applied to the dataset.
**B)** Correlation analysis between PLK4 expression and top CRISPR screen hits. Genes from the metabolic enzyme library are marked in red; genes in ROS detoxification pathways in black; and genes in N-glycan synthesis/amino sugar metabolism in green.
**C)** Scatter plots showing the correlations of UGDH and DPAGT1 expression with PLK4 in TCGA pancreatic adenocarcinoma dataset.
**D)** GENI analysis results showing enrichment patterns associated with PLK4 expression in the TCGA pancreatic adenocarcinoma dataset.
