## Supplemental Figures and Legends - 2 for "Stress adaptation pathways and HA–CD44 signaling maintain the survival of pancreatic cancer cells with centrosome amplification"

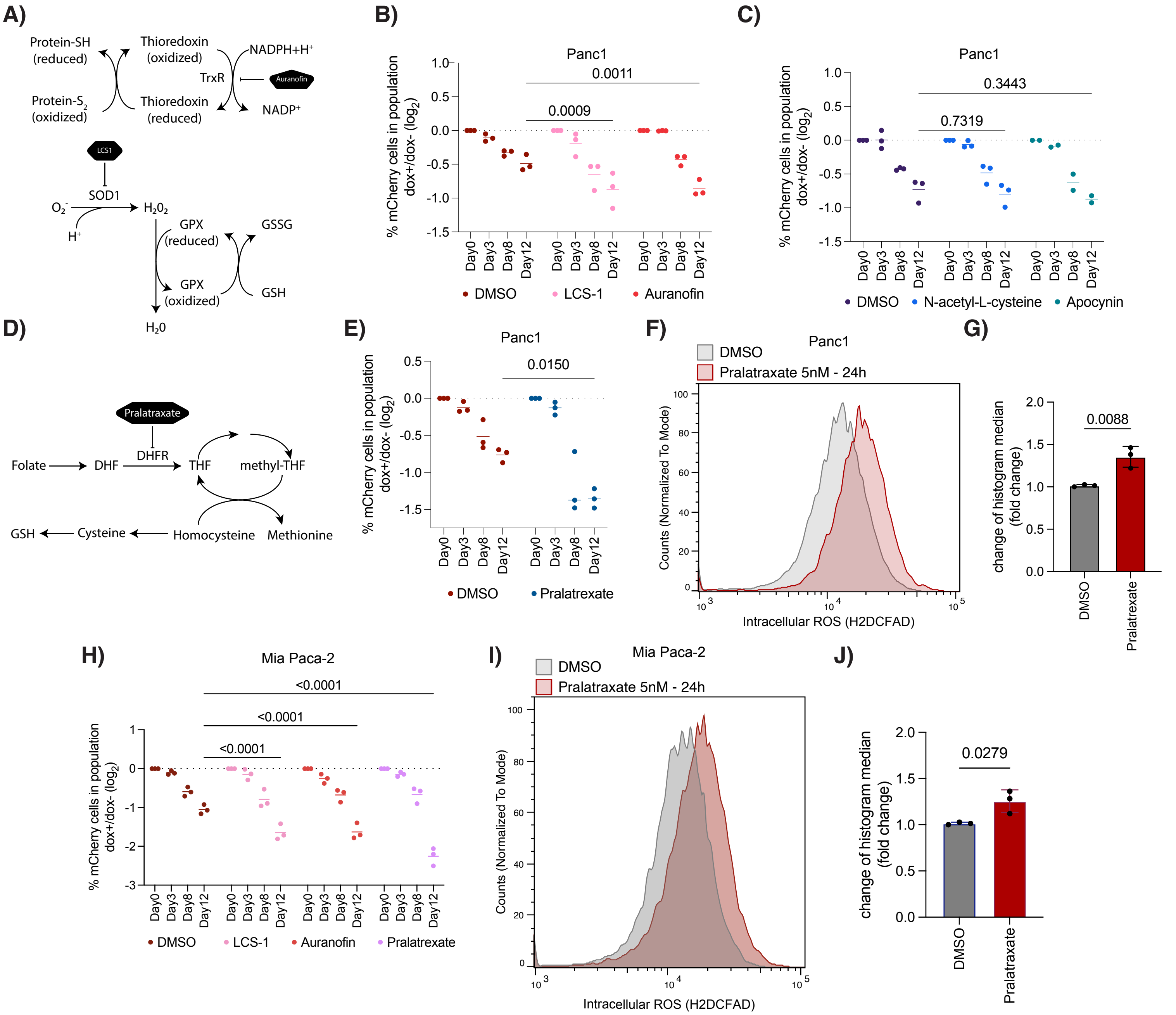


**Supplementary Figure 8: ROS elimination pathways are selective dependencies in PDAC cells with centrosome amplification.**

**A)** Schematic illustration showing the roles of SOD1 and thioredoxin reactions in oxidation–reduction processes.

**B)** Treatment with LCS-1 or auranofin increases depletion of centrosome-amplified Panc1 cells in competition assays.

**C)** Treatment with N-acetyl-cysteine or apocynin does not significantly affect centrosome-amplified Panc1 cells in competition assays.

**D)** Schematic illustration showing the reaction catalyzed by DHFR.

**E)** DHFR inhibition by Pralatrexate enhances depletion of centrosome-amplified Panc1 cells in competition assays.

**F)** Pralatrexate treatment increases intracellular ROS levels in Panc1 cells with CA.

**G)** Quantification of ROS level changes in centrosome-amplified Panc1 cells. Each dot represents an independent replicate.

**H)** Treatment with LCS-1, Auranofin and Pralatraxate increases depletion of centrosome-amplified Mia Paca-2 cells in competition assays.

**I)** Pralatrexate treatment increases intracellular ROS levels in Mia Paca-2 cells with CA.

**J)** Quantification of ROS level changes in centrosome-amplified Mia Paca-2 cells. Each dot represents an independent replicate.

Statistical significance was determined by two-way ANOVA for panels B, C, E, and H, and by two-way t-test for panel G, J. p values were shown on plots.

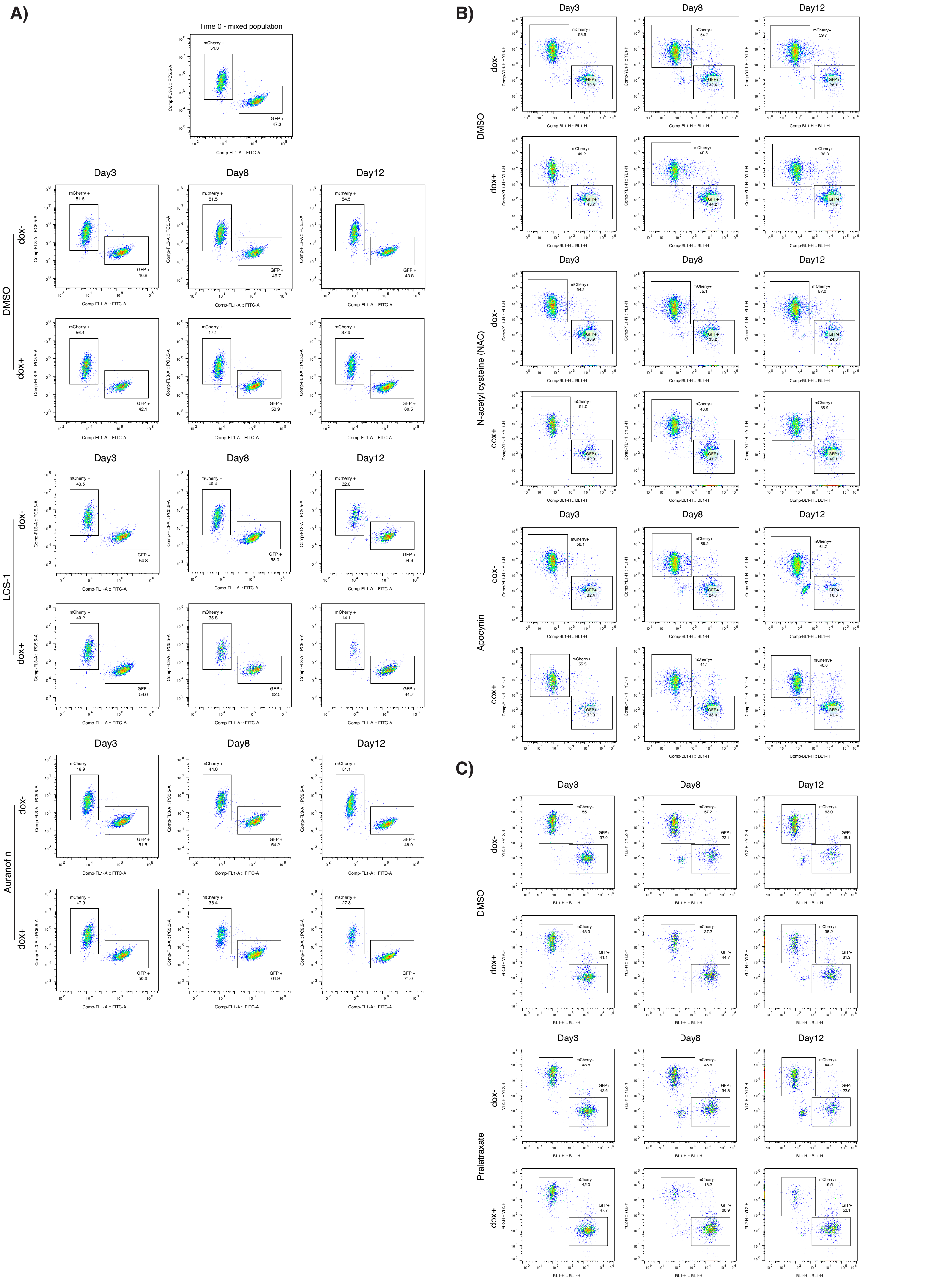
A)
Time 0 - mixed population
mCherry + 51.3
GFP + 47.3
Comp-FL3-A :: PC5.5-A
Comp-FL1-A :: FITC-A
Day3
Day8
Day12
DMSO
dox-
mCherry + 51.5
GFP + 46.8
mCherry + 51.5
GFP + 46.7
mCherry + 54.5
GFP + 43.8
dox+
mCherry + 56.4
GFP + 42.1
mCherry + 47.1
GFP + 50.9
mCherry + 37.9
GFP + 60.5
LCS-1
dox-
mCherry + 43.5
GFP + 54.8
mCherry + 40.4
GFP + 58.0
mCherry + 32.0
GFP + 64.8
dox+
mCherry + 40.2
GFP + 58.6
mCherry + 35.8
GFP + 62.5
mCherry + 14.1
GFP + 84.7
Auranofin
dox-
mCherry + 46.9
GFP + 51.5
mCherry + 44.0
GFP + 54.2
mCherry + 51.1
GFP + 46.9
dox+
mCherry + 47.9
GFP + 50.6
mCherry + 33.4
GFP + 64.9
mCherry + 27.3
GFP + 71.0
B)
Day3
Day8
Day12
Comp-YL1-H :: YL1-H
Comp-BL1-H :: BL1-H
DMSO
dox-
mCherry+ 53.6
GFP+ 39.8
mCherry+ 54.7
GFP+ 32.4
mCherry+ 59.7
GFP+ 26.1
dox+
mCherry+ 49.2
GFP+ 43.7
mCherry+ 40.8
GFP+ 44.2
mCherry+ 38.3
GFP+ 41.9
N-acetyl cysteine (NAC)
dox-
mCherry+ 54.2
GFP+ 38.9
mCherry+ 55.1
GFP+ 33.2
mCherry+ 57.0
GFP+ 24.3
dox+
mCherry+ 51.0
GFP+ 42.0
mCherry+ 43.0
GFP+ 41.7
mCherry+ 35.9
GFP+ 45.1
Apocynin
dox-
mCherry+ 58.1
GFP+ 32.4
mCherry+ 58.2
GFP+ 24.7
mCherry+ 61.2
GFP+ 10.3
dox+
mCherry+ 55.3
GFP+ 32.0
mCherry+ 41.1
GFP+ 38.0
mCherry+ 40.0
GFP+ 41.4
C)
Day3
Day8
Day12
YL2-H :: YL2-H
BL1-H :: BL1-H
DMSO
dox-
mCherry+ 55.1
GFP+ 37.0
mCherry+ 57.2
GFP+ 23.1
mCherry+ 63.0
GFP+ 18.1
dox+
mCherry+ 48.9
GFP+ 41.1
mCherry+ 37.2
GFP+ 44.7
mCherry+ 35.2
GFP+ 31.3
Pralatrexate
dox-
mCherry+ 48.8
GFP+ 42.6
mCherry+ 45.6
GFP+ 34.8
mCherry+ 44.2
GFP+ 22.6
dox+
mCherry+ 42.0
GFP+ 47.7
mCherry+ 18.2
GFP+ 60.9
mCherry+ 16.5
GFP+ 53.1

Fig S9

**Supplementary Figure 9: Supplemental to Supplementary Figure 8.** Representative flow cytometry plots from competition experiments shown in Supplementary Figure 8B, 8C and 8E.
**A)** Representative flow cytometry plots of DMSO, LCS-1 and auranofin treated cells. (Fig. S8B)
**B)** Representative flow cytometry plots of DMSO, NAC and apocynin treated cells. (Fig. S8C)
**C)** Representative flow cytometry plots of DMSO or Pralatraxate treated cells. (Fig. S8E)

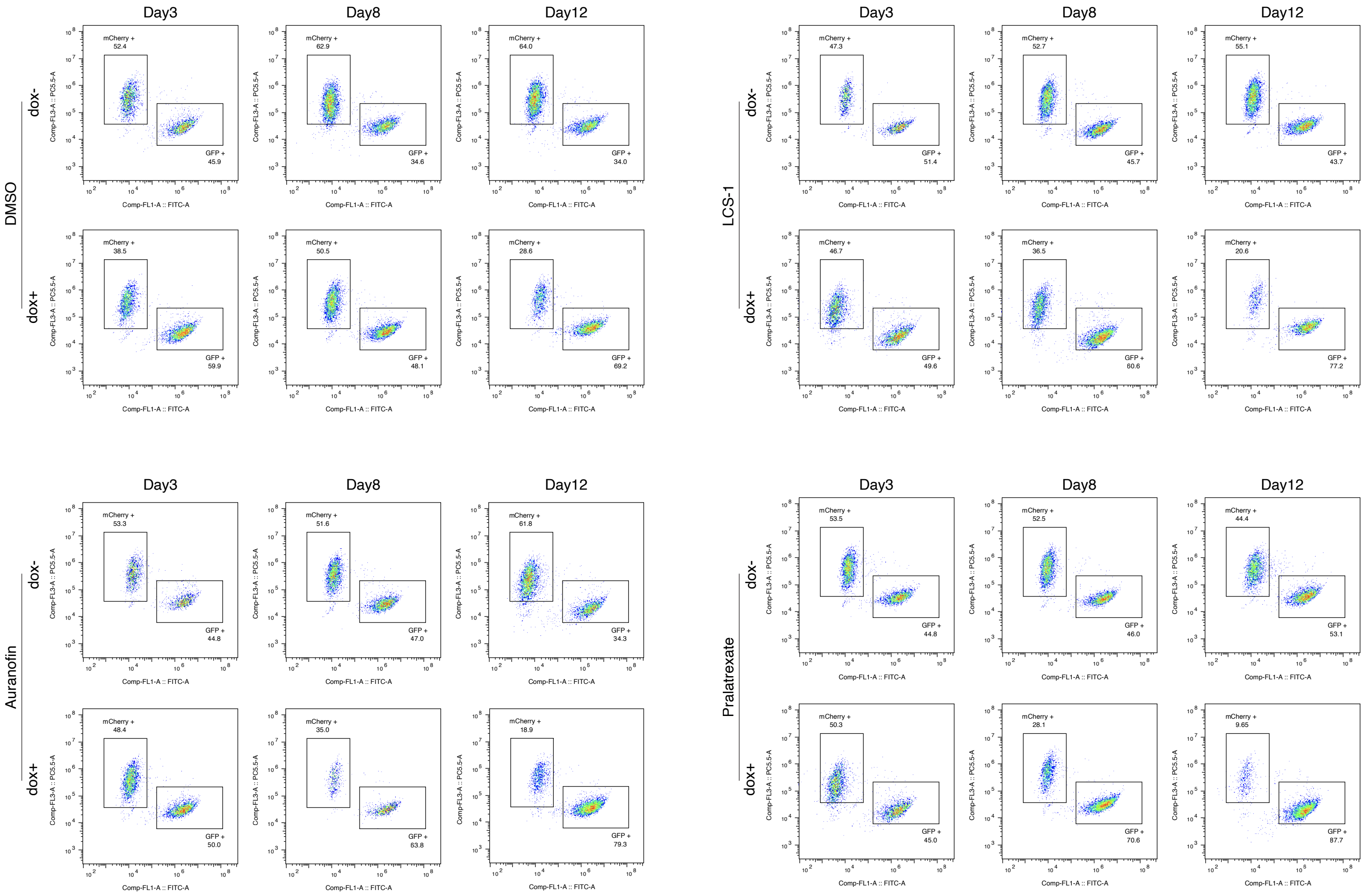


**Supplementary Figure 10: Supplemental to Supplementary Figure 8.** Representative flow cytometry plots from competition experiments shown in Supplementary Figure 8H.

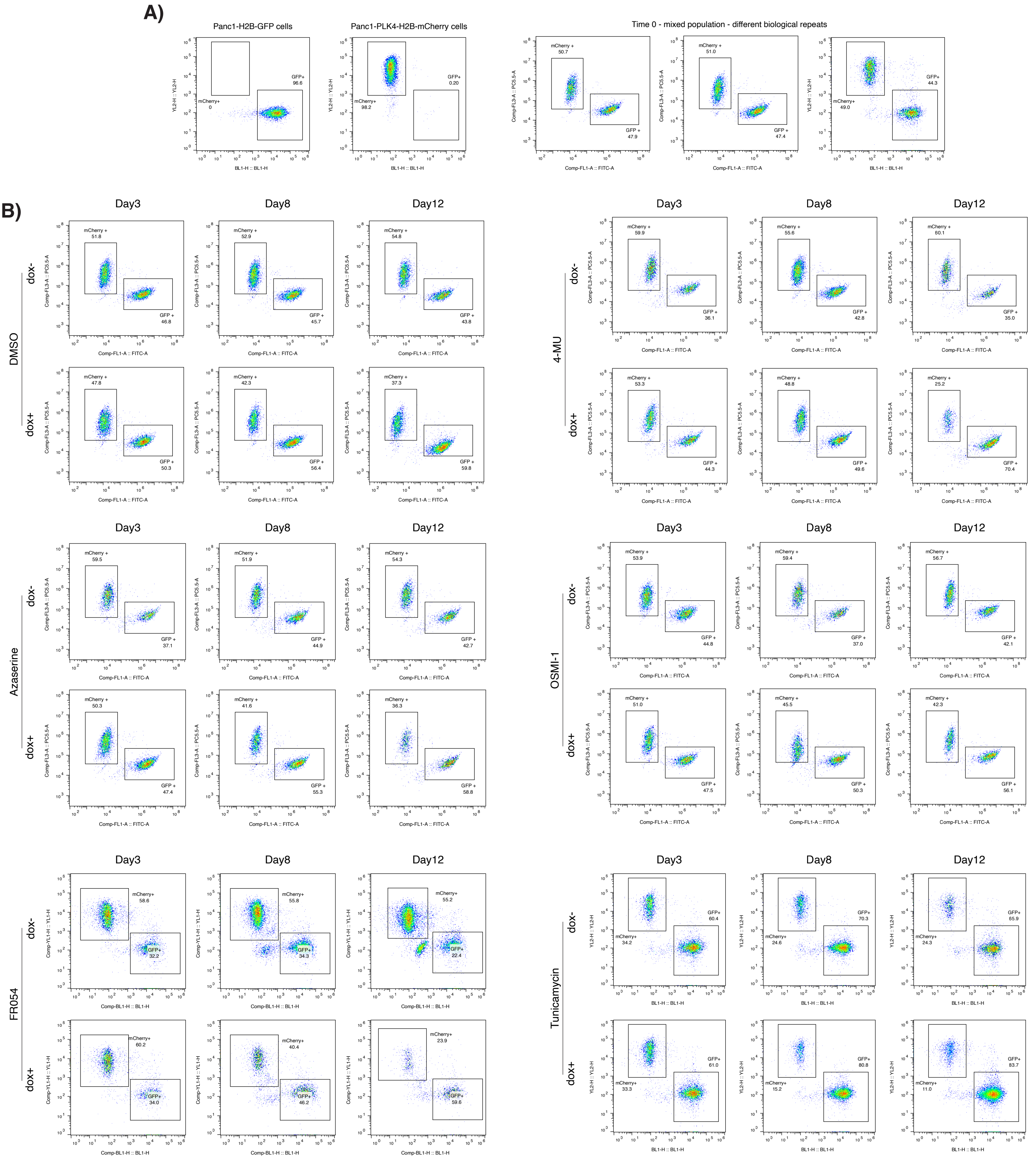


**Supplementary Figure 11: Supplemental to Figure 4.** Representative flow cytometry plots from competition experiments shown in Figure 4B.

**A)** Representative GFP and mCherry fluorescence profiles of Panc1-PLK4-H2B-mCherry, Panc1-H2B-GFP, and mixed populations at the initial time point.

**B)** Representative flow cytometry plots from competition experiments shown in Figure 4B.

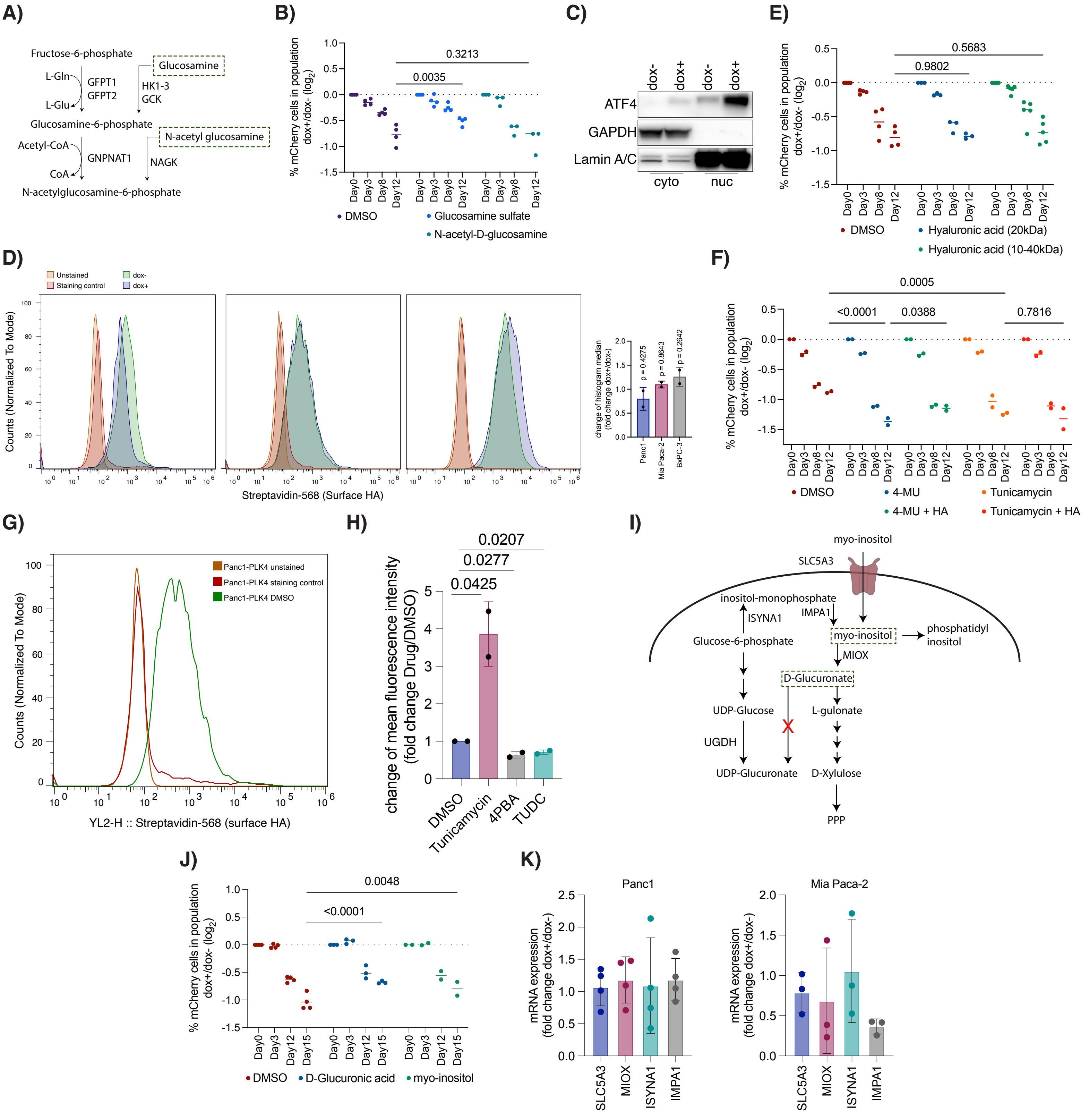

Fig S12

**Supplementary Figure 12: Supplemental to Figure 4.**
**A)** Schematic illustration of the salvage pathways of the hexosamine biosynthesis pathway. Metabolites used in the study are highlighted in green.
**B)** Glucosamine sulfate treatment alleviates the depletion of centrosome-amplified cells in competition experiments.
**C)** Centrosome amplification increases nuclear localization of ATF4. GAPDH and Lamin A/C served as fractionation and loading controls.
**D)** Doxycycline treatment of cells lacking the doxPLK4 construct does not affect surface HA levels.
**E)** Hyaluronic acid supplementation does not alter the reduction of centrosome-amplified cells in competition experiments.
**F)** Hyaluronic acid supplementation rescues the depletion of 4-MU–treated centrosome-amplified cells but does not affect depletion with tunicamycin treatment.
**G)** Control stainings for Fig. 4J. The Panc1-PLK4 DMSO group shown here is identical to that in Fig. 4J.
**H)** Tunicamycin treatment increases surface HA levels in centrosome-amplified cells, whereas TUDC and 4-PBA cause a decrease.
**I)** Schematic of the SLC5A3–myo-inositol pathway.
**J)** D-glucuronic acid and inositol supplementation alleviate depletion of centrosome-amplified cells in competition experiments.
**K)** mRNA expression levels of SLC5A3, MIOX, ISYNA1, and IMPA1 after three days of centrosome amplification.

Statistical significance was determined by two-way ANOVA for panels B, E, F, and J, and by Student's t-test for panel D and H. p values were shown on plots.

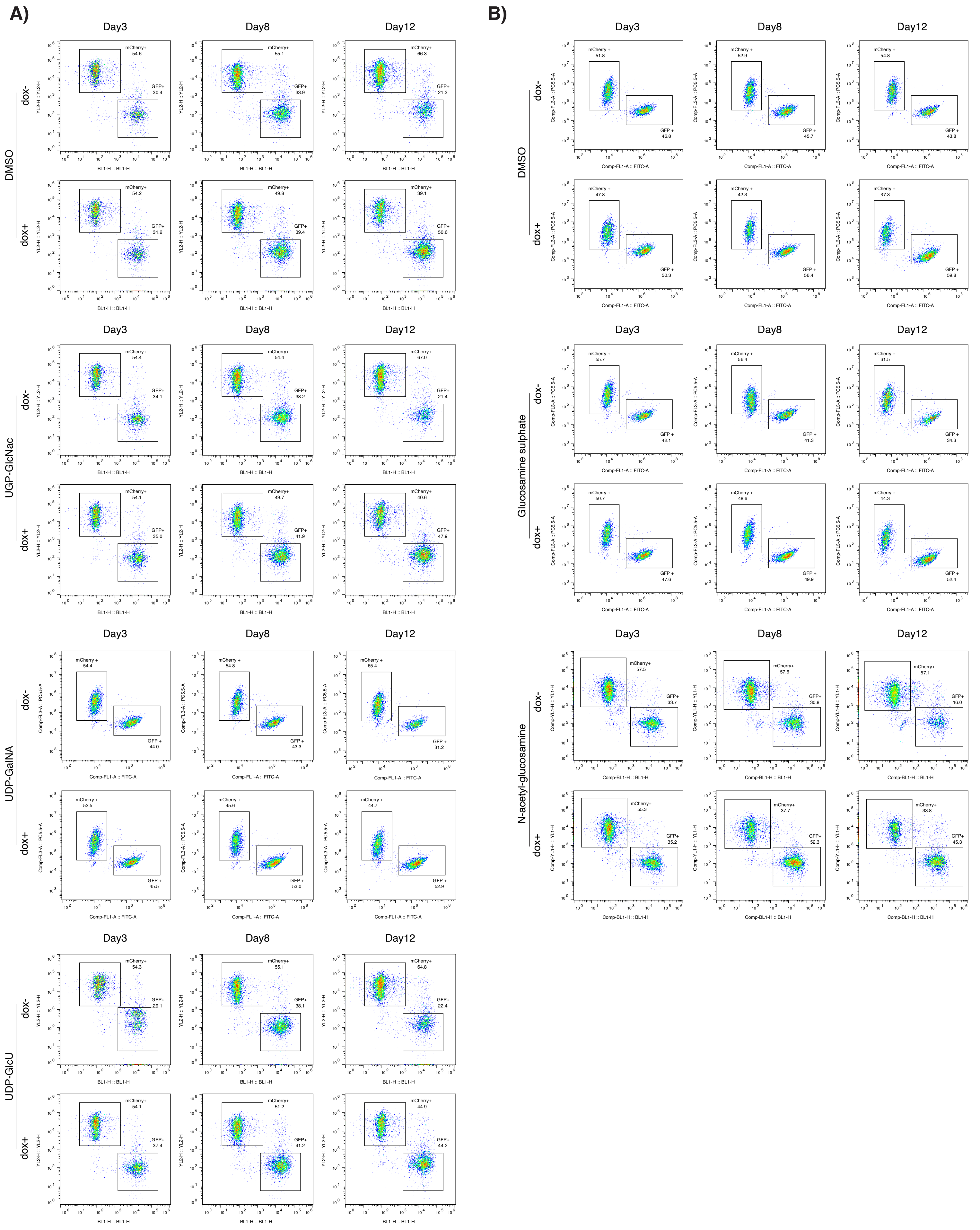


**Supplementary Figure 13: Supplemental to Figure 4.** Representative flow cytometry results of competition experiments in Figure 4C and S12B.
**A)** Representative flow cytometry plots from competition experiments shown in Figure 4C.
**B)** Representative flow cytometry plots from competition experiments shown in Figure S11B.

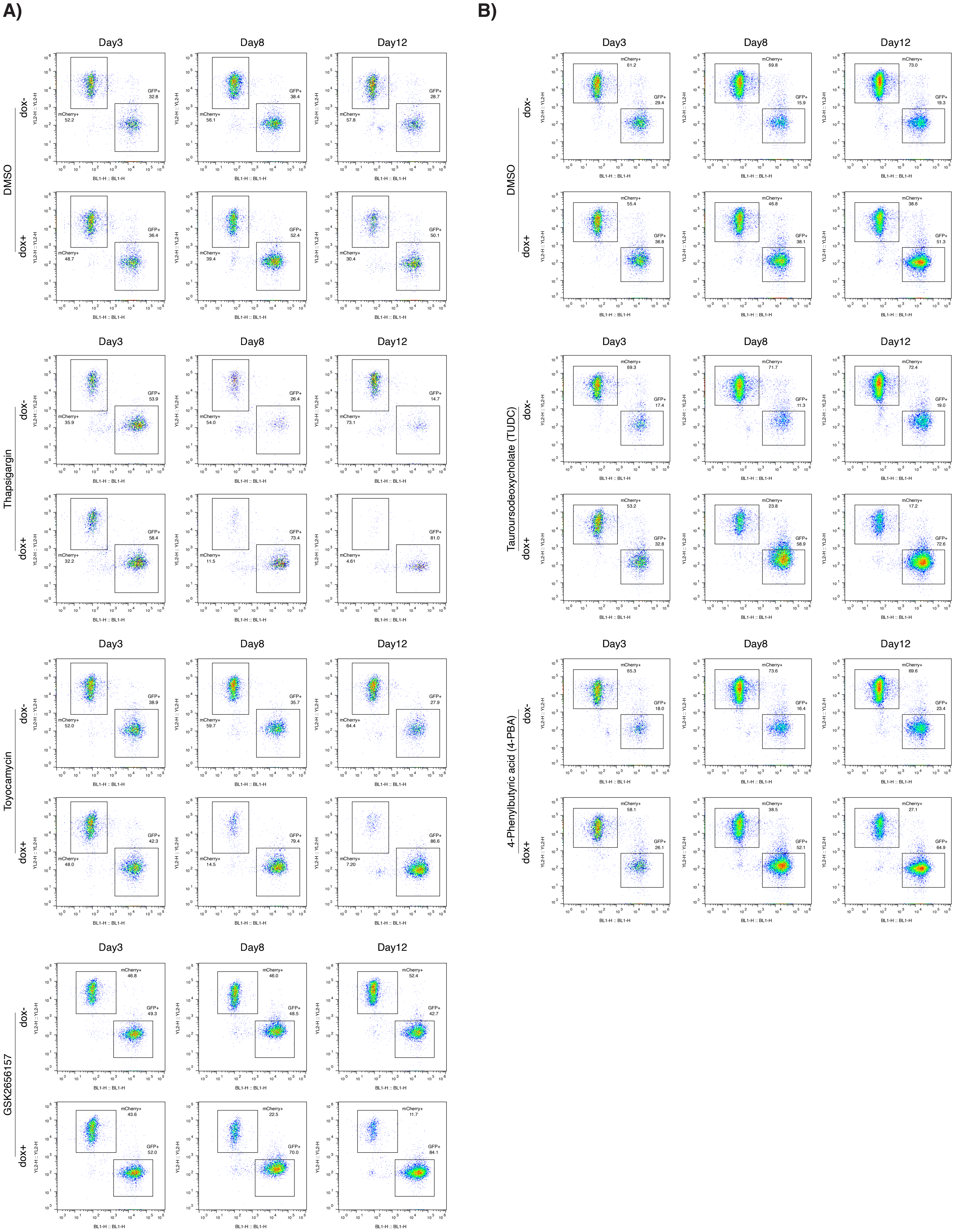


**Supplementary Figure 14: Supplemental to Figure 4.** Representative flow cytometry results of competition experiments in Figure 4E and 4F.

**A)** Representative flow cytometry plots from competition experiments shown in Figure 4E.

**B)** Representative flow cytometry plots from competition experiments shown in Figure 4F.

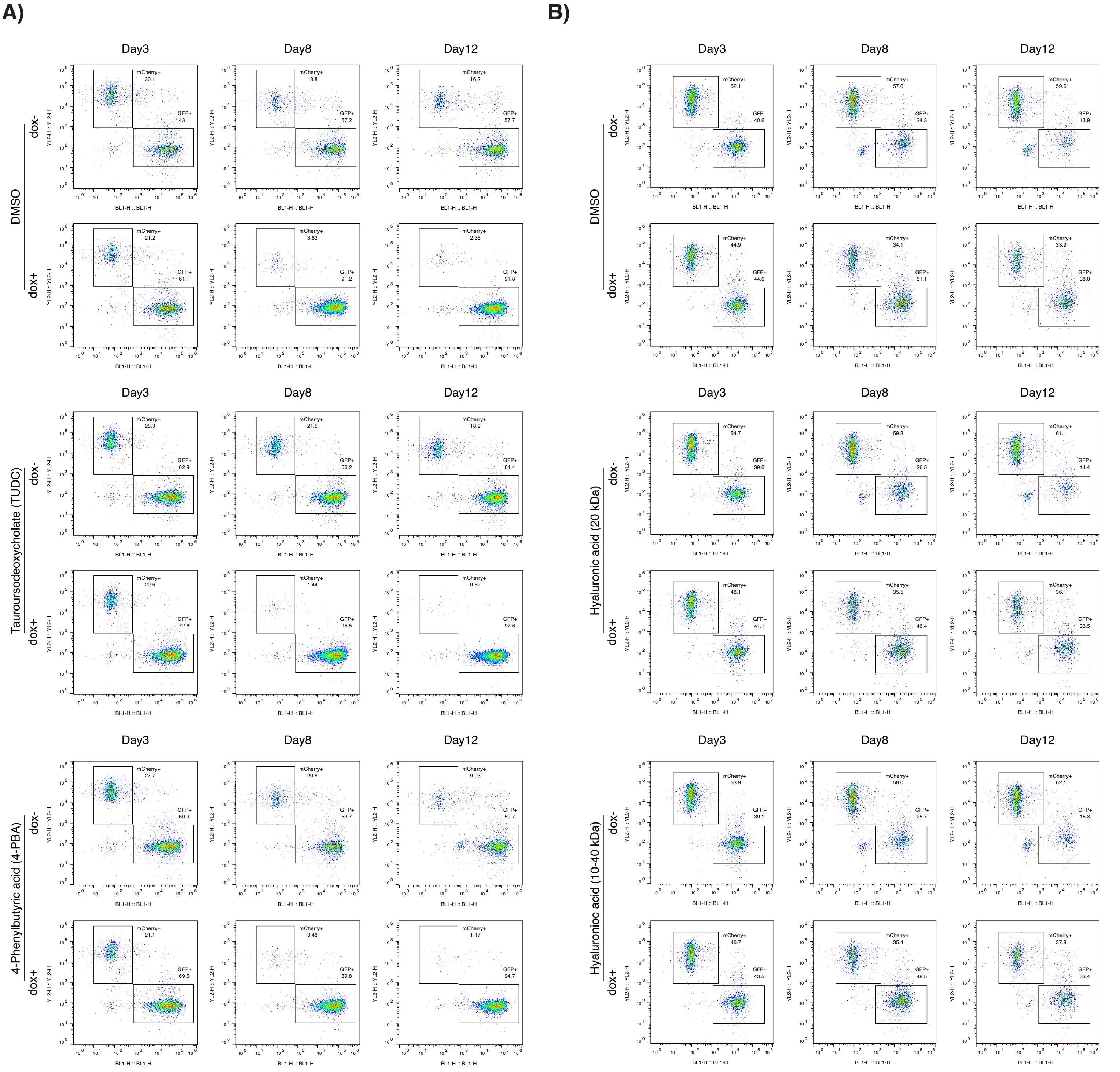


**Supplementary Figure 15: Supplemental to Figure 4.** Representative flow cytometry results of competition experiments in Figure 4H and S12E.

**A)** Representative flow cytometry plots from competition experiments shown in Figure 4H.

**B)** Representative flow cytometry plots from competition experiments shown in Figure S11E.

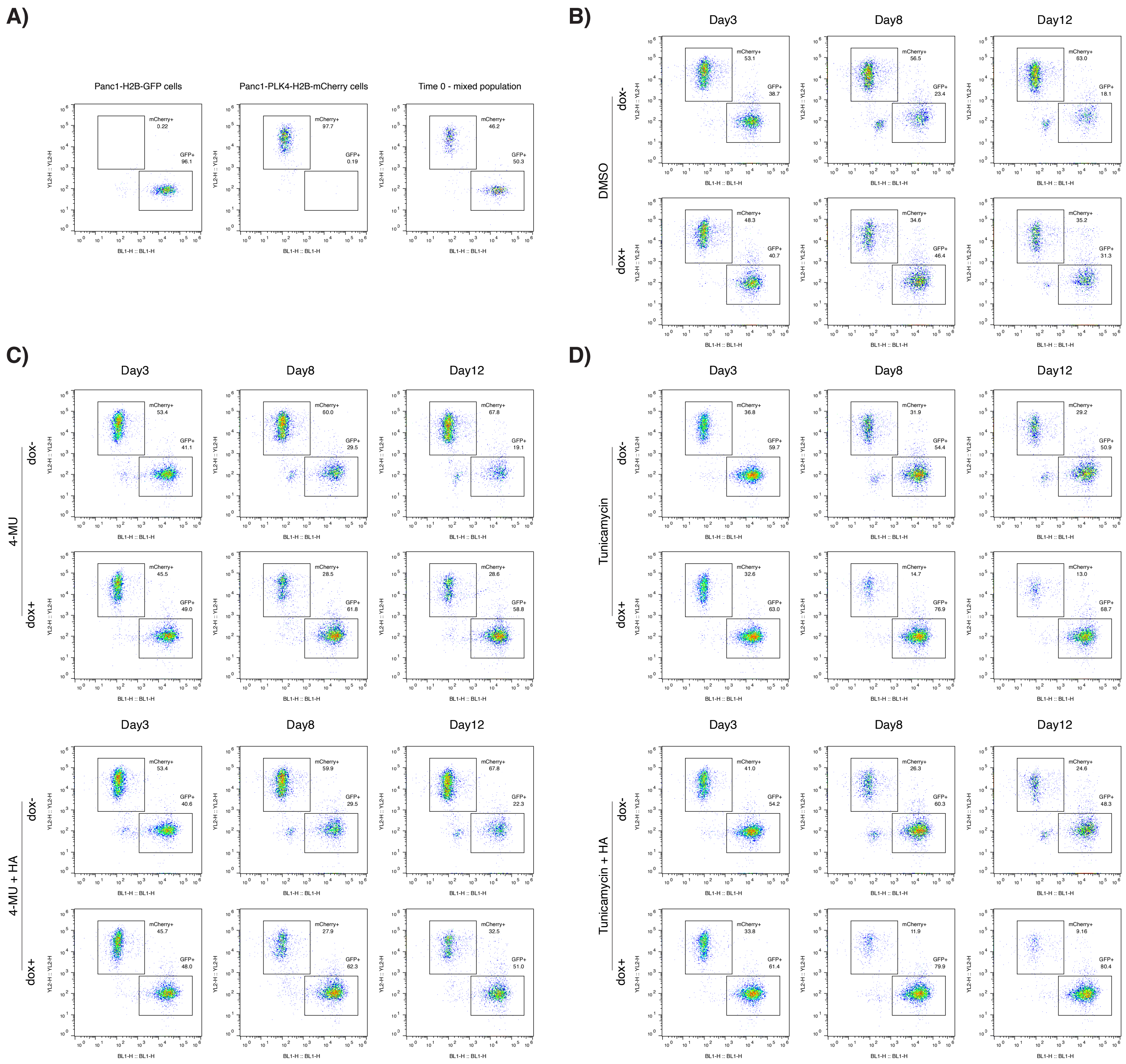


**Supplementary Figure 16: Supplemental to Supplementary Figure 12.** Representative flow cytometry results of competition experiments in Figure S12F.
**A)** Representative GFP and mCherry fluorescence profiles of Panc1-PLK4-H2B-mCherry, Panc1-H2B-GFP, and mixed populations.
**B)** Representative flow cytometry plots of DMSO treated mixed cell populations.
**C)** Representative flow cytometry plots of 4-MU and 4-MU+HA treated mixed cell populations.
**D)** Representative flow cytometry plots of Tunicamycin and Tunicamycin+HA treated mixed cell populations.

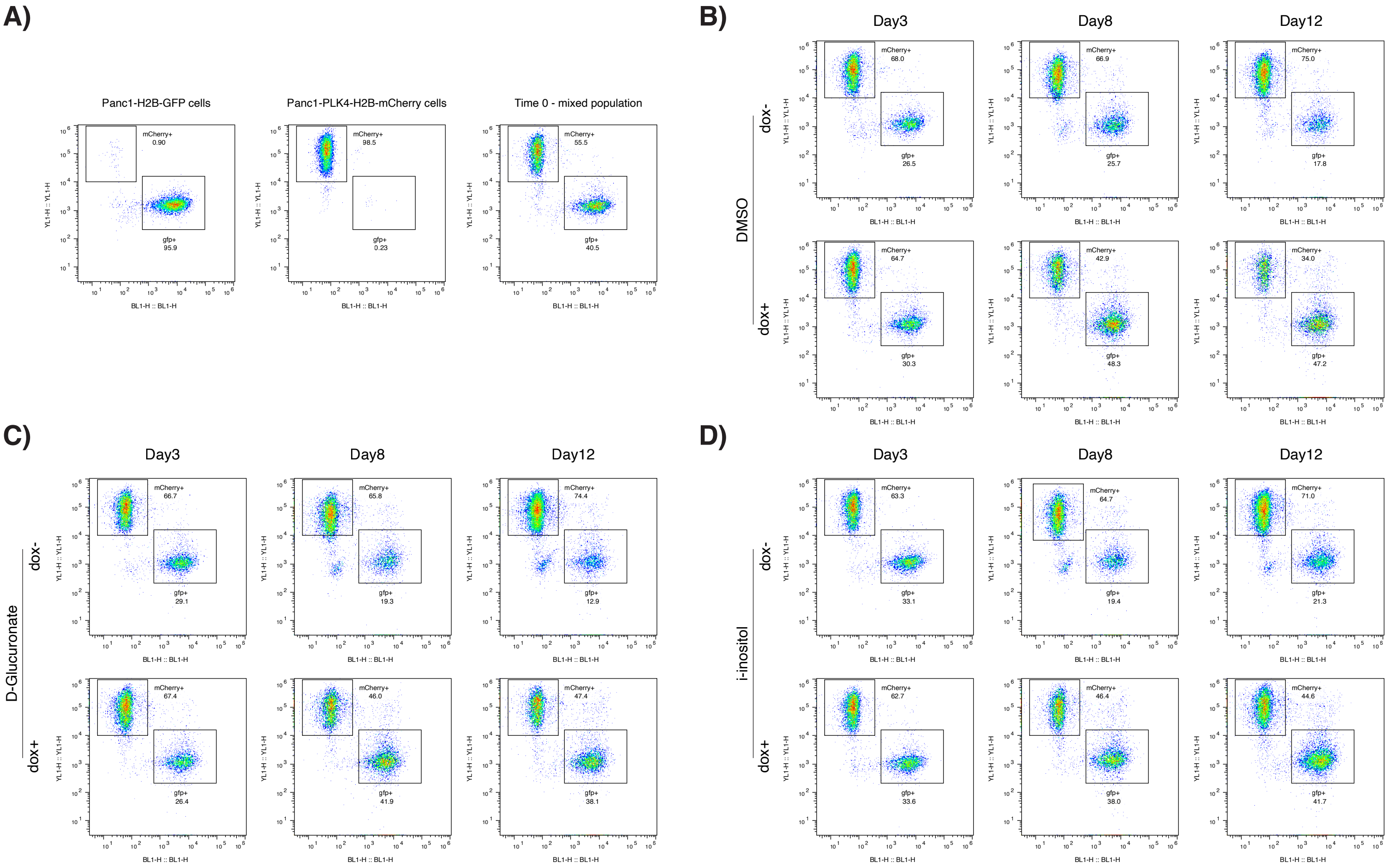


**Supplementary Figure 17: Supplemental to Supplementary Figure 12.** Representative flow cytometry results of competition experiments in Figure S12J.
**A)** Representative GFP and mCherry fluorescence profiles of Panc1-PLK4-H2B-mCherry, Panc1-H2B-GFP, and mixed populations.
**B)** Representative flow cytometry plots of DMSO treated mixed cell populations.
**C)** Representative flow cytometry plots of D-Glucuronate supplemented mixed cell populations.
**D)** Representative flow cytometry plots of i-inositol supplemented mixed cell populations.

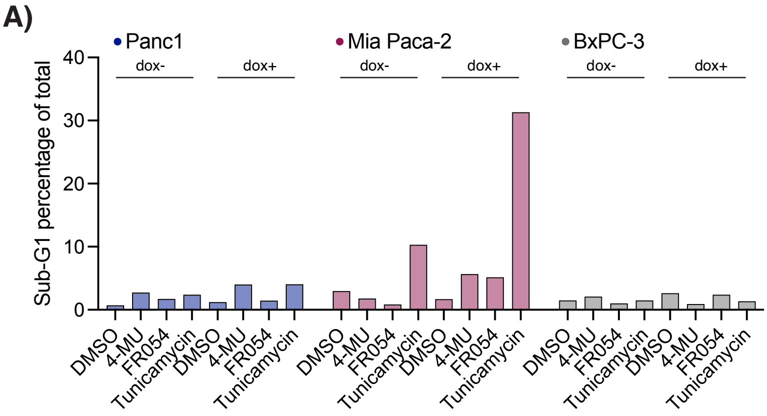
A)
Panc1
Mia Paca-2
BxPC-3
dox-
dox+
Sub-G1 percentage of total
DMSO
4-MU
FR054
Tunicamycin

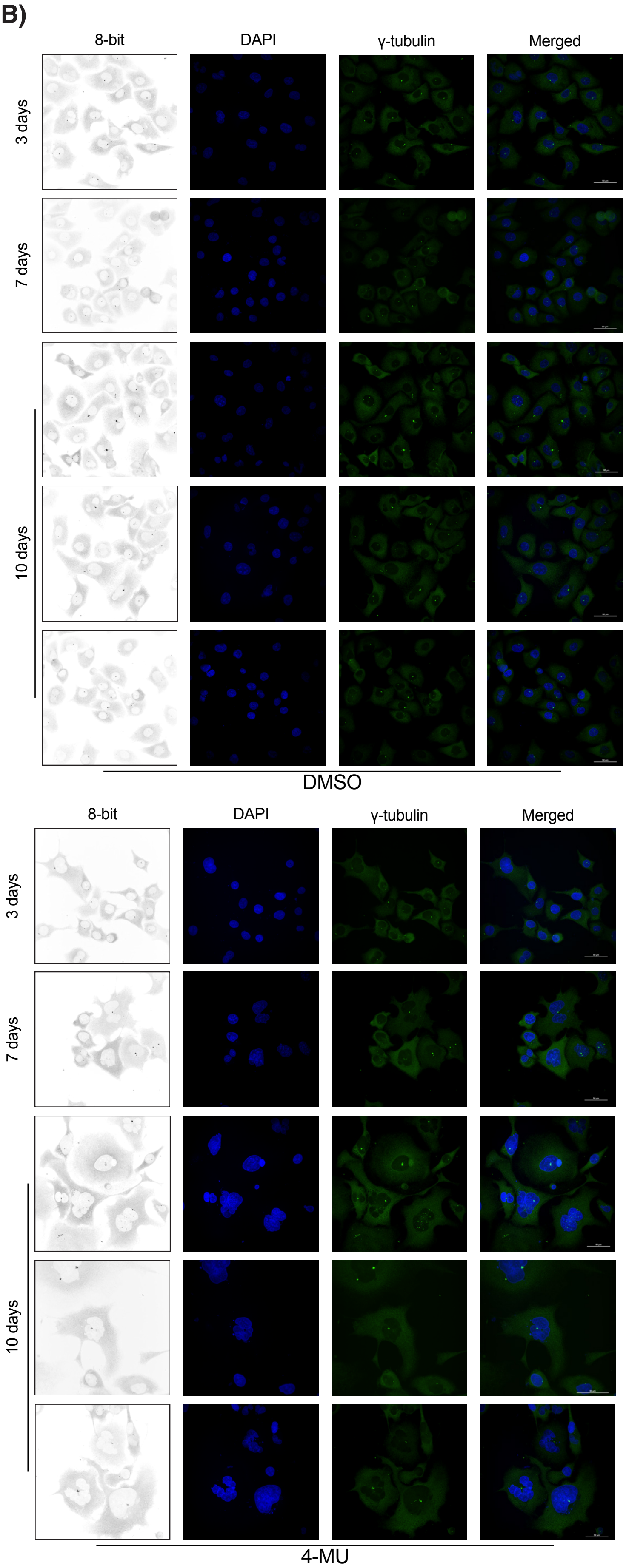
B)
8-bit
DAPI
γ-tubulin
Merged
3 days
7 days
10 days
DMSO
4-MU

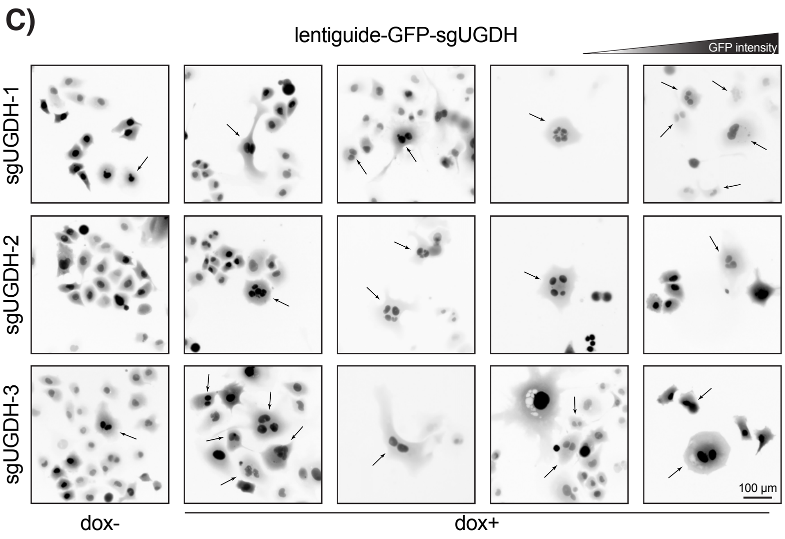
C)
lentiguide-GFP-sgUGDH
GFP intensity
sgUGDH-1
sgUGDH-2
sgUGDH-3
100 µm
dox-
dox+

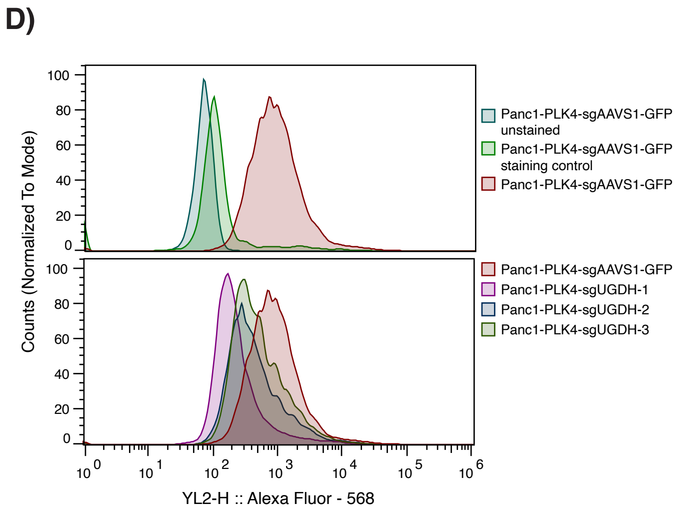
D)
Counts (Normalized To Mode)
Panc1-PLK4-sgAAVS1-GFP unstained
Panc1-PLK4-sgAAVS1-GFP staining control
Panc1-PLK4-sgAAVS1-GFP
Panc1-PLK4-sgUGDH-1
Panc1-PLK4-sgUGDH-2
Panc1-PLK4-sgUGDH-3
YL2-H :: Alexa Fluor - 568

Fig S18

**Supplementary Figure 18: Supplemental to Figure 5.**
**A)** Percentage of sub-G1 cell populations identified in flow cytometry experiment in Figure 5A.
**B)** Confocal images of DMSO or 4-MU treated centrosome amplified Panc1 cells. Blue:DAPI, DNA; Green: γ-tubulin, centrosomes. Inverted images were used in Figure 5E.
**C)** Representative images of lentiguide-GFP-UGDH expressing cells. Multinucleated cells were marked with arrows in the images.
**D)** Surface HA levels in sgAAVS1 (control) and sgUGDH expressing cells. Top panel shows control stainings and bottom panel shows experiment groups. Same flow cytometry result from Panc1-PLK4-sgAAVS1 control cells were used in both panels to make the comparisons easier.

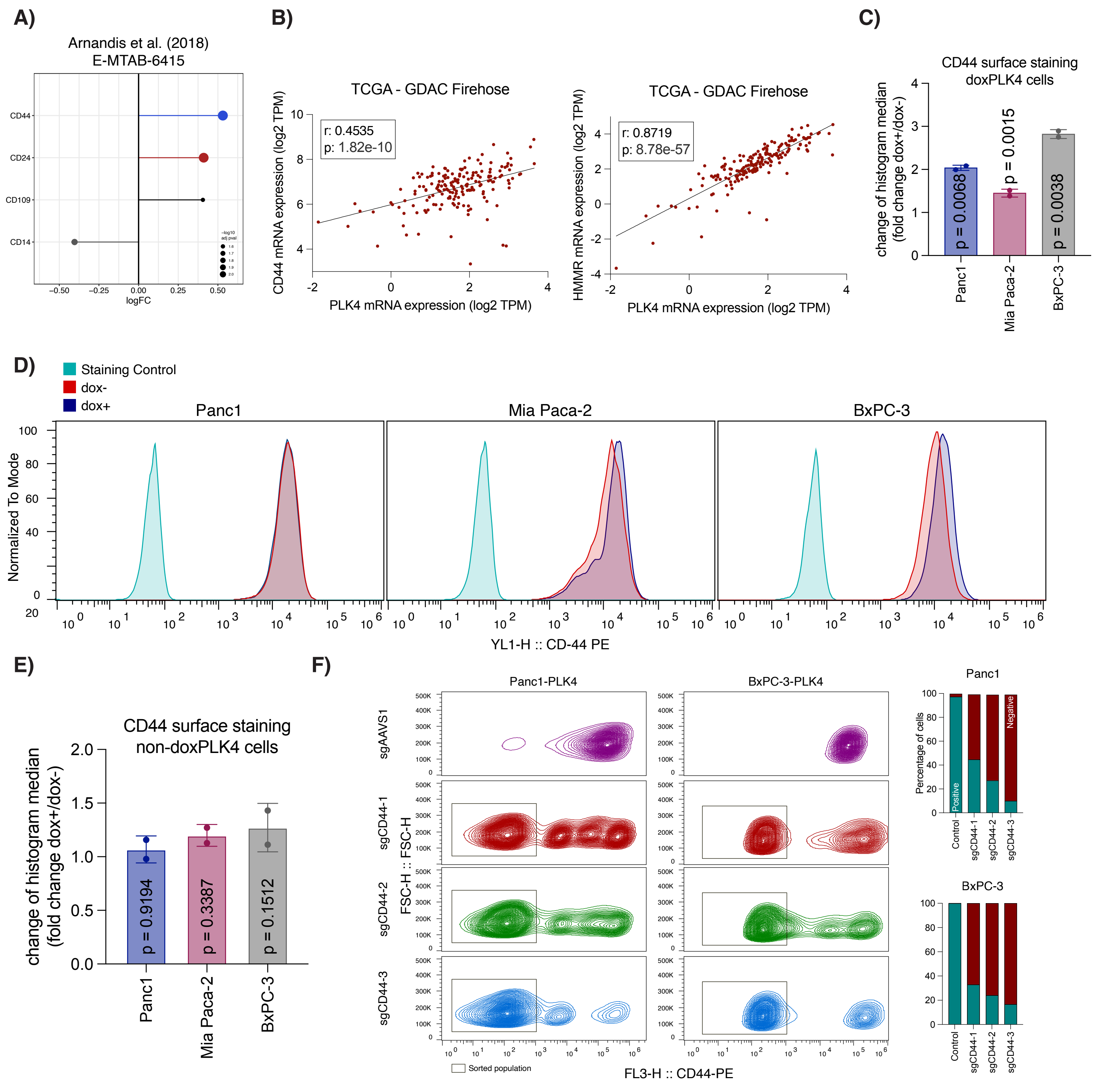


**Supplementary Figure 19: Supplemental to Figure 6.**

**A)** Analysis of microarray data from Arnandis et al. (2018) reveals CD44 upregulation in cells with centrosome amplification.

**B)** Correlation of PLK4 expression with CD44 in TCGA PDAC data.

**C)** Quantification of CD44 surface staining results. Histogram median of dox+ cells were compared to dox- cells and presented as fold change. p values were calculated by two-way t-test and presented on plot.

**D)** Surface CD44 levels does not change upon dox treatment in cells does not carry a doxPLK4 construct.

**E)** Quantification of results in panel C. p values were calculated by two-way t-test and presented on plot.

**F)** Left panel: CD44 surface staining in CD44 targeted sgRNA transduced cells. Sorted populations were marked with rectangulars. Right panel: Quantification of sorted populations (cells that lose CD44 expression after sgCD44)

Fig S19

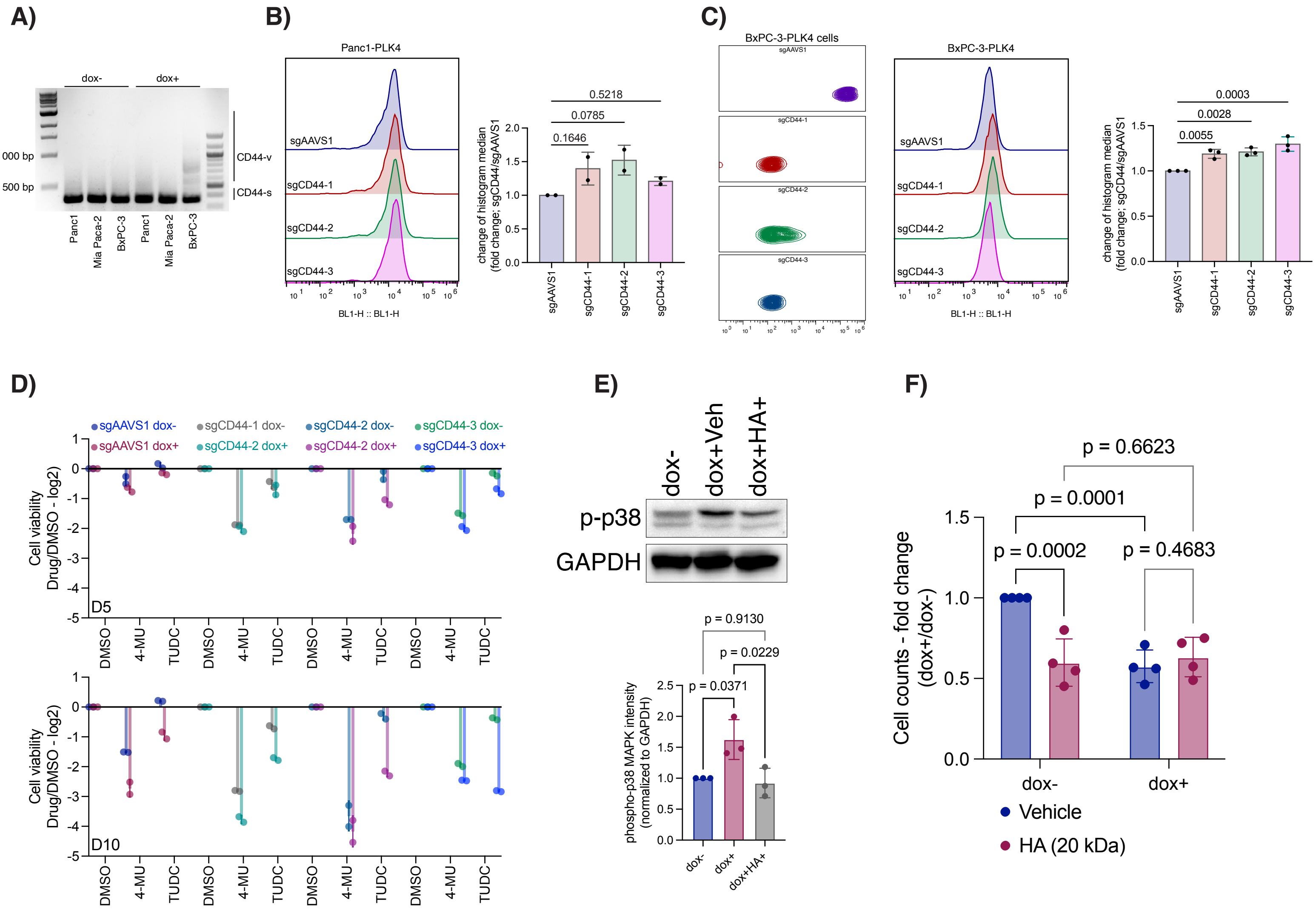


**Supplementary Figure 20: Supplemental to Figure 6.**

**A)** CD44 splicing changes upon centrosome amplification in PDAC cell lines.

**B)** CD44 knockout does not affect intracellular ROS levels in Panc1-PLK4 cells. Left panel: representative ROS measurements. Right panel: quantification of fold changes.

**C)** CD44 knockout slightly increases the intracellular ROS levels in BxPC-3-PLK4 cells. Left panel: sorted cell populations. Middle panel: representative ROS measurements. Right panel: quantification of fold changes.

**D)** TUDC treatment reduces cell viability more strongly in centrosome-amplified CD44-KO cells compared to sgAAVS1 controls.

**E)** Centrosome amplification increases phosphorylation of p38, which is attenuated by HA treatment. Top panel: Western blot results. Bottom panel: Quantification of western blot results.

**F)** HA treatment decreases the viability of dox– control cells but does not affect dox+ cells.

One way ANOVA test was used to determine the statistical significances in panels B, C and E. Two-way ANOVA was used in panel F. p values were presented on plots. Dots represent independent repeats.

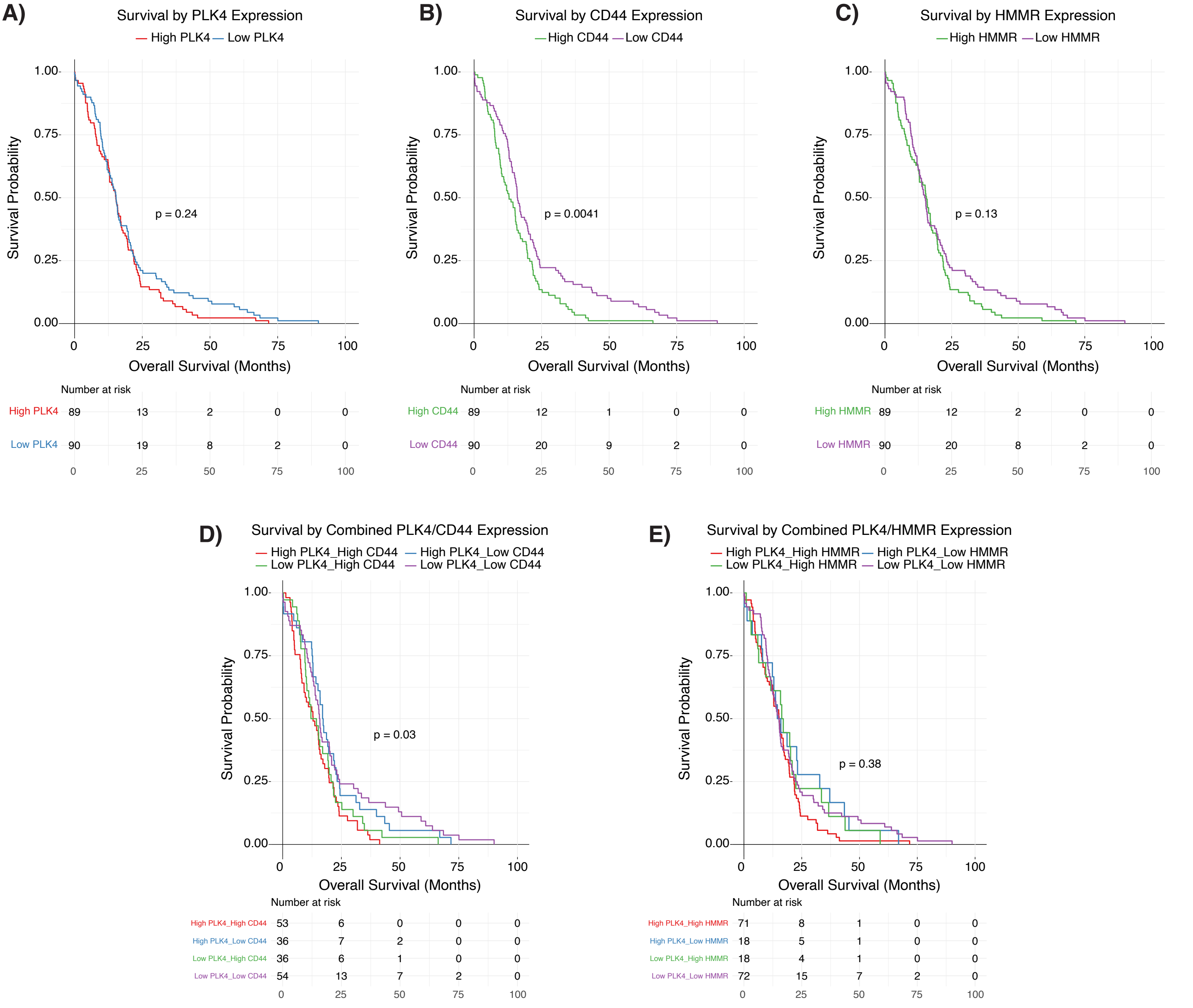


**Supplementary Figure 21: Supplemental to Figure 6.**
Kaplan–Meier survival analyses of pancreatic cancer patients stratified by gene expression levels.
**A)** Survival probability of patients with high versus low PLK4 expression.
**B)** Survival probability of patients with high versus low CD44 expression.
**C)** Survival probability of patients with high versus low HMMR expression.
**D)** Survival probability of patients stratified by combined PLK4 and CD44 expression levels.
**E)** Survival probability of patients stratified by combined PLK4 and HMMR expression levels.

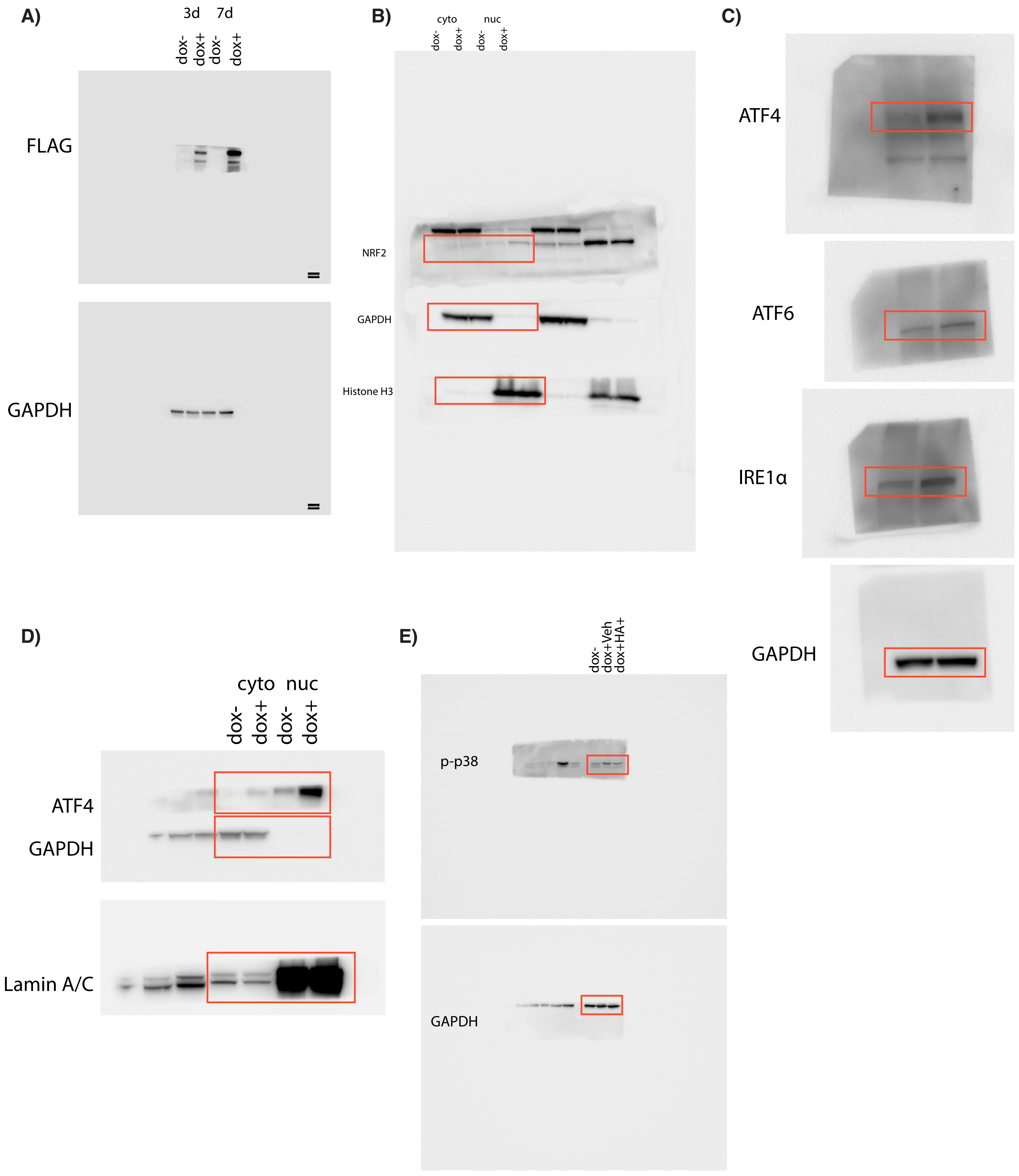


**Supplementary un-cropped Western blots.** The uncropped western blot images corresponding to the main and supplementary figures are shown here. Red rectangles indicate the sections (samples) used in the manuscript figures. Other lanes represent samples from different experiments that were run on the same gel, and were not associated with the current manuscript.

**A)** Uncropped western blot results of Figure 1A.
**B)** Uncropped western blot results of Figure 2E.
**C)** Uncropped western blot results of Figure 4D.
**D)** Uncropped western blot results of Figure S12C.
**E)** Uncropped western blot results of Figure S20E.
